# Truthful visualizations for mass spectrometry imaging enable high spatial resolution interactive *m/z* mapping and exploration

**DOI:** 10.1101/2025.03.18.643852

**Authors:** Jacob Gildenblat, Jens Pahnke

## Abstract

Mass spectrometry imaging (MSI) produces high-dimensional molecular data, but practical interpretation remains limited by visualizations that incompletely preserve global structure. We present MSI-VISUAL, an open-source framework for interactive MSI analysis that integrates truthful dimensionality-reduction visualizations with region-of-interest selection, statistical comparison, and direct *m/z* mapping. MSI-VISUAL introduces four visualization strategies: SALO and SPEAR, optimization-based methods designed to improve global structure preservation across distance metrics; and TOP3 and PR3D, lightweight approaches for memory-efficient, rapid visualization of large datasets. Across benchmarks and lipidomics case studies, including mouse brain and kidney pathology examples, the proposed methods outperform commonly used alternatives in our benchmarks, improve detection of subtle tissue differences, and reveal fine molecular-anatomical patterns that support new biological insights. These results establish MSI-VISUAL as a scalable framework for discovery-oriented and diagnostic MSI workflows.

**Teaser:** Truthful MSI maps reveal hidden tissue patterns, making complex molecular images easier to explore and understand.

## Introduction

Mass spectrometry imaging (MSI) is a powerful analytical approach for spatially resolved molecular profiling of metabolites, lipids, and proteins, with growing importance in experimental biology and diagnostics (*1, 2, 3, 4, 5*). MSI datasets are intrinsically high-dimensional (**rows** × **cols** × **N**), where each pixel contains a full *m/z* spectrum with thousands of molecular features. To make such data interpretable, these spectra must be mapped into low-dimensional visual representations that retain biologically and anatomically meaningful structure. In practice, the central challenge is not only dimensionality reduction, but the preservation of biologically meaningful, interpretable contrast between tissue regions.

MSI data are commonly stored in standardized formats such as imzML (*6*) and are supported by a growing ecosystem of analysis tools and repositories (*7, 8, 9*). Established platforms such as SCiLS Lab provide practical MSI visualization support, including segmentation workflows, and recent versions also include UMAP-based visualizations. However, robust MSI-focused visualization workflows for routine experimental and especially for diagnostic interpretation remain limited.

Most existing MSI visualization pipelines rely on clustering or dimensionality reduction (DR) approaches (*10, 11, 12, 13, 14, 15*). Clustering methods are practical and computationally efficient, but they assign a single discrete label to each pixel, producing hard boundaries and masking subtle intra-region variation. This often obscures fine anatomical detail and reduces sensitivity to biologically meaningful heterogeneity needed for diagnostic workflows in pathology. In contrast, DR methods provide continuous representations (including RGB mappings) that can reveal gradients and substructures not captured by discrete labels.

However, DR quality in MSI depends critically on objective design and distance behavior. In diagnostic settings, visualization quality directly affects the ability to delineate tissue boundaries, detect subtle pathological changes, and guide region-of-interest analysis. A central challenge is preserving both major regional differences and subtle molecular differences within and between neighboring subregions. Many relevant biological differences arise either from coordinated small changes across many *m/z* values or from large shifts in a small subset of ions. Subtle intra-ROI differences are preserved only when pixels with slightly different spectra maintain their ranked distance relationships after projection; if these relationships collapse, distinct pixels appear too similar, global-structure fidelity decreases, and diagnostically relevant detail is lost. This motivates explicit optimization of global structure preservation and the use of complementary distance functions to capture both distributed molecular changes and hot-spot-driven variation. This is particularly problematic in structures with partially shared molecular composition, such as metabolically distinct tumor clones, infiltrative tumor margins blending into adjacent parenchyma, metastasis borders, small mixed protein-lipid deposits in the brain, neighboring thalamic or hippocampal subregions, and muscle fibers with focal metabolite accumulation.

A wide range of general-purpose DR methods have been applied to MSI, including PCA, t-SNE, UMAP, and related approaches (*16, 17, 18*). While effective in many settings, these methods are not designed specifically for MSI and do not jointly address key practical requirements: (i) preservation of global molecular-spatial relationships, (ii) robustness across complementary distance behaviors, and (iii) scalability to large datasets with thousands of features per pixel.

In addition to visualization, practical MSI analysis requires fast, hypothesis-driven exploration. Users must select regions of interest (ROIs), compare molecular profiles between regions, and verify *m/z* specificity directly in image space. Existing workflows often separate these steps across tools, slowing analysis and reducing reproducibility in iterative exploratory settings. Previous work mapped region-associated *m/z* values by registering MSI to MRI (*19*) or using MALDI-IHC (*20*), an effective but added cross-modality step.

Here, we present an open-source framework for truthful, interactive MSI visualization and exploration. We introduce SALO and SPEAR, two optimization-based DR methods that explicitly target structure preservation, and TOP3 and PR3D, lightweight approaches for rapid visualization of large datasets. MSI-VISUAL integrates ROI-based *m/z* mapping, statistical comparison, SpecQR summaries, and Virtual Pathology Stains (VPS) for efficient interactive analysis. Across controlled mouse brain data and diverse METASPACE datasets, MSI-VISUAL improves interpretability, enables detection of fine-grained molecular-anatomical patterns, and supports scalable experimental and diagnostic workflows. Importantly, it supports pathologist-guided MSI-native analysis without requiring registration to other modalities, such as MRI or IHC.

## Results

### Overview of key improvements

MSI-VISUAL delivers a substantial advancement over current MSI visualization workflows. MSI-VISUAL is designed to provide complementary visualizations rather than replace existing methods, because different views reveal different spatial-molecular details. Quantitatively, SALO and SPEAR show improved global-structure preservation and strong benchmark performance across datasets (see Section Quantitative improvements). Qualitatively, the proposed visualizations reveal biologically and clinically relevant fine detail, including subtle intra- and inter-subregion differences beyond major regional contrasts, that is frequently missed by both continuous embeddings (e.g., UMAP) and hard-partitioning approaches (e.g., K-means segmentation) (see Section Qualitative improvements). At scale, TOP3 and PR3D enable fast, memory-efficient exploration of large MSI images (see Section Scalability improvements). Finally, we demonstrate an MSI-native workflow that combines ROI selection, comparison, and ROI-to-*m/z* mapping in one interface, without requiring registration to other modalities, such as MRI or IHC, and we showcase this across multiple translational examples (see Section Translational utility of MSI-VISUAL). Together, these advances establish MSI-VISUAL as a high-fidelity, scalable, and practical framework for diagnostic- and discovery-oriented MSI analysis.

### Quantitative improvements: improved structure preservation across benchmarks

We asked whether MSI-specific objectives can improve global-structure fidelity while remaining robust to complementary distance behaviors. We therefore developed SALO and SPEAR:

#### High global structure preservation for revealing anatomical details in sub-regions

This is achieved through direct optimization of global structure preservation. These methods sample representative reference points in the image and optimize an objective based on the ranked distances of all image pixels from those reference points in both the high-dimensional data and the visualization. Both SPEAR and SALO rely on a differentiable approximation of the ranking operation (*21*). SPEAR optimizes the Spearman rank correlation (*22*). SALO optimizes a new objective, termed *Saliency*, which requires ranks of point-pair distances in the visualization (in LAB color space to improve perceptual contrast) to be at least as high as in the MSI data, thereby preserving detail. Compared with PCA, our methods are non-linear and non-parametric, and can model complex relationships that are not captured by linear projection. Compared with UMAP (*18*), which emphasizes local neighborhoods and local structure preservation, our methods explicitly target global structure preservation. PaCMAP (*23*) also samples distant point pairs, which is conceptually related to our reference point strategy. However, unlike PaCMAP, we directly optimize global structure by maximizing distance correlations and jointly considering relationships among multiple points. On the mouse brain dataset, this yields higher global structure preservation (Figure S2); SALO achieves an average correlation of approximately 0.8, compared with approximately 0.45 for PaCMAP. Because we avoid graph structures used by UMAP and PaCMAP, our methods have lower memory footprints and better scalability. On larger MS images (*>*50 GB), UMAP did not complete even on high-memory hardware (Figure S4).

#### Robustness to different distance metrics

Euclidean (and similarly cosine) distances are sensitive to coordinated changes across many *m/z* values, whereas the *L*_∞_ distance emphasizes large changes in a single *m/z* value. SALO and SPEAR achieve robustness to both behaviors by fusing Euclidean and *L*_∞_ information through the maximum of their ranks. By jointly preserving global structure and complementary distance behavior, SALO and SPEAR improve separation of subtle intra-ROI variation while retaining major inter-region organization. In parallel, TOP3 and PR3D use continuous, intensity-ordered RGB mappings that avoid hard cluster boundaries, preserving gradients and local molecular deviations while enabling fast, memory-efficient exploration at scale.

#### Benchmark performance across datasets

To quantitatively evaluate performance, we benchmarked the proposed visualizations on two datasets (Tables S1, S2) using established MSI metrics (*24, 25, 26*).

We benchmarked against modern methods (UMAP (*18*), PaCMAP (*23*), Phate (*27*), t-SNE (*17*), TriMap (*28*)) as well as classic methods (PCA (*16*), NMF (*29*), Isomap (*30*), and FastICA (*31*)). Averaged results across images are presented in Figure S2. SALO performs best across datasets and individual images, including versus SPEAR (Figure S3). SALO and SPEAR perform best among the tested methods on both datasets. TOP3 is the strongest existing method for the *L*_∞_ distance and consistently outperforms UMAP while using far less memory and runtime on large images (Figure S4). PR3D is competitive but less robust on the more diverse METASPACE dataset, likely because these images vary substantially in molecular content and effective binning characteristics, while PR3D currently uses fixed percentile settings. This sensitivity is detailed in the supplementary PR3D percentile panel (Figures fig:PR3D-percentiles-brain and S23), where different fixed percentile settings produce markedly different recovered structures. Automatic percentile tuning should therefore improve robustness and reveal additional detail across heterogeneous datasets.

### Qualitative improvements: improved biological and clinical detail recovery

To assess the practical improvements of our new visualizations compared with currently used methods, we analyzed examples containing different known histomorphological details. For direct comparison against both continuous embeddings and hard-partitioning approaches, see UMAP in the main figures and K-means segmentations in Figures S1 and S19.

#### Detection of pathological changes of diagnostic relevance

i) Parenchymal organs have a well-defined histomorphological architecture with several functional substructures and subregions. The kidney consists of many functional units, each comprising a glomerulus in the cortex and a connected tubular system extending from the kidney cortex to the medulla. This is not only true for proteins, but also for lipids. Figure 2 shows two visualizations highlighting these anatomical details in a glomerulonephritis case using lipidomics MSI at 5 µm pixel resolution (see also Figure S7). SALO clearly highlights different parts of the tubular system (proximal and distal tubuli, collecting duct), as well as numerous inflammatory cells between the tubuli and surrounding the glomeruli in their typical pattern. The number of glomeruli is reduced, and the few remaining glomeruli are strongly pathologically changed. SALO also enables clear detection of remaining glomerular capillaries and single inflammatory cells.
ii) Alzheimer’s disease-related A*β* plaques are readily detectable with H&E staining by trained observers (Figure S16) or immunohistochemical stains (Figure S17) and appear at different sizes and locations in mouse models (*32, 33*). The plaques are circumscribed, highly pathological proteinaceous structures that also accumulate several specific lipids (*34*). Their peptide and lipid composition changes over time (*35, 36, 37, 38, 39*). Figure 1 demonstrates improved detection of these histopathological changes using the new visualizations (white open arrows). The assessment is based solely on a lipidomics MSI scan of a mouse brain hemisphere (bregma -2.06 mm, pixel resolution 20 µm) from a 100-day-old APP-transgene mouse with numerous small and large A*β* plaques. Whereas the previous visualizations primarily detect coarse regions, the proposed new visualizations (SALO, TOP3, and PR3D) highlight numerous contrasted deposits, including small A*β* plaques. K-means segmentation misses these plaques entirely and does not assign a specific category (see also Figure S19).

#### Detection of functional microanatomy in hippocampus

We sought to assess whether the proposed visualizations can reveal subtle functional details in mouse brain tissue, such as hippocampal interconnections between neuronal groups. In contrast to circumscribed pathological A*β* plaques, these physiological structures comprise axonal and dendritic connections, neuronal cell bodies, and intercalated glia, and may appear less pronounced because nearby regions can share similar *m/z* content due to close spatial organization. Figures S8, S9, and S10 present overviews and zoomed-in UMAP, SALO, SPEAR, PR3D, and TOP3 visualizations, showing different levels of hippocampal anatomical detail and global-structure preservation compared with conventional H&E and immunohistochemical (IHC) stains. Such visualization panels can support the discovery of functional subregions beyond current atlas annotations and reveal subanatomical structures relevant to disease research and treatment evaluation.

**Figure 1:**
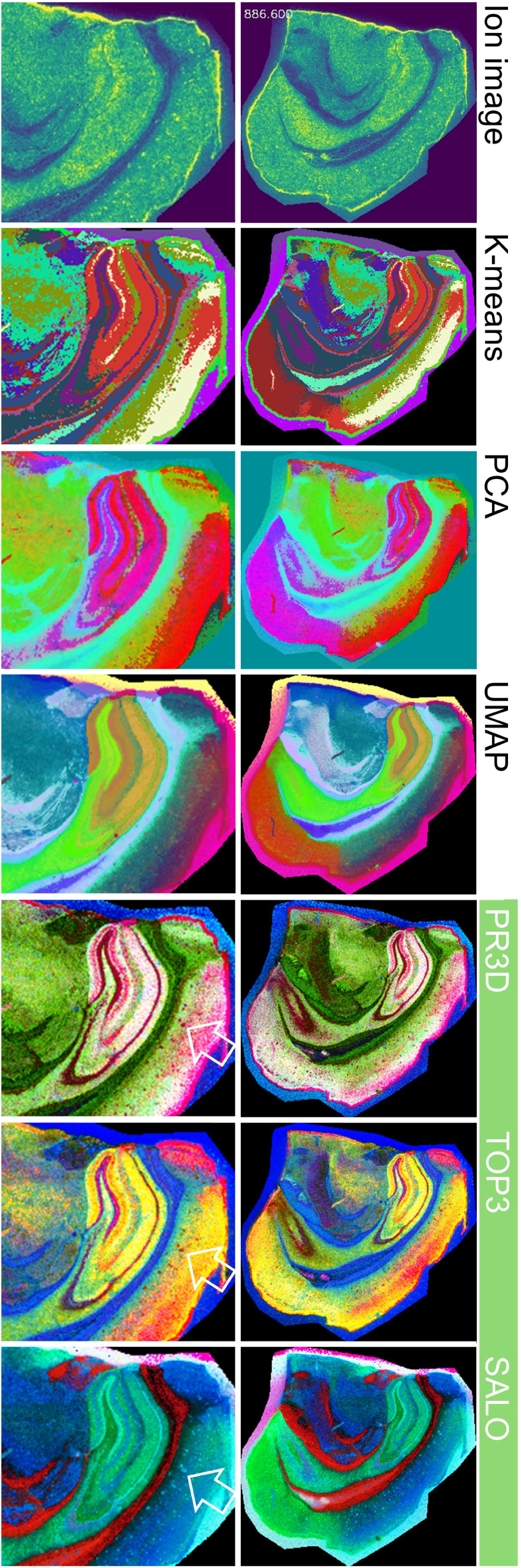
**Improved visualizations** for a 20 *μ*m/pixel MSI scan of a mouse brain with Alzheimer’s disease and *β*-amyloid plaques. Existing methods (left) fail to reveal most plaques and show limited intra-region detail. Ion images at specific *m/z* values indicate plaque distribution but require prior knowledge or manual search. K-means and PCA miss most plaques and capture limited variation, while UMAP reveals more detail but still misses plaques. In contrast, the proposed new methods clearly reveal plaques and finer sub-region structure. PR3D and TOP3 are lightweight and scale to large images and many bins, and SALO has substantially lower memory requirements and runtime than UMAP (Fig. S4). As single views may miss details, MSI-VISUAL presents multiple visualizations, enabling interactive validation via *m/z* mapping (Section Translational utility of MSI-VISUAL).

**Figure 2:**
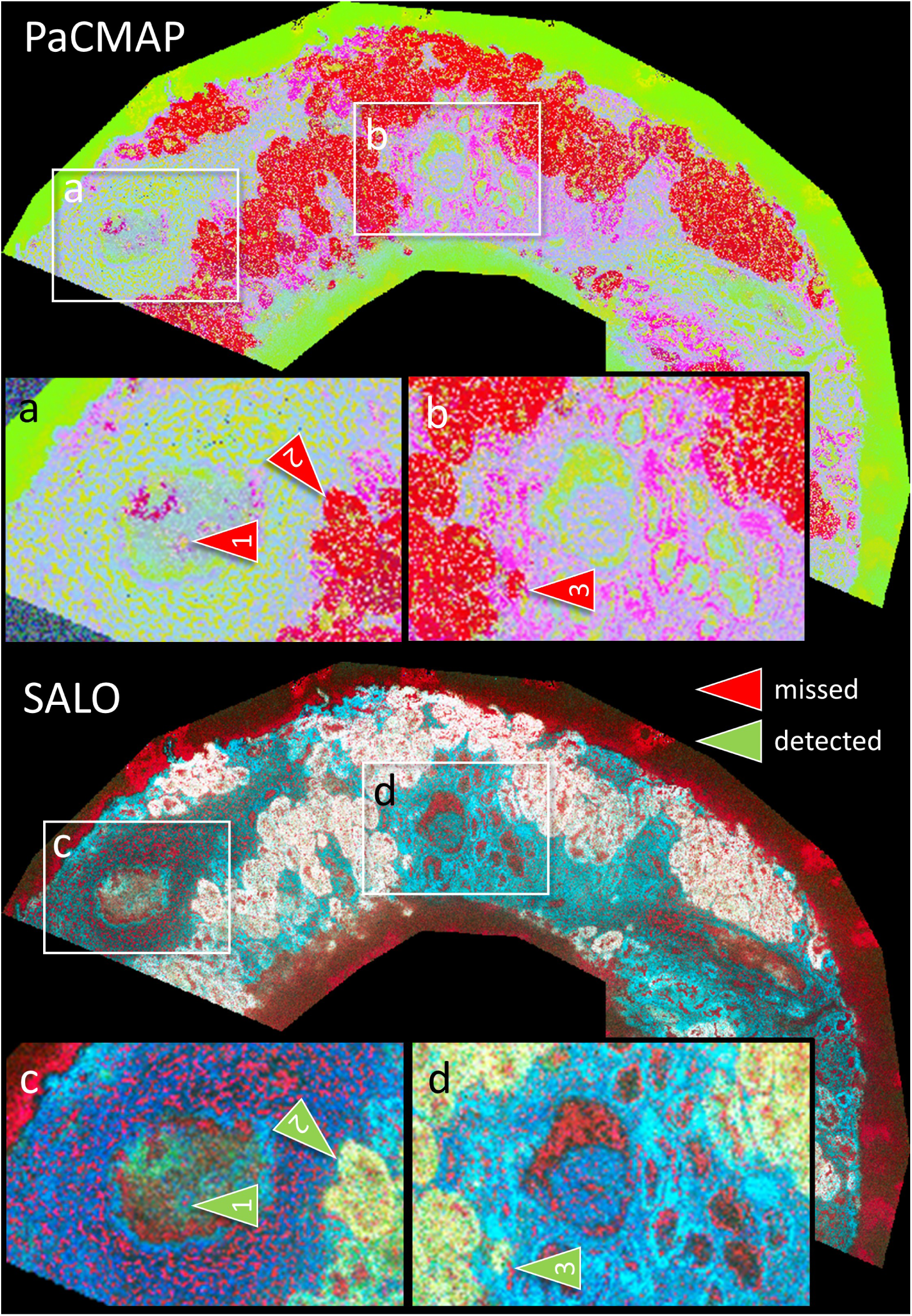
Diagnostic example of a human kidney needle biopsy. Surprisingly, even lipidomics-only data (5 *μm*/pixel) can reveal detailed kidney microanatomy and disease-related features. SALO reveals pathological structures more clearly and better resolves internal organ architecture, including tubule types. (**A** to **D**) show two examples of glomeruli in PaCMAP (a recent method that improves UMAP (*23*)) (**A, B**) and the improved SALO visualizations (**C, D**). The arrows mark features that are more clearly detectable in SALO: (1) inflammatory cells in a glomerular capillary in (**A, C**), (2) proximal versus distal tubules in (**A, C**), and (3) inflammatory cells (lymphocytes) in a tubule in (**B, D**).

#### Detection of substructures in brain nuclei

Detecting subtle anatomical or functional details within known anatomical regions is essential for research on disease pathomechanisms and treatment development. Figure 3a demonstrates how the proposed visualizations highlight subtle detail in thalamic nuclei. Small thalamic nuclei are especially pronounced in SALO and TOP3 visualizations. SALO was used to annotate representative thalamic subnuclei (dotted lines, arrows). The resulting annotation mask was then overlaid on the other visualizations to enable comparison and orientation within the thalamic region. UMAP shows substantially less detail within the thalamic annotation mask. PR3D and TOP3 reveal some, but not all, thalamic nuclei.

**Figure 3:**
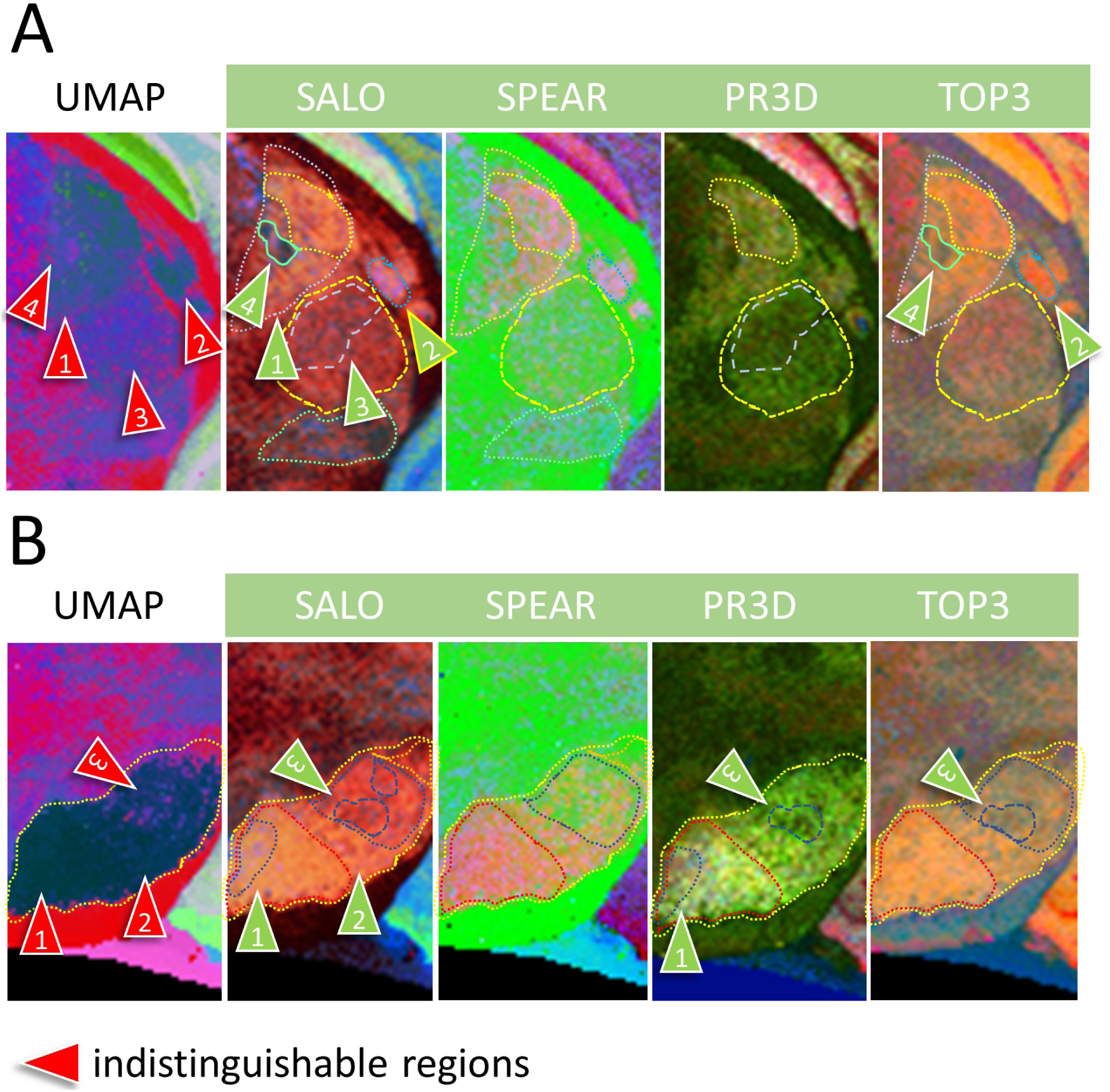
Improved substructures - delineation of thalamic and substantia nigra subregions. (**A**) Improved detection of thalamic nuclei. SALO and TOP3 reveal greater subregional detail within the annotations than UMAP, which shows a largely homogeneous thalamic area. Red arrows indicate thalamic nuclei that are not distinguishable in UMAP but are identifiable and annotatable in SALO. (**B**) Subregions of the substantia nigra (SN) are detectable with the proposed visualizations, whereas UMAP shows a predominantly homogeneous area. Green arrows indicate examples of subregions detectable in SALO, SPEAR, PR3D, and TOP3. Arrow legend: green, detection; red, no detection.

Another example of increased structural detail is shown in Figure 3b, which illustrates subregions (functional domains) of the substantia nigra (SN). Several SN subregions representing functional subdomains are visually detectable with the proposed visualizations. By contrast, with UMAP-based DR, these subregions are not detectable: UMAP shows only a homogeneous dark-blue area, whereas the new visualizations enable clear separation of subregions. For comparison, we include a panel of visualizations and K-means segmentations in Figure S1.

### Scalability improvements: faster, lower-memory visualization for clinical-scale MSI

#### TOP3 and PR3D enable lightweight scalability

We next asked whether lightweight visualizations can support large, otherwise intractable datasets. Because total ion current (TIC) thumbnails offer limited regional discrimination, we developed lightweight multicolor alternatives.

The motivation for these lightweight visualizations is the observation that, in our mouse brain dataset, most image pixels contain a small number of dominant high-intensity *m/z* values (approximately 75% with one dominant *m/z*, 20% with two dominant *m/z* values, and the remaining pixels with three or more). The proposed TOP3 visualization measures the three largest intensities in each pixel and, after outlier removal, converts these values from LAB color space into RGB. The proposed Percentile-Ratio (PR3D) visualization generalizes this approach and measures ratios of fixed, predetermined intensity percentiles. TOP3 and PR3D provide a complementary view by being sensitive to intensity changes in a small number of dominant *m/z* values, enabling visualization of subtle subregional contrast even when most spectral content is shared with surrounding tissue. As shown in Figure S4, both methods are computationally lightweight, have low memory footprints, and are hundreds of times faster to compute. This enables rapid MSI visualization for scans with higher computational demands (more pixels, more *m/z* values, or less binning) that other methods fail to handle.

#### Clinical-scale evaluation

We evaluated scalability on a 5 µm human kidney biopsy (Table S2, dataset 4; 550 × 824 pixels, 5,255 features; 9.3 GB .npy). Despite 320 GB RAM, several methods - including UMAP - did not complete because of memory demand (Figure S4). TOP3 was over 600× faster than PaCMAP and required only 2 MB additional memory, versus approximately 17.5 GB for PaCMAP/SALO. SALO memory depends on reference points (default **N** = 1,000); reducing **N** trades memory for detail. Overall, TOP3 is the most practical option for rapid exploration of very large MSI scans, whereas SALO provides higher-fidelity structure preservation.

### Translational utility of MSI-VISUAL: integrated interactive ROI-to-*m/z* mapping for interpretation and diagnosis

Determining the spatial distribution of region-specific *m/z* values (mapping) is central for both biological discovery and diagnostics. Here, mapping is performed in an MSI-only workflow, without requiring registration to other modalities, such as MRI or IHC. In future, disease-associated *m/z* signatures from MSI analyses may support treatment stratification to target metabolic pathways.

To assess generalizability across *m/z* ranges and spatial resolutions, we analyzed (mapped) five representative cases:

i) Fast generation of region-specific lipid m/z signatures.
ii) Mapping lipidomics and metabolomics examples from a public database.
iii) Mapping human diagnostic brain samples with neurodegenerative diseases.
iv) Mapping mouse brainstem nuclei and white matter tracts.
v) Mapping single neurons of the spinal cord.

#### Fast generation of region-specific lipid m/z signatures

We demonstrate rapid interactive *m/z* mapping across cortex, hippocampus, white matter, and brain stem in a lipid MSI scan from an AD mouse model with pronounced *β*-amyloidosis (*33, 36*). Figure 4 shows region-specific SpecQR profiles and top-five *m/z* values with direct ion-image verification, without MRI/IHC registration required in earlier workflows (*19*).

**Figure 4:**
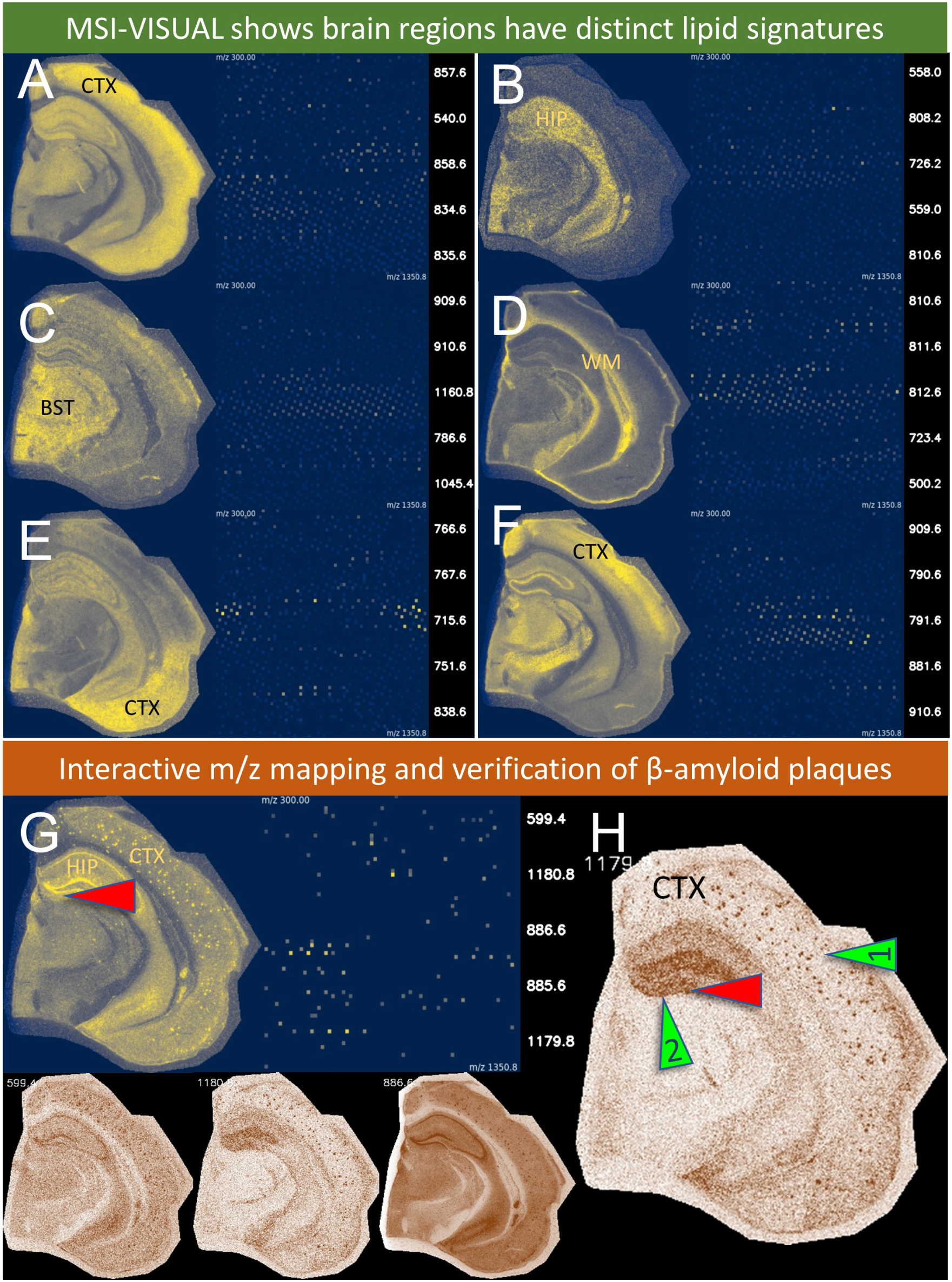
Interactive brain-region *m/z* mapping in an AD mouse model. (**A** to **H**) brain lipid MSI scan of an AD mouse model with A*β* plaques. (**A** to **F**) ion current summary images and SpecQR graphs selective for (**A**) the isocortex (CTX), (**B**) the hippocampus (HIP), (**C**) the brain stem (BST), (**D**) white matter (WM), (**E**) piriform cortex (CTX), and (**F**) upper isocortex regions (CTX) and connections in the hippocampus and brain stem. (**G**) A*β* plaques have a distinct lipid *m/z* signature with 599.4, 885.6, 886.6 (M+1), 1179.8, and 1180.8 (M+1) being the 5 highest *m/z* values in this example. The SpecQR summary image and the Virtual Pathology Stains (VPS) of the top *m/z* values are used for verification (in brown). VPS delivers a continuous ’stain’ similar to those used for pathology diagnostics. (**H**) An image with numerous plaques (arrow 1) in the isocortex (CTX), containing the plaque-specific *m/z* 1179.8 (ganglioside GM3 (*37*)), which is also found in hippocampal connections (arrow 2). The red arrow depicts an A*β* plaque also labeled in the consecutive H&E-stained section of the frozen tissue block used for MSI (Figure S16).

Using MSI-VISUAL, we mapped these regions directly without registration to MRI or IHC. Region selection is performed nearly instantly and in a fraction of the time reported previously (*19*). This is relevant for both diagnostics and discovery-oriented MSI workflows.

We map A*β* plaques simply by selecting them in the MSI-VISUAL viewer. Of note, some *m/z* values, such as 885.6 Da, highlight the plaques in addition to cortex regions, showing that the A*β* plaques absorb and concentrate these lipids from the cortex. Other *m/z* values, such as 1179.8 are more unique to cortical plaques and are mostly seen in the hippocampus. To our knowledge, this is the first direct determination of plaque-associated *m/z* values from MSI images, rather than restricting analysis to pre-specified plaque *m/z* targets.

#### Cross-cohort mapping in public metabolomics/lipidomics datasets

For external validation, we analyzed open-access datasets from METASPACE (*9*). Figure S11 shows successful *m/z* mapping in a human kidney biopsy (lipidomics, *m/z* 300–1350, 5 *μm*) and a mouse brain with experimentally implanted tumor cells (metabolomics, *m/z* 50–400, 20 *μm*), including inflammation-associated and tumor-associated signatures.

#### Diagnostic differentiation of mixed neuropathological lesions

We analyzed *post-mortem* human brain tissue from a patient with vessel-related dementia and AD using peptide MSI (*40*). Figure 5 shows that MSI-VISUAL separates AD-related A*β* deposits from vascular-dementia-related capillary leakage. Color-based ROI selection followed by top-five *m/z* extraction correctly localized both processes in this mixed-dementia case.

**Figure 5:**
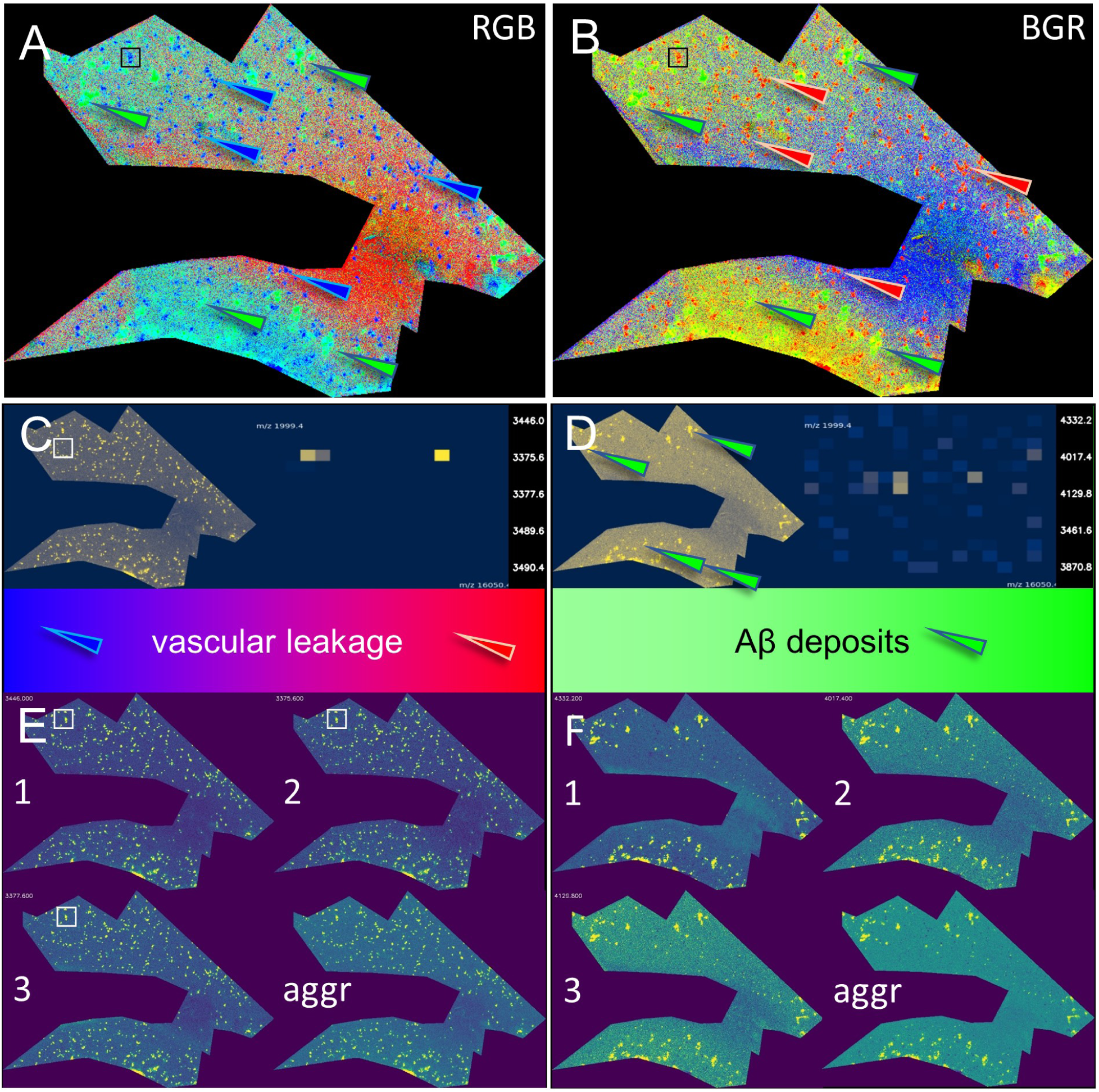
Differential m/z signatures distinguish mixed neuropathological lesions. (**A, B**) NMF visualization (2000 iterations) in 3 color channels (RGB), presented with two colorizations (RGB and BGR). The box and red/blue arrows mark examples of numerous smaller, partially accumulating/aggregating structures. The green arrows highlight larger, more confluent structures. These structures were selected for *m/z* mapping and identification of their potential pathological origin. (**C**) The *m/z* fingerprints and SpecQRs reveal extensive capillary leakage (albumin, ≈3.4 kDa), and (**D**) larger A*β* depositions with A*β*1–40 (4332.2 Da) and A*β*1–38 (4129.8 Da). (**E, F**) presentations of the three most significant *m/z* values (1–3) and the aggregated mean images of these three *m/z* values (aggr). (**E**) highlights severe capillary leakage as seen in vascular dementia. (**F**) shows A*β* pathology as seen in Alzheimer’s disease. Thus, the neuropathological *post mortem* diagnosis is mixed-type dementia. ProteomeXchange dataset PXD049325, peak-selected, peptide MSI, case S2-SAD provided by (*40*).

#### Functional mapping of brain nuclei and white-matter tracts

A additional examples, MSI-VISUAL resolved functional nuclei and white-matter connections (Figures S12 and S13). In Figure S13, TOP3 separates substantia nigra (SN) and cerebral peduncle (cpd), which were used as ROIs for *m/z* comparison. ROI1 (SN) mapped to neuron-rich gray matter, whereas ROI2 (cpd) mapped to white matter, with clear separation in ion images and SpecQR.

#### Mapping single neurons of the spinal cord

We assessed whether MSI-VISUAL can support single-cell detection and analysis. As a favorable example, we used large spinal cord *α* motor neurons (*α*MNs) in anterior-horn gray matter, consistent with prior guided-MSI and related work (*41*). We applied this workflow to MSI sections with 5 *μ*m spatial resolution (Table S2, dataset 5; Figure S14). First, we employed a digital selection step for a ’single cell’ ROI (repeated 3 times) to be able to assess the molecular profile of the *α*MNs vs. the surrounding spinal cord tissue. Each time, the diagram showed signal enrichment in the *α*MNs and partially in the surrounding gray matter of the spinal cord. The selection was guided to be negative in the spinal cord’s white matter (myelinated tracts). The mapped *m/z* values can be used to confirm the cell entity (*42*) and to evaluate changes during disease processes, e.g., in amyotrophic lateral sclerosis (ALS), a neurodegenerative disease involving the first (in the brain’s motor cortex) and second (in the spinal cord) motor neurons.

### Improvement of biological interpretation through mapped *m/z* values

For neurobiological interpretation, we provide two examples showing how mapped *m/z* values enable biological insight. For very large MSI files and restricted computational resources, TOP3 and PR3D enable rapid tissue screening, after which selected sub-regions can be interactively interrogated with *m/z* mapping.

#### Enriched lipids in a mouse model

Figure 6 illustrates the identification of a lipid that is enriched in a sub-region of the Substantia nigra (SN) in an ABCA7-deficient mouse (A7ko) (*33*), highlighting the localized effects of ABCA7 loss. To explore region-specific lipid differences, we compared UMAP and SALO visualizations for mapping *m/z* values across brain sub-regions. While SALO successfully distinguished two sub-regions within a selected region of interest (ROI1), UMAP failed to resolve these differences. Within ROI1, we identified *m/z* 722.6 Da, which was strongly enriched in this region. A second *m/z* 885.6 Da exhibited a weaker, though noticeable, preference for ROI1. The *m/z* 722.6 lipid, detected in negative mode, is a candidate for several lipid species, including phosphatidylethanolamine (PE 34:1), phosphatidylserine (PS 32:0), and ceramide (Cer 42:2;O3 [M+OAc]-).

**Figure 6:**
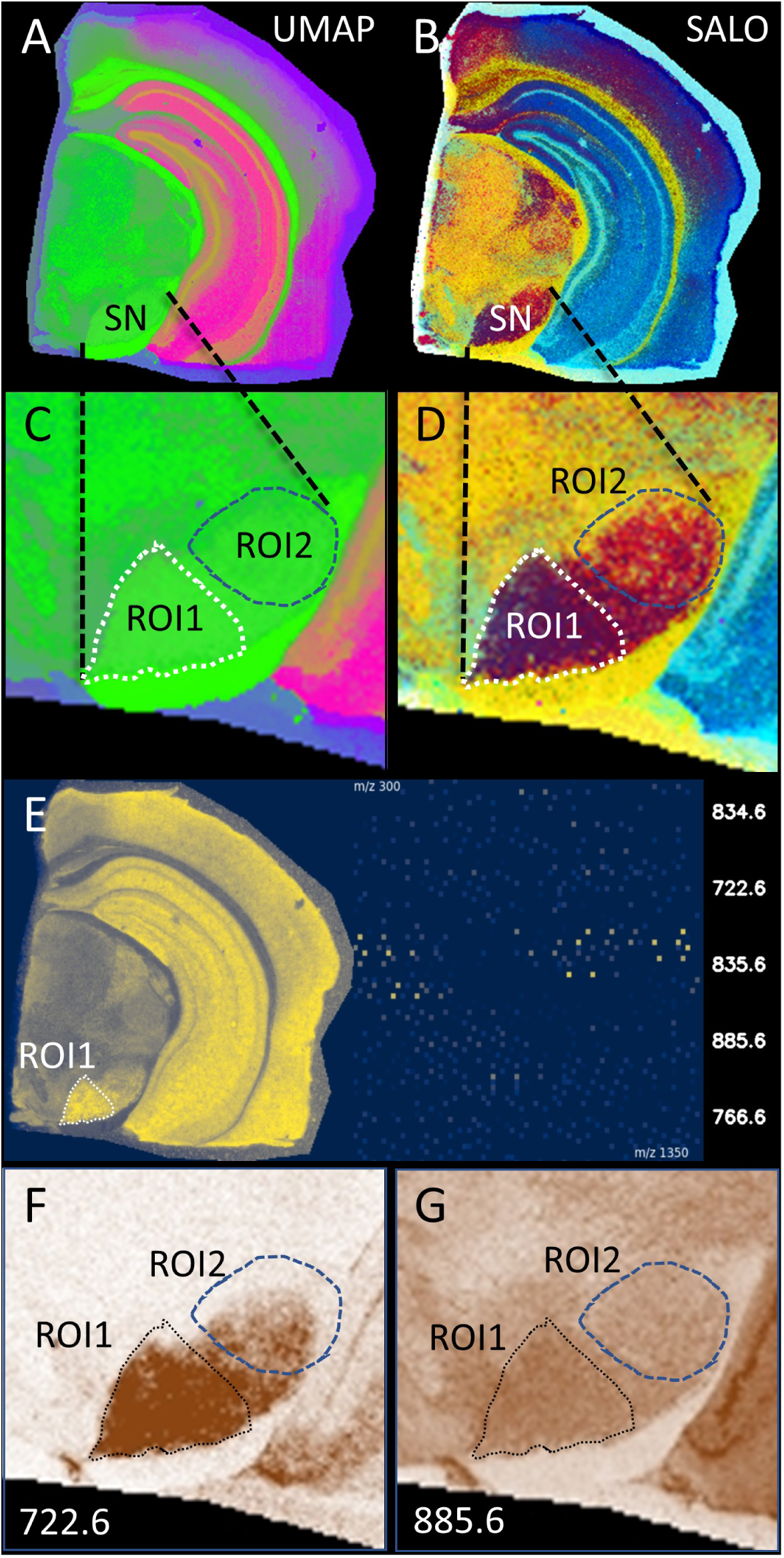
Index lipids reveal region-specific neuronal differences in an ABCA7-knockout mouse. (**A, C**) UMAP visualization, (**B, D**) SALO visualization identified two distinct sub-regions (ROI 1,2) within the Substantia nigra (SN), which were not resolved by UMAP. (**E**) The mapping analysis reveals the five most significant *m/z* values. The Virtual Pathology Stain and the SpecQR show the distribution of the *m/z* values. (**F**) *m/z* 722.6 is strongly enriched in ROI1. (**G**) *m/z* 885.6 is only slightly increased in ROI1.

#### Lipid composition of amyloid plaques mimicks neuronal composition

Figure S15 demonstrates the determination of *m/z* values associated with A*β* plaques. To investigate the lipid composition within these plaques, we examined the spatial distribution of *m/z* 885.6 (PI 38:4) using MSI-VISUAL. This lipid, known to be abundant in neuronal connections in the brain (*43, 44*), was also detected by us in spinal cord neurons. When comparing visualization methods, TOP3 successfully highlighted numerous A*β* plaques, while UMAP failed to resolve them. SpecQR presentation of spinal cord and brain datasets revealed shared *m/z* values between spinal neurons and cortical A*β* plaques. Ion images of *m/z* 885.6 in two AD mouse models showed this signal in large spinal neurons, A*β* plaques, and hippocampal neuronal connections. These findings suggest that *m/z* 885.6 is a physiological neuronal component that may accumulate in A*β* plaques as neuronal interconnections are destroyed.

### Integrated MSI-VISUAL workflow: rapid, combined ROI selection, comparison, and ROI-to-*m/z* mapping

We asked whether MSI-VISUAL can support fast, reliable ROI-to-*m/z* exploration. The framework provides a GUI for data extraction, multi-visualization generation, ROI selection/comparison, and mapping of significant *m/z* values, thereby reducing interpretation time and effort (Figure 7).

**Figure 7:**
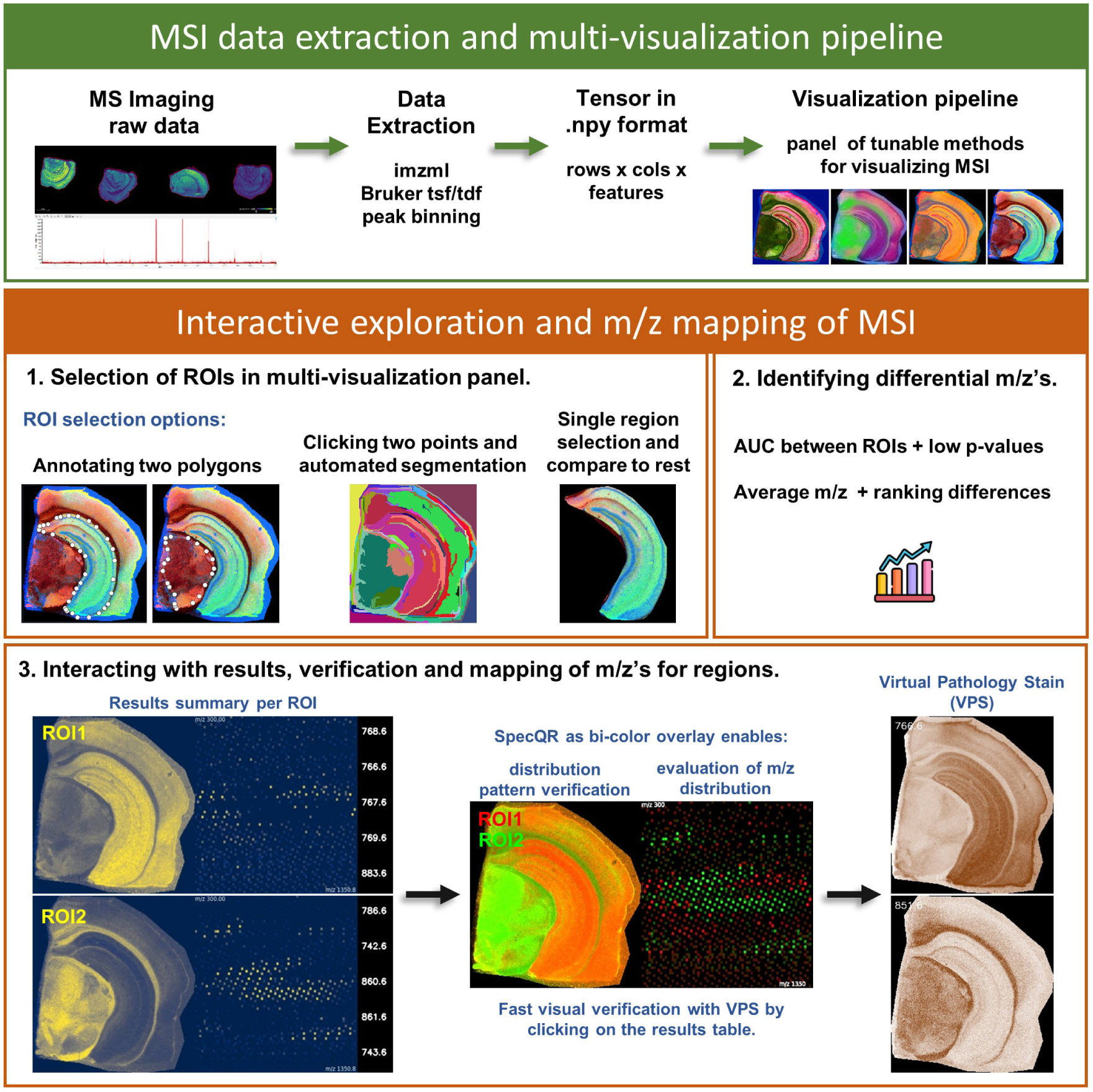
MSI-VISUAL workflow for interactive ROI-to-m/z analysis of MSI omics datasets. Main stages include: MSI data extraction and the visualization pipeline; interactive exploration; ROI selection; *m/z* mapping and results tables; and confirmatory representations, including monochrome images (Cividis color scheme (*46*)), bicolor (red-green) images, the SpecQR 2D graph, and Virtual Pathology Stains (VPS), which mimic immunohistochemical (IHC) stains and enable rapid visual evaluation of ROI properties.

First, users create a data pipeline that extracts raw MSI data (either in the open-source imzML format (*6*) or, additionally, in the Bruker hardware format) into a fixed-shape tensor of size **rows** ·**cols** · **N** in open-source .npy format. Here, **N** (the number of features per pixel) is determined by the total *m/z* range in the scan and the number of bins allocated per 1 Da *m/z* unit.

Multiple tunable visualizations are automatically or user-determined generated from the high-dimensional tensor to form a panel in which different visualizations highlight different microscopic, anatomical, or functional details. Users can then select ROIs directly in these visualizations via several interaction modes:

i) **Manual region annotation:** Users annotate polygons around ROIs in different visualizations of the same MSI scan.
ii) **Automatic segmentation with click selection:** Visualizations are first spatially segmented. We use the Felzenszwalb-Huttenlocher segmentation method (*45*) to partition MSI visualizations into sub-regions, which are then selected by clicking.
iii) **Point selection:** Users click individual points to focus on smaller structures.

After ROI selection, MSI-VISUAL compares regional *m/z* content and identifies significantly different intensities. Results are shown in a table and in SpecQR, which aggregates *m/z* spatial images on a 2D grid (Figure S18). Each square’s intensity is scaled by statistical score, and red-green contrast provides rapid visual verification of region-specific signals. Users may then select individual *m/z* values in the table to generate Virtual Pathology Stains (VPS) of single *m/z* images (often termed ion images), enabling visual confirmation that these *m/z* values correspond to the tissue pattern of interest.

## Discussion

Mass spectrometry imaging (MSI) is increasingly used in research and diagnostics, but its practical value depends on trustworthy visualizations that preserve biologically meaningful structure. Our framework addresses this need by combining structure-preserving dimensionality reduction with interactive *m/z* mapping and Virtual Pathology Stains (VPS), enabling rapid, pathology-like interpretation and hypothesis-driven exploration. MSI-VISUAL is designed to provide complementary visualizations rather than replace existing methods, because different views reveal different spatial-molecular details.

Current MSI workflows often depend on alignment to external histological stains for ROI definition (*5*). Our results indicate that continuous RGB visualizations plus direct ROI-to-*m/z* interrogation can reduce that dependency, while preserving compatibility with standard pathology workflows and immunohistochemical-style interpretation.

Across benchmark datasets, SALO and SPEAR show the strongest global-structure preservation in our benchmarks. SALO substantially outperforms UMAP (*18, 47*) (approximately 0.3 − 0.4 vs *>* 0.8 in our setting), and also outperforms PaCMAP (*23*) and PCA (*16*). This pattern is consistent with SALO’s direct optimization of global structure and non-linear mapping, in contrast to indirect or linear alternatives, and aligns with our previous observations with MiCS-LMC DR (*48*). The joint use of complementary distance behaviors further improves robustness for MSI-relevant signal patterns (*24, 47*).

At the same time, method trade-offs are important for practice. TOP3 is computationally lightweight and remains competitive – often outperforming UMAP in our tests – but does not reach SALO/SPEAR fidelity. This makes TOP3 a strong option for rapid screening and resource-constrained settings, with a clear future path toward block-wise/partial data loading because it operates pixel-wise. PR3D showed variable performance across METASPACE datasets, likely because its parameters were tuned to mouse brain data; automatic parameter selection is therefore a key next step (Figures S22 and S23).

Several limitations should be considered. A general limitation of all MSI visualization methods is that low-dimensional visualizations can overemphasize apparent local structure, creating visually salient details that may not reflect robust molecular differences in the original high-dimensional spectra. This risk is particularly relevant for complex non-linear dimensionality-reduction approaches. At the same time, there is also a risk of missing subtle but biologically meaningful details during dimensionality reduction. In MSI-VISUAL, we mitigate these limitations by coupling visualization with interactive ROI-to-*m/z* mapping and rapid MSI-native comparison, enabling fast verification of whether a visual feature is supported by underlying molecular signals. In addition to this confirmation and the use of a panel of complementary visualizations offered in MSI-VISUAL, diagnostic use routinely requires validation across additional sections/slides, orthogonal modalities, and independent methods. Furthermore, we note that current quantitative metrics for MSI visualization remain imperfect and may not fully capture biological relevance or pathologist interpretation. Improving these metrics, and systematically studying their correspondence with expert (e.g., pathologist) interpretation, represents an important direction for future work.

Another limitation is that optimization targets global rather than local structure, so local specificity can degrade in some cases (e.g., TOP3 assigning similar colors to unrelated regions). SALO and SPEAR also do not guarantee cross-image color consistency because they are non-parametric and depend on initialization and reference-point sampling. Benchmarking used mostly default hyperparameters (except UMAP distances), so additional tuning could improve both baseline and proposed methods, though this is often impractical for routine use.

For SALO specifically, fixed reference points are an additional constraint: re-sampling may improve quality but increases runtime and computational cost. Future work should balance adaptive reference strategies against practical performance.

Binning is another common concern in MSI. In our experiments, binning did not undermine truthful visualization and, in several cases, improved practical output quality for lightweight methods. On the mouse brain data, 5 bins performed as well as or better than higher bin counts for PR3D/TOP3, while advanced methods showed substantial robustness (Figures S20 and S21). This distinction is important: faithful visualization and high-precision mass identification are related but separate goals. For annotation and exploratory ROI analysis, binned data are often sufficient and computationally efficient; for precise identification, full-resolution extraction remains preferable, for example with Gaussian mixture modeling pipelines (*14, 34*).

Overall, MSI-VISUAL provides a fast and integrated workflow from visualization to ROI-based molecular interpretation. In our biological examples, this enabled detection of fine subregions and shared lipid patterns across neurons, plaques, and substantia nigra domains. These properties support both exploratory studies and time-sensitive diagnostic settings. MSI-VISUAL has already supported construction of a high-resolution lipid brain atlas with 123 annotated mouse brain regions (*34*); the next step is prospective clinical validation, including reproducibility testing across cohorts, instruments, and acquisition sites.

## Material and Methods

### The lipidomics MSI mouse brain dataset

The mouse brain MSI data files were generated using frozen hemispheres from four 100-day-old animals (C57Bl/6J genomic background). Two mice expressed combined human mutated APP and PS1-mutated transgenes (APPtg) for the generation of A*β* plaques (*36, 32, 49*). Frozen mouse hemispheres were coronally cut into 10 *μm*-thick sections using a cryomicrotome (CM1950, Leica, Nussloch, Germany) and placed on a single Intellislide (Bruker Daltonics, Bremen, Germany). The sections were dried in a vacuum chamber and sprayed at 30 ^◦^C with N-(1-Naphthyl) ethylenediamine dihydrochloride (NEDC) matrix dissolved to 7 mg/ml in 70% methanol using an HTX3+ TM Sprayer (HTX Technologies, Carrboro, NC, USA), as described by Andersen et al. (*50*). Matrix-assisted laser desorption ionization trapped ion mobility mass spectrometry imaging (MALDI-TIMS-MSI) was performed in negative mode using a timsTOF fleX™ mass spectrometer (Bruker Daltonics, Bremen, Germany). Mass calibration was performed using red phosphorus and ion mobility using ESI-L Low Concentration Tuning Mix (Agilent Technologies, Santa Clara, CA, USA) in ESI mode. For MALDI-TIMS-MSI analysis, a mass range of *m/z* 300 − 1350 and a range of inverse reduced mobility (1/K0) of 0.7 − 1.8 *Vs*/*cm*^2^ was used. The laser was operated at a frequency of 10 kHz with a burst of 200 shots per pixel and a spatial resolution of 20 *μ*m. Four mouse hemispheres were assessed in a single overnight run (Table S1). The mouse brain lipidomics dataset files are available at the ProteomeXchange PRIDE repository with the dataset number PXD056609.

### The human proteomics MSI dataset

The human brain MSI data files have been downloaded from the ProteomeXchange PRIDE repository with the access code: PXD049325 (*40*).

### Open-access metabolomics and lipidomics datasets

The METASPACE datasets used for the benchmarking and testing are listed in Table S2.

### Data extraction and pre/post-processing information

For extraction and binning of the mouse brain dataset, we used the ”extract bruker tims.py” script from MSI-VISUAL. For extraction and binning of slides from METASPACE, we used ”extract pymzml.py” from MSI-VISUAL. Binning can be adjusted to user needs and available computational power for later visualizations. We evaluated 5, 10, and 20 bins per *m/z* (respectively 0.2 = 200 mDa, 0.1 = 100 mDa, and 0.05 = 50 mDa precision). During extraction, *m/z* values are rounded to their closest bin and summed there. Using 5 bins showed the best visualization results for the lightweight methods PR3D and TOP3 compared with 20 bins; highly resource-dependent visualizations (e.g., UMAP) did not show significant differences between 5, 10, and 20 bins (Figure S20). Thus, the presented results were generated using 5 bins. When creating visualizations, we use TIC (*51*) normalization, which divides the intensity of each *m/z* value by the sum of intensities for that pixel. In all visualizations, we apply outlier clipping at the 0.01 and 99.99 percentiles of the data (Winsorizing) (*52*).

### Visualization methods

#### Notation and intuition

We denote by *x_u_* ∈ ℝ*^*N*^* the full MSI spectrum of pixel *u*, where **N** is the number of binned *m/z* features (one value per mass/charge bin). A visualization method maps each high-dimensional spectrum *x_u_* to a 3D color vector *y_u_* ∈ ℝ^3^. For a pair of pixels (*u, v*), we define distances explicitly as

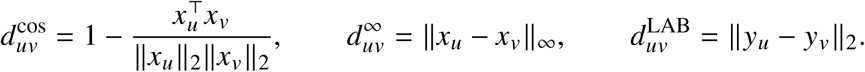

Here, 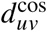 captures distributed differences across many *m/z* bins, 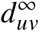 captures the largest single-bin intensity difference, and 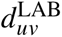 measures perceptual color separation in the visualization. Let P be the set of pixel pairs and let rank(·) assign larger ranks to larger distances. We then define

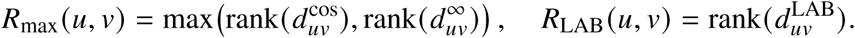

Intuitively, *R*_max_(*u, v*) is the input-space ”importance rank” of pair (*u, v*) under the stronger of the two MSI distances, and *R*_LAB_(*u, v*) is how strongly that same pair is separated in the final color image.

#### MSI terms used in this section

*m/z* denotes mass-to-charge ratio. A pixel ”spectrum” is the vector of ion intensities across all binned *m/z* values. ”TIC normalization” divides each intensity in a pixel spectrum by the sum of all intensities in that same pixel. ”LAB color space” is a perceptual color space where Euclidean distance better reflects human color difference than raw RGB distance.

#### Motivation for the saliency metric used in Saliency Optimization (SALO)

The saliency metric is used both as (i) an optimization objective in SALO and (ii) an evaluation metric for MSI visualizations. We designed it to satisfy the following properties:

i) **Preservation of important details.** Pairs of points that exhibit large distances in the input space but are mapped to small distances in the visualization should be penalized. This ensures that meaningful structural differences are preserved.
ii) **Higher weight for rare but significant differences.** Pairs of points with large distances in the input space should have a greater influence on the saliency score. While such pairs may constitute a small fraction of the total, they are critical for salience. Without this weighting, structures comprising a small percentage (e.g., *<* 1%) of all pairs might be completely overlooked; yet, the overall metric could still appear high. Additionally, points with very small input-space distances can artificially inflate the saliency score, despite not contributing to meaningful structure identification.
iii) **Integration of cosine and** *L*_∞_ **distances.** The metric should accommodate multiple distance functions to capture diverse biological structures. Some structures manifest through a few distinct *m/z* values with large intensity changes (better captured by *L*_∞_), whereas others involve coordinated changes across many *m/z* values with smaller individual intensity shifts (better captured by cosine distance). Both pattern types should be detected.
iv) **Incorporation of perceptual color differences.** The metric should account for human perceptual biases in color interpretation. For instance, subtle variations in shades of pink may be difficult to distinguish, even if the visualization encodes meaningful numerical differences. A saliency metric that aligns with human visual perception ensures that the detected structures are recognizable to users.

#### The Saliency objective definition

Using the previously defined input-space importance rank *R*_max_(*u, v*) and output-space rank *R*_LAB_(*u, v*), we define pairwise and global saliency as follows. For a tolerance parameter Δ, define the pairwise indicator

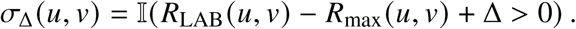

The Δ-saliency score is the weighted mean

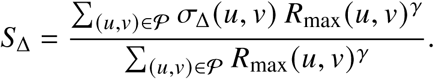

We report **Average Saliency** as

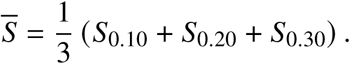

#### Saliency Optimization

We optimize SALO with stochastic gradient descent to maximize the saliency objective. Directly evaluating all pixel pairs is quadratic in the number of pixels, so we approximate Δ-saliency using fixed reference points. The optimization set is:

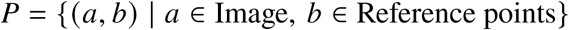

We therefore compute saliency on *P* rather than all pairs. Because exact ranking is not differentiable, we use soft ranking (a differentiable rank approximation) from (*21*), with the default parameters from the official PyTorch implementation (*53*). Replacing *R*_max_(*u, v*) by *R*^^^_max_ (*u, v*) yields

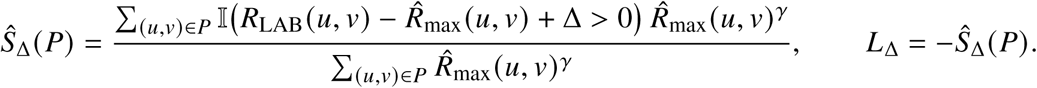

In all experiments, we set Δ = 0 and *γ* = 2 in the loss function, based on the hypothesis that this would enhance the detection of rare objects, such as plaques or neurons. However, these values could be further refined. As previously noted, this parameter assigns greater weight to larger distance ranks, which may help in identifying rare objects. A lower value of *γ*, such as 0 or 0.5, could potentially improve overall performance while reducing effectiveness in rare sub-groups. Further investigations into this trade-off would be valuable in future work. As a strategy for selecting reference points for optimization, we consider the following options:

i) **Lightweight core sets method** (*10*). This method is a fast sampling method that samples points for both diversity and coverage of the main clusters.
ii) **K-means++** (*11*), which selects points in a way that maximizes their spread: the first point is chosen randomly, and subsequent points are sampled with probability proportional to their squared distance from the nearest existing centroid.
iii) **Random sampling**.

Figure S24 shows the effect of different sampling strategies. For a large number of points, the difference is smaller; however, run time will increase (the complexity is proportional to the number of reference points). Cosine distance correlation performs better with K-means++; however, this is somewhat expected because the distance function in K-means++ is Euclidean.

By default, we use the lightweight core sets method (*10*) for sampling reference points because of its speed. For optimization, we use the Adam optimizer (*54*) with a learning rate of 1. The visualization is initialized with a random normal distribution in the range [-5, 5].

#### Spearman Rank Correlation Optimization (SPEAR)

SPEAR uses the same input/output distance definitions as SALO, but optimizes a simpler target: the Spearman rank correlation between *R*_max_(*u, v*) and *R*_LAB_(*u, v*) across pairs. In other words, it directly maximizes monotonic agreement between high-dimensional spectral dissimilarity and low-dimensional color dissimilarity. We use SPEAR as a simpler baseline to test the added value of the saliency objective.

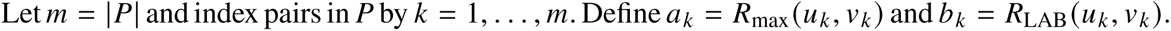

Then *ρ_s_* denotes the Spearman rank correlation coefficient between vectors *a*and *b*.

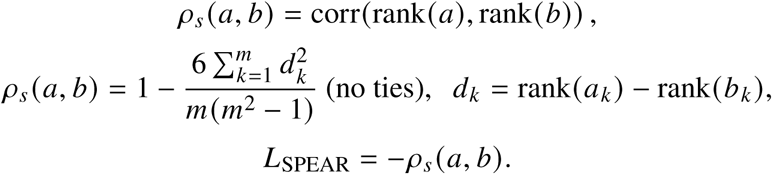

#### Lightweight visualizations

TOP3 selects the three highest intensity values per pixel and then maps them to the LAB color space. We implement this method efficiently using the Numba Python package. Our Percentile Ratio (PR3D) visualization is a generalization of TOP3, measuring how intensities are distributed. We define the PR function, which operates on pixel coordinates *i* and *j* for two fixed percentiles *p*_1_ and *p*_2_:

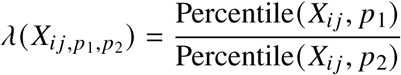

The *L*, *A*, and *B* channels in LAB space are then computed as follows:

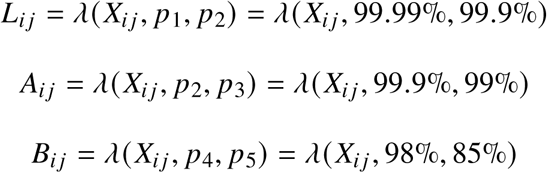

The percentiles *p*_1_, *p*_2_ and *p*_3_ were selected by a grid search on the first image of the mouse brain dataset (containing 5,255 features), evaluating 1,000 predefined combinations within the range [85%, 100%]. The values of *p*_4_ and *p*_5_ were manually adjusted to achieve the desired pink shade. Future work may focus on making these values differentiable for direct optimization or developing a more efficient search method.

### Interactive *m/z* mapping methods

#### Terminology for interactive analysis

A region of interest (ROI) is a user-selected set of pixels in the tissue image. A ”feature” corresponds to one binned *m/z* value. Statistical score refers to the effect-size or significance-based quantity used to rank features in pairwise ROI comparisons.

#### Region selection

MSI-VISUAL facilitates automatic selection of regions by interacting with the visualization. One method is based on the approach by Felzenszwalb et al. (*45*), a classical, efficient graph-based computer-vision algorithm that performs spatial segmentation of images into sub-regions, accounting for both spatial arrangement and visual appearance. An alternative method supported by MSI-VISUAL involves segmentation of color channels without using spatial information. In this case, each of the three color channels (Red, Green, and Blue) in the visualization is segmented into **N** sub-regions (with **N** = 5 by default, though this can be adjusted). Segmentation is achieved by digitizing each channel into **N** equally spaced bins in the range [0, 1]:

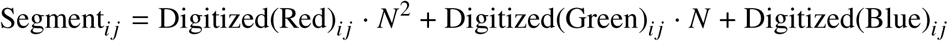

Here, *i* and *j* denote the pixel coordinates, Digitized(image) represents the quantization of an image or region into **N**bins, and Segment*_i j_* refers to the sub-region to which pixel (*i, j* ) belongs.

#### Region vs. region comparisons

For two selected ROIs, MSI-VISUAL compares each *m/z* feature between groups using the Mann–Whitney *U* statistic. We additionally compute p-values and retain *m/z* features with p-value *<* 0.05 as significantly different between ROIs. For interpretability, we report the corresponding effect size as AUC = *U*/(*n*_1_*n*_2_), where *n*_1_ and *n*_2_ are the numbers of pixels in ROI_1_ and ROI_2_.

#### Region vs. all comparisons

In this mode, one ROI is compared against all remaining pixels in the image (the complementary mask). To reduce computation while preserving global contrast, every second pixel in the complementary mask is sub-sampled.

#### Point-to-Point comparisons

In this mode, users select two pixels by clicking on them within the visualization. For each *m/z* feature, we compute the within-image rank of intensity at each selected pixel and then subtract these ranks. Features are sorted by this rank difference: positive values indicate enrichment in the first pixel, negative values indicate enrichment in the second pixel.

#### Presenting the Results: The SpectrumQR (SpecQR) visualization method

The goal of SpecQR is to provide an interpretable molecular ”fingerprint” of differences between two ROIs. The input to SpecQR is the ranked list of discriminative *m/z* features and their statistical scores.

In the first part of SpecQR, we generate a summary ion image (Cividis color scheme (*46*)) by averaging the five top-ranked *m/z* features. This gives a rapid spatial overview of the strongest molecular differences.

In the second part, we produce a 2D spectrum map across the full acquired *m/z* range. Each cell corresponds to one *m/z* feature and is color-coded by its score. This allows rapid visual inspection of broad versus focal molecular differences between ROIs. An in-depth example is shown in Figure S18.

Additionally, we provide a bi-color overlay image (ROI_1_ in red, ROI_2_ in green) for intuitive spatial comparison of region-specific signal patterns.

#### Presenting the Results: Interactive exploration of *m/z* values

To support interactive exploration, significant *m/z* features are presented as a ranked list. Users can click any feature to instantly generate a Virtual Pathology Stain (VPS; see next) for visual verification of its spatial pattern.

This workflow is designed as an iterative loop: identify candidate structures in the visualization, select ROIs, compare them with SpecQR, and then verify or reject candidate *m/z* features by checking their spatial maps.

#### Presenting the Results: The Virtual Pathology Stain (VPS)

To simulate a commonly used immunohistochemical (IHC) stain, the grayscale ion intensity images were mapped to a brown color scale. Total ion current (TIC) intensities of fully resolved *m/z*s were normalized for each *m/z* to 0–1 for all pixels, and a linear interpolation was performed for each pixel between white (RGB: 255, 255, 255) and dark brown (RGB: 139, 69, 19). This mapping ensures that low-intensity pixels remain light, while high-intensity regions are rendered in progressively deeper shades of brown, visually resembling 3,3’-diaminobenzidine (DAB) staining used in routine pathology diagnostics (immunohistochemistry stains).

### Quantitative evaluation of the MSI visualizations

#### Evaluation metrics

As the evaluation metric, we use Pearson distance correlation, which quantifies how well pairwise distances are preserved between high-dimensional and low-dimensional spaces. Specifically, we computed the Euclidean distances between all pairs of points (pixels) in the original high-dimensional MSI data and separately between the corresponding points in the three-dimensional RGB space used for visualization. We then compute the Pearson correlation between these two sets of distances. A higher correlation indicates better preservation of the global structure of the high-dimensional data in the reduced RGB representation. This approach has previously been applied to MSI visualization evaluation (*24*) and is a well-established metric to evaluate dimensionality reduction (*25, 26*). Following prior work in MSI visualization (*24*) and to assess performance under different distance metrics corresponding to different biological scenarios of *m/z* differences between regions, we computed correlations using both cosine and *L*_∞_ distances (Figure S2). The distance *L*_∞_ is defined as the maximum absolute difference in intensity between all *m/z* values between two pixels.

In addition, we report benchmarking with similar results for the Spearman rank correlation metric (Figure S6), and our Saliency metric (Figure S5), which have not been used before for evaluating MSI, which may be meaningful for future comparisons.

#### Benchmarking of the MSI visualizations

We evaluated the new visualization methods on two datasets:

i) Our mouse model brain dataset (Table S1) consisting of four brain hemisphere scans at 20µm resolution. This dataset represents controlled, high-quality MSI data, where special care has been taken in sample preparation and the measurement protocol.
ii) A diverse collection of six datasets from the METASPACE repository (*9*) (Table S2), covering various tissue types, low and high resolutions, different numbers of features, and different *m/z* ranges. Some scans were created with peak selection, reducing the number of features. Some image files are huge, making them computationally challenging to process.

Only our proposed methods (SALO, SPEAR, TOP3, and PR3D), along with PaCMAP, PCA, and NMF, were able to run on the full-size images using a high-performance machine with 320GB of RAM. To facilitate comparisons with all methods on the second dataset, we subsampled every second pixel from the large kidney (dataset 4) and lung (dataset 6) MS images.

#### Figure panels

The generated computational visualizations, ion images, SpecQR images, or parts of these were used in several figures by intention.

#### Ethical approval

Animal breeding and tissue harvesting was approved by the Department of Komparative Medicine (KPM) (IV2-2022).

## Supporting information

Response to reviews R1 / SciAdv

## Acknowledgements

The authors would like to thank Dr. Jorunn Stamnæs for performing the MSI measurements of the mouse brains and for technical discussions, and Thomas Brüning for cutting and preparing the mouse brain hemispheres and the H&E and immunohistochemistry stains. The mass spectrometry-based analyzes of mouse brains were performed at the Proteomics Core Facility at the University of Oslo/Oslo University Hospital, which is supported by the Core Facilities Program of the South-Eastern Norway Regional Health Authority (HSØ) and NAPI (www.napi.uio.no, NFR, Norway; 295910).

The authors would like to thank Dr.-Ing. Marcin Grzegorzek, Professor at the Institute of Medical Informatics, University of Lübeck/Germany, for supporting our work.

The authors would like to thank Manuel Liebeke (MPI Bremen, Germany), Michaela Schwaiger-Haber (Washington University in St. Louis, USA), Tingting Lu (South China Agricultural University, China), Brittney Gorman and Dušan Veličković (Pacific Northwest National Laboratory, USA) for sharing their datasets as open-access on METASPACE.

The authors would like to thank especially Lars Gruber, Thomas Enzlein, and Carsten Hopf (Hochschule Mannheim, Germany) for sharing their datasets directly with us and on METASPACE.

The datasets were used for benchmarking and in Figures 2, 5, S11,S14, S15, and S21.

## Author contributions

Conceptualization (J.G. and J.P.), Methodology (J.G. and J.P.), Software (J.G. and J.P.), Validation (J.G. and J.P.), Formal analysis (J.G. and J.P.), Investigation (J.G. and J.P.), Resources (J.P.), Data curation (J.P.), Writing - original draft (J.G. and J.P.), Writing - review and editing (J.G. and J.P.), Visualization (J.G. and J.P.), Supervision (J.P.), Project administration (J.P.), Funding acquisition (J.P.).

## Data, code, and materials availability

The mouse dataset files are available in PRIDE (https://proteomecentral.proteomexchange.org/cgi/GetDataset?ID=PXD056609).

We release the code as an open-source package on Zenodo (https://doi.org/10.5281/zenodo.19465293). This study did not generate new materials.

## Supplementary Materials

Figures S1 to S24

Tables S1 to S2

