## Supplementary material for "Truthful visualizations for mass spectrometry imaging enable high spatial resolution interactive *m/z* mapping and exploration": Response to reviews R1 / SciAdv

**Manuscript number: aed3650 , Gildenblat J. et Pahnke J.**

### Reviewer: 1

In this manuscript, Gildenblat and Pahnke introduce MSI-VISUAL, an open-source framework designed for fast analysis and visualization of MSI data. The authors focus on dimensionality reduction methods to visualize mass spectrometry data, aiming to preserve global structure when projecting high-dimensional MS data into low-dimensional visualizations. Specifically, they propose two new visualization strategies for this purpose: Saliency Optimization (SALO) and Spearman Optimization (SPEAR).

In addition, they present TOP3 and Percentile Ratio (PR3D) as memory-efficient and computationally rapid visualization approaches for large-scale datasets. The authors benchmark these new strategies against a wide range of existing dimensionality reduction techniques commonly used for MS data visualization and demonstrate superior performance in preserving global structure. Importantly, they show how these strategies enable more detailed visualization, facilitate differentiation of anatomical structures, and reveal pathological changes such as protein deposits and inflammatory infiltrates, and lipid/metabolite enrichment in specific anatomical or pathological regions.

Overall, MSI-VISUAL represents an exciting new tool with significant potential impact. It is likely to be of high interest to the broad readership of Science Advances. I view the tool and the advancements presented in this manuscript very favorably.

> We thank the reviewer for the supportive comments!

However, the authors need to address several aspects of the current manuscript before publication:

> We hope the following explanations and numerous changes in the manuscript will substantially improve the readability and presentation of the results.

**Writing and Structure:**

The manuscript is poorly written and lacks a clear structure.

> We thank the reviewer for this straightforward comment and used the opportunity to rewrite and restructure the entire manuscript accordingly.
>
> In general, we rewrote the introduction, the results, and the methods sections. For the results section, we:
>
> - Modified the examples.
> - Added an overview section that clarifies the improvements that we show and links them to the subsequent sections.
> - Rearranged and grouped different results into quantitative improvements, qualitative improvements, scalability improvements, and workflow improvements.
> - reframed subsections to follow a clearer **question → approach → result → significance** narrative.
> - Reduced verbosity by tightening repetitive and overly long paragraphs.
> - Improved transitions between result blocks to enhance logical flow.
> - standardized language to a more formal, scientific tone.
> - Corrected grammar.
> - Renamed several results subsection titles to be **outcome-focused rather than method-focused**, improving readability and scanability.
> - Moved some examples to the supplementary materials to make sure the remaining examples are more focused and clear.

• The introduction should be more concise.

We thank the reviewer for this comment. Following it, we completely rewrote the introduction, and it is now 3 times more concise.

• The results section could be streamlined to focus on fewer key examples that clearly illustrate the advances provided.

We thank the reviewer for this suggestion and followed it. The results now have fewer key qualitative examples, which we revised and that we hope are easier to follow, while the supplementary materials contain many more examples.

• Some sections are cryptic and unclear (e.g., "iv) Detecting single neurons" on page 12).

We thank the reviewer for this opportunity to improve the sections. We improved both section titles (including the section mentioned for detecting single neurons) and the content of those sections.

- We revised the five Results subsection labels to be explicitly descriptive and outcome-focused (instead of terse/ambiguous phrasing).
- We performed a targeted readability edit of the corresponding section text, preserving scientific content while simplifying sentence structure and improving flow.
- We clarified technical statements in the lesion-differentiation and single-cell sections to make goals, methods, and outcomes more immediately understandable.
- We standardized terminology and style across these sections.

Overall, we hope these edits improve accessibility for reviewers without changing data interpretation or conclusions.

• A better description of where the new visualization methods outperform existing ones is needed.

We thank the reviewer for this comment and, following this, rewrote the results section. It is now split into sections that show:

1. Quantitative improvements. Here, we demonstrate on quantitative metrics previously used in the field for MSI visualizations that the new visualizations clearly improve.
2. Qualitative improvements: We show several examples where the new visualizations are better, and then also show examples of how we can use those for creating biological insights.

   In addition, we modified the examples.

3. Scalability improvements: We show how this allows faster visualization of large data.
4. Workflow improvements: We show how, with everything combined, and the MSI-Visual interactive workflow, we can quickly create many biological insights.

**Methodological Clarity:**
As an expert in the computational domain, I found it difficult to fully grasp the technical details of the method. The reasons are manifold:

• Certain aspects of the method are not clearly explained. For instance, how is RYLAB generated?

We thank the reviewer for this comment, and following this, we revisited the methods section in depth (please see the list of changes below). In particular, we also addressed RYLAB and hope the notation is simpler and clearer now.

• All equations lack relationship signs (e.g., equal signs), making them nearly impossible to interpret.

We thank the reviewer for this comment. The problem occurred due to an Overleaf formatting bug and was addressed as soon as it was noticed. We deeply apologise for having missed it in your manuscript pdf version.

• The methods section should be expanded so that non-MSI experts can follow the methodology.

We thank the reviewer for this comment, and expanded and rewrote sections of the methods section:

- Added a clearer Notation and intuition block with explicit definitions of x_u, y_u, pair set \mathcal{P}, and rank operator.
- Added explicit distance equations for the distances in the high-dimensional data in the visualizations, and plain-language interpretation of each.
- Clarified ambiguous notation by explicitly defining R_max and R_LAB and linking to earlier shorthand.
- Removed duplicate R_max definition later in Methods and replaced with a forward reference.
- Rewrote Saliency objective to be shorter text + more formal math.
- Improved the Spearman objective notation.
- Aligned optimization notation with the new objective.
- Improved MSI jargon accessibility by adding short definitions (m/z, spectrum, TIC normalization, LAB space).
- Expanded interactive-analysis terminology (ROI, feature, statistical score).
- Rewrote Region/Point comparison paragraphs to be clearer for non-experts.
- Clarified significance workflow in ROI comparisons: ranking/effect size and significance threshold (p-value < 0.05).
- Updated stats wording from AUC-first to Mann–Whitney U first, with explicit equivalence AUC = U/(n_1 n_2).
- Rewrote SpecQR/VPS text for clearer interpretability and workflow framing.
- Applied consistency cleanup (capitalization, hyphenation, wording consistency, section header style).
- Applied typography polish in Methods (spacing/formatting of units, p-value formatting, minor phrasing cleanup).

• Some procedures are not explained at all. For example: How are statistically significant differential m/z values determined (ANOVA?) For Cosine and L∞ distances, are only pairs with significantly differential m/z assessed? If not, why?

We thank the reviewer for the opportunity to clarify this, and we added clarifications for how the m/z values are selected in the region vs regions comparisons methods section (Mann-Whitney U statistic on each m/z, between the two regions).

The cosine and L∞ distances are used on all pairs. These distances are not used in the context of the interactive region differentiation. Instead, they are used for metric computation (the correlation metric) and internally for optimizing the SALO and Spear methods. There, it is used with all m/z's, since it is done in the context of point-to-point comparisons, and not comparing two regions that were selected by the user.

**SpecQR Clarification:**

I have difficulty understanding the purpose of SpecQR. I understand the need to highlight the m/z value that differs most for an ROI, but what additional insight does 2D plotting provide?

The SpecQR code enables the fast overlay (or as a single layer) of information for **many** m/z at once - actually the whole MS spectrum, presented from low to high m/z.

The user can also choose two different ROIs to confirm the unique m/z pattern of each ROI. It is similar to fluorescent images in biological research, where co-expression of two proteins is visualized in the same image.
Here, the m/z information from the ROIs, e.g., ROI1 in green and ROI2 in red, results in yellow if expressed equally. If not, either green (preference for ROI1) or red (preference for ROI2) appears in the m/z SpecQR pixel.
The presented example shows a unique, ROI-specific pattern for **multiple** m/z values and enables a fast visual control of two ROIs.

Here, a magnified example of the SpecQR:

from the lowest m/z value

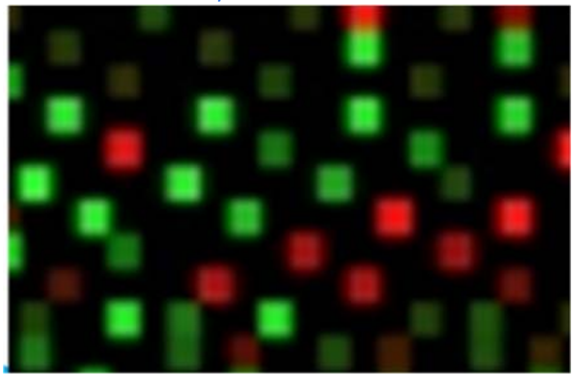

-> line-wise (0.2 Da per pixel)
-> highest m/z value

Each pixel represents one m/z value bin. The statistical score is coded as color intensity. The SpecQR could be used to evaluate the specific m/z values by clicking on the pixels.

Figure 4: How are the polygons drawn—manually or automatically? If manually, there may be bias.

We thank the reviewer for this helpful comment

The polygons in Figure 4 are drawn manually. MSI-VISUAL can also automatically draw ROIs by clicking a pixel. Both options allow the comparison of the ROI's m/z content later.

We agree that manual annotations may introduce observer-dependent bias and may differ from boundaries produced by automated segmentation methods. However, we emphasize that the intended use case of our visualizations is to support expert-driven analysis. In practice, pathologists interact with such visualizations to delineate regions of interest based on perceived biochemical differences.

From this perspective, we believe that the key question is not whether the boundaries match an automated segmentation, but whether the visualization makes relevant transitions between regions clearly visible to a human expert. Methods that better expose these boundaries are therefore more useful in downstream workflows. Figure 3 (Improved detection of sub-nuclei in mouse brains) illustrates this scenario, where an expert pathologist uses the visualization to identify and annotate meaningful region boundaries within an anatomical region.

Figure 8: How was it determined that panels (c) and (e) highlight vascular leakage?

The human brain MSI dataset from our colleagues in Mannheim (UCL brain bank sample) included in this specific analysis, consisted of a peptidomics dataset.

Vascular leakage was identified through the plasma albumin peptides (m/z 3.4kDa) and the morphology - lakes of albumin around vessels, whereas the ß-amyloid deposits were determined by a specific peptide composition (m/z 4.0-4.4kDa) and the plaque-like appearance.

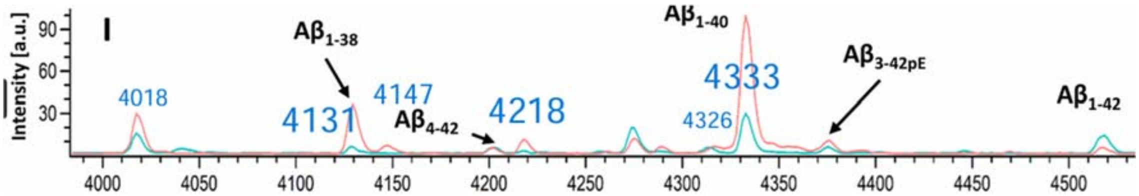

**Manuscript number: aed3650, Gildenblat J. et Pahnke J.**

### Reviewer: 2

The manuscript by Gildenblat and Pahnke describes a new platform termed MSI-VISUAL for analyzing mass spectrometry imaging datasets. This new platform used existing datasets to highlight different visuals from other statistically driven outputs to yield four different visualization strategies termed SALO, SPEAR, TOP3, and PR3D. The latter two methods employ binning as a form of data reduction to facilitate computationally rapid visualizations.

We thank the reviewer for the very helpful comments. **In response, we have made substantial revisions, including a full reorganization of the manuscript and a comprehensive revision of the Introduction, Results, Discussion, and Supplementary Material.** We hope these changes provide a clearer presentation of the contributions of MSI-VISUAL and a more explicit discussion of its limitations.

Regarding binning, we use user-defined binning across all visualization methods. While binning has traditionally been viewed as a limitation due to potential loss of spectral resolution, we show that, for visualization purposes, appropriate binning can improve robustness and computational efficiency without compromising interpretability.

We emphasize that binning should be selected based on the specific dataset and visualization goals. To illustrate its effects, we have included additional examples in the Supplementary Material (Figures S20 & S21).

While the resulting images are works of art, this reviewer has a hard time reconciling whether this is 'better.'

We thank the reviewer for the comment and the opportunity to clarify. We have substantially revised the Results section to better articulate the improvements provided by MSI-VISUAL.

We now distinguish four types of improvements:

**Quantitative improvements.**
To assess visualization quality in a user-independent manner, we performed computational benchmarking across multiple dimensionality reduction methods (including approaches not previously applied to MSI). Using established MSI visualization metrics, we show that the proposed methods improve global structure preservation.

**Qualitative improvements.**
For morphology-driven analysis (e.g., by pathologists), we present revised and more focused examples highlighting structural details revealed by the new methods. **Figure 1 has been redesigned to clearly illustrate improvements over ion images, as well as commonly used approaches such as K-means, PCA, and UMAP.**

**Scalability improvements.**
Large MSI datasets are often limited by memory and runtime constraints in existing methods (e.g., UMAP, K-means). We therefore introduce PR3D and TOP3, which enable efficient analysis of large datasets while maintaining high visualization quality.

**Workflow improvements.**
We demonstrate how the integration of these methods within the MSI-VISUAL framework, together with interactive ROI-based m/z mapping, enables rapid exploration and generation of biological insights.

We agree that no single visualization can serve as a universal ground truth. Accordingly, MSI-VISUAL provides multiple complementary visualizations, each capturing different aspects of the data. These views are intended to support exploratory analysis and can be further validated through downstream statistical interrogation. **We now further clarify this in the manuscript.**

**We do not claim that the proposed methods universally replace existing approaches (and further clarified this now in the manuscript in the results overview section and the discussion)** , but rather that they provide complementary advantages in many relevant scenarios.

The authors state several times that identification of m/z's is beyond the scope of this manuscript and that the majority of the datasets are taken from public repositories.

We thank the reviewer for this comment. Precise m/z identification can be performed using established approaches such as Gaussian mixture modeling (GMM), and we provide links to analyses conducted with MSI-VISUAL.

We note, however, that the primary focus of this work is on developing improved visualization strategies that facilitate exploration, comparison, and hypothesis generation in MSI data. These visualizations are intended to guide downstream analyses, including detailed m/z identification and interpretation, rather than replace them.

While the resulting images are indeed different, it is unclear if these represent 'ground truth' or are just simply forcing data to separate computationally. Further validation to highlight that these differences that the four new visualizations manifest are biologically meaningful would be important to show with orthogonal experimentation.

We thank the reviewer for this important comment. We fully agree that establishing biological relevance is essential when introducing new visualization approaches.
Following this comment and the comment below about orthogonal data, we added a supporting IHC figure (from the same mouse model) to the Supplementary Materials, Figure S17. **We expand at length in the comment below.**

**Additionally, we have included a limitations section discussing diagnostic use, emphasizing the need for complementary visualizations, validation across additional sections, and integration with orthogonal modalities.**

Below, please find a list of major and minor critiques for the author's consideration.

We thank the reviewer for the helpful comments.

**Major**

1. **Page 3**. It would be helpful to address how this approach takes into account hot spots, which definitely exist as a point of variation within these datasets, versus the ability to define actual subtle differences within an ROI within the data.

We thank the reviewer for this important point. Handling hotspots while capturing subtle intra-ROI variation is central to our method.

**Following this comment, we have clarified this in the Introduction, expanded it in the quantitative results, and further discussed it in the Discussion. We also revised the Methods section to better describe the optimization procedures and how global structure preservation is enforced.**

Conventional approaches such as PCA, UMAP, and K-means are primarily driven by dominant sources of variation ("hotspots"), such as major anatomical differences. As a result, they tend to emphasize separation between large regions (e.g., cortex vs. hippocampus), often at the expense of resolving finer intra-region variation.

In contrast, our method is **specifically designed to capture subtle intra-ROI differences** while remaining robust to dominant variation. This is achieved through two key components:

**(1) Sensitive distance metrics.**
The distance functions used in SALO/SPEAR are designed to detect small but meaningful differences between spectra, enabling sensitivity to subtle variations even when overall profiles are similar.

**(2) Global structure preservation.**
Our approach preserves global relationships by maintaining the relative ranking of distances to reference points. This prevents nearby points with subtle differences from being collapsed in the embedding, a common limitation of standard dimensionality reduction methods.

Importantly, the method preserves relative relationships rather than introducing arbitrary separations, which we validate quantitatively using global structure metrics.

Together, these components allow the method to avoid being dominated by hotspots while faithfully representing fine-grained variation within regions of interest.

2.. "Many m/z features are shared across different sub-regions and nuclei, which can bias visualization methods to assign them **the same color**, as is done in clustering methods." This is a grand statement; there might be some reasons m/z would be shared across tissue regions, and this could still be important, especially for multimodal imaging considerations. Microscopy still continues to offer high spatial resolutions for certain structures within tissues. Please consider changing this claim.

**We thank the reviewer for this important comment and agree that the original statement was overly broad, and following this comment have rewritten the introduction.** Shared m/z features across regions can reflect meaningful biological processes and are not inherently a limitation—particularly in multimodal settings where complementary techniques (e.g., microscopy) provide additional spatial context.

To address this, we have revised the Introduction to better reflect this nuance. Specifically, we now emphasize that relevant biological differences may arise either from coordinated small changes across many m/z values or from larger shifts in a smaller subset of ions.

Our goal is therefore not to discount shared m/z signals, but to ensure that visualization methods remain sensitive to subtle yet meaningful differences, especially in cases where regions share much of their molecular content. In such settings, important variation may be driven by relatively small changes that can be obscured by methods that overly average or emphasize dominant signals.

This motivation underlies our use of multiple distance metrics (e.g., cosine and $L_\infty$ distances) in SALO and related methods. For example, amyloid plaques (Figures 4, S9, S11, S15) share much of their m/z profile with the surrounding cortex but differ in a small subset of ions; methods sensitive to such localized differences are better able to resolve these structures.

We hope this revision clarifies that we aim to capture both shared and differential signals, while improving sensitivity to subtle but biologically meaningful variation.

3. **Page 6**, results section. Can the authors comment on which types of sources/mass analyzers were used to build the pipeline, and their recommendations for usage?

We thank the reviewer for this helpful question.

We developed and validated the pipeline using our own datasets acquired on a Bruker timsTOF flex instrument. To ensure general applicability, we additionally evaluated MSI-VISUAL on publicly available datasets from METASPACE generated using a range of different instruments.

The pipeline is designed to be vendor- and platform-independent. It supports any data that can be exported in the standard imzML format or accessed via vendor-specific export tools (e.g., Bruker DLL). These data are converted into an open-access array format (NumPy), with user-defined binning, and then processed within the MSI-VISUAL framework.

Based on our evaluation, we do not observe limitations tied to specific mass analyzers. The method is therefore broadly applicable across MSI platforms, provided standard data export is available.

For instance, would someone with an axial MALDI-TOF be able to use this software, or was it mainly built around a MALDI-QqTOF (timsTOF flex)? This would be helpful to clearly delineate any possible limitation considerations based on hardware.

We thank the reviewer for this clarification.

The pipeline is designed to be vendor- and platform-independent and can be applied to data from different MSI instruments, including axial MALDI-TOF systems (we used Neoflex and Rapiflex data for other projects), provided the data can be exported in a standard format (e.g., imzML) or via vendor-specific tools.

Our development and testing included data from a timsTOF flex instrument as well as publicly available datasets from multiple platforms (e.g., METASPACE), supporting its general applicability.

While we do not observe inherent limitations tied to specific hardware, performance may depend on factors such as data quality, spectral resolution, and preprocessing choices (e.g., binning). We have clarified this in the revised manuscript.

4. It would be helpful to define what 'sufficiently different' means on page 6 for the ROI search. Do the authors mean intensity? Also, does this algorithm take adducts and isotopes into account?

We thank the reviewer for this helpful comment.

Following this comment, we have clarified the wording in the manuscript: “sufficiently different” refers to **statistically significant differences in signal intensity** between regions of interest.

In practice, intensities are first binned during data preprocessing, and ROI comparisons are performed on these binned signals. The binning resolution can be adjusted by the user depending on the desired trade-off between spectral resolution and computational efficiency. In our experiments, we typically use a binning scheme corresponding to ~0.2 Da resolution (e.g., 5 bins), which we found to provide a good balance.

Regarding adducts and isotopes: these are not explicitly modeled or annotated within the current pipeline. Instead, they are implicitly included within the binned m/z signals and therefore contribute to the ROI comparisons. We note that more detailed handling of adducts and isotopic patterns could be incorporated as a future extension of the framework.

5. A concern with the SALO visualization is that the data methods are non-linear. This seems like it could assign more weight to a region of pixels that may not reflect biological heterogeneity. It is difficult to reconcile how to visualize differences in regions versus how to weight areas that are driven by biological heterogeneity. An important consideration for using segmentation or k-means in Cardinal is that these differences match histological differences well. I am not convinced yet by page 7 of this submission that non-linear visualizations will reflect true biological variation.

We thank the reviewer for raising this important point. We agree that nonlinear methods can, in principle, distort relationships and potentially emphasize patterns that do not reflect true biological heterogeneity.

**In response to this comment, we have addressed this point in the Discussion (Limitations section), revised Figure 1, and emphasised more in the results section.**

**We note that all MSI visualizations have inherent limitations, and therefore, the use of multiple complementary visualizations, interactive m/z mapping for verification, and, where available, external modalities and histological references, will be important for diagnostic applications.**

Our goal with SALO is not to replace established approaches such as clustering (**now further emphasized in the results and discussion)**, but to complement them. In contrast to clustering methods like k-means, which impose discrete boundaries, SALO produces continuous visualizations that can reveal intra-region variation. As illustrated in the **new Figure 1,** this allows finer-grained inspection of spatial structure that may be missed by hard clustering, as well as by linear methods such as PCA, which primarily capture dominant sources of variation.

To address concerns about nonlinearity, SALO explicitly incorporates global structure preservation, ensuring that relative relationships between points are maintained in the embedding. This reduces the risk of introducing artificial separations and is supported by improved performance on established structure-preservation metrics.

We also demonstrate this **in the new revised Figure 1**, where we compare linear methods such as PCA and k-means with the proposed visualizations. In addition, we quantitatively and qualitatively compare SALO to both clustering-based and continuous approaches (e.g., PCA, NMF) across multiple datasets, where it shows consistent improvements.

We acknowledge, however, that **existing evaluation metrics** in MSI visualization are still imperfect and may not fully capture biological relevance or pathologist interpretation.

Importantly, SALO is used within the broader MSI-VISUAL workflow, which includes complementary methods (e.g., k-means, PCA). The visualization serves as an exploratory tool: candidate regions identified visually can be interactively interrogated, and their validity assessed using statistical comparisons (e.g., Mann–Whitney U tests on m/z features). If a visually distinct region is not supported by meaningful molecular differences, this becomes apparent during downstream analysis.

We therefore emphasize that SALO is intended to enhance, rather than replace, existing analysis strategies—providing continuous visual cues that can be validated through complementary methods and quantitative interrogation.

6. I would have liked to have seen the claims supported by orthogonal data, such as an LCM for the Abeta plaques presented on page 10. If this workflow is superior, it would be important to provide ground truth examples of results with orthogonal experiments.

**We thank the reviewer for this important comment. We agree that orthogonal validation is valuable for confirming biological interpretation. In response, we have included an IHC staining from the same mouse model as an orthogonal reference to support the biological interpretation of the observed structures.**

In this work, our primary goal is to improve the visualization of MSI data, enabling a more faithful representation of spatial molecular patterns. To support the biological relevance of the observed structures, we rely on two complementary aspects.

First, the visualizations are consistent with known morphological and molecular features. For example, amyloid plaques are identified based on their characteristic spatial morphology together with known lipid signatures, such as GM3 (m/z 1179.7–1179.8), which can be assessed using MSI-VISUAL's ROI-based analysis tools. Similarly, neuronal structures can be identified through their spatial organization and associated m/z patterns. **In addition, we provide a corresponding histological reference (H&E staining)** in the **Supplementary Material (Figure S16),** enabling comparison between MSI-derived structures and tissue morphology.

As an orthogonal reference, we include **(and added to the supplementary material, Figure S17)** a representative IHC stain of amyloid plaques (4G8 staining, 100-day-old APP transgenic mouse), which illustrates the characteristic morphology of these structures:

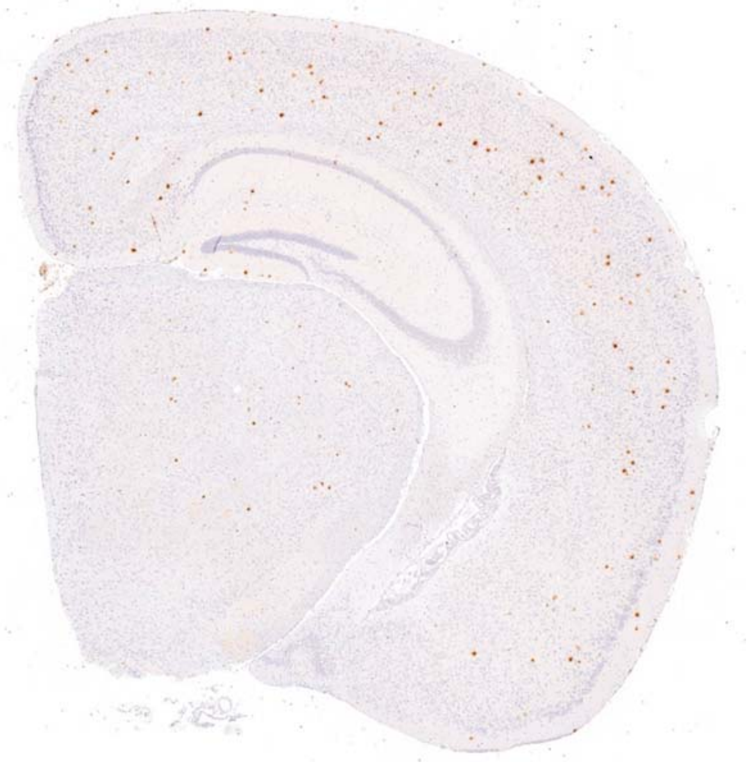

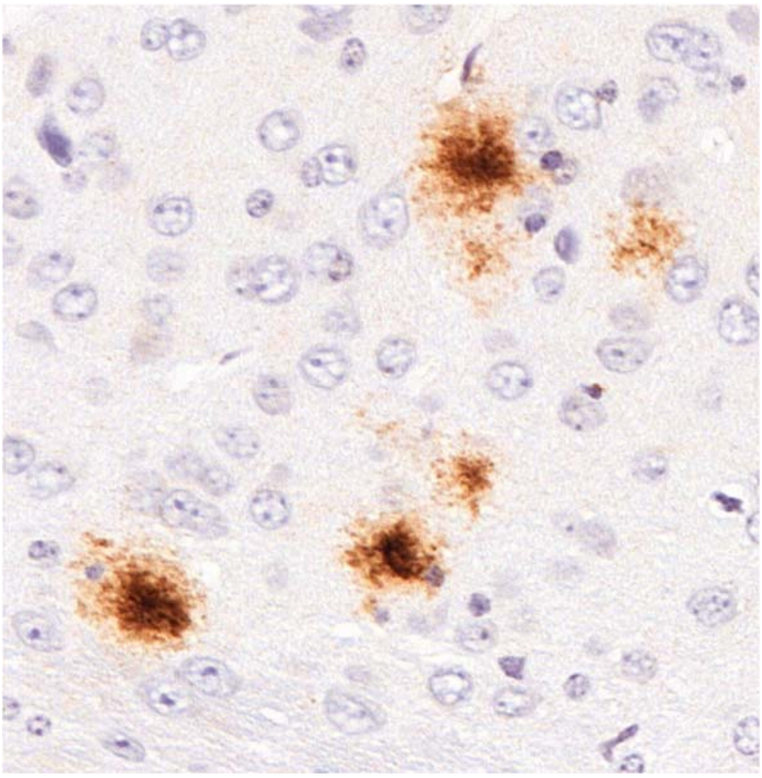

Second, MSI-VISUAL is designed as part of an interactive workflow, where **visualizations serve as an entry point** for discovery. Candidate regions identified visually can be further interrogated using statistical comparisons (e.g., Mann–Whitney U tests on m/z features), allowing validation of whether observed differences are supported by meaningful molecular variation. ROI-based analysis further enables the extraction of characteristic lipid profiles that support the identification of pathological structures.

These features can also be visualized using our virtual pathology stains (VPS), which map selected m/z values to spatial patterns analogous to targeted staining. The corresponding MSI-derived VPS, highlighting plaque-associated lipid signals (e.g., GM3 m/z 1179.7), is shown below for comparison:

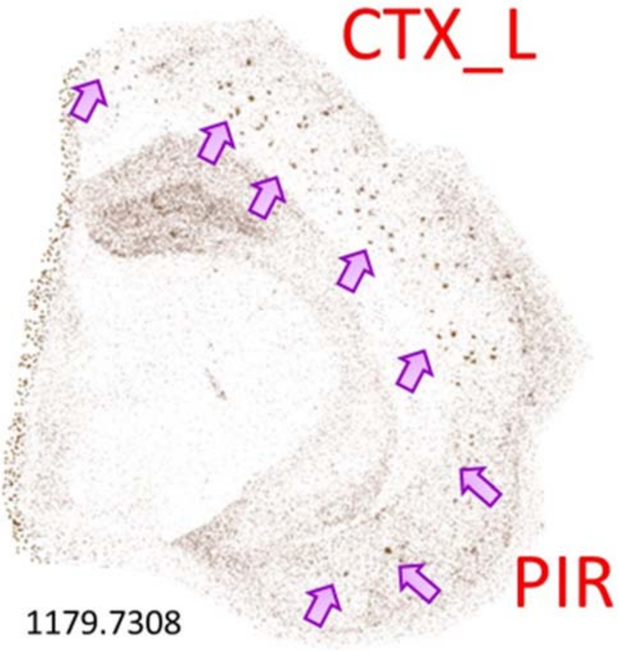


We note that, in the context of well-characterized structures such as amyloid plaques, the observed patterns are consistent with established morphological and molecular knowledge, including routinely performed IHC staining.

We agree that additional orthogonal validation (e.g., histology, IHC, MALDI-IHC, or LCM-based approaches) would further strengthen the findings, particularly for previously unknown structures, and represent an important direction for future work.

Overall, our results demonstrate that the proposed visualizations reveal structures that are consistent with known biology and can be systematically validated within the MSI framework using established m/z signatures.

**We have also added to the discussion section that, for diagnostics, checking with more data and modalities is essential.**

7. It might be helpful if the authors, instead of just including pictures of the segmentation, could point to specific m/z's that helped to generate this level of differentiation for Figure 3.

We thank the reviewer for this helpful suggestion and agree that linking visual features to specific m/z values is important.

In MSI-VISUAL, this connection is enabled through the interactive exploration framework. As shown in Figure S11 for the kidney or brain tumor example, users can select regions in the visualization and directly identify m/z values that differentiate these regions. **The specific m/z's are shown in the figure (list).**

This allows users to move from visual patterns to specific molecular drivers and verify the biological relevance of observed structures.

Our goal is for the visualization to serve as an entry point for discovery, rather than a standalone interpretation. The improved visualizations (e.g., in Figure 1, S8, S10, S19) arise from better preservation of global structure and the use of multiple distance metrics, which together highlight biologically meaningful regions. These regions can then be interrogated interactively to identify the contributing m/z features.

We note that, unlike linear methods such as PCA and NMF—or approaches like TOP3—nonlinear methods such as SALO and UMAP do not provide a direct, built-in mapping from pixels to specific m/z contributions. We agree that providing such mappings explicitly would be valuable, and view this as an important direction for future work, potentially through parametric extensions or post hoc interpretability methods.

Finally, we draw an analogy to pathology workflows: MSI visualizations can be viewed as analogous to histological overview stains, highlighting regions of interest, while individual ion images, or our DAB-like VPS (specific m/z values), act as "virtual immunostains" that explain and validate those regions. MSI-VISUAL is designed to support this two-step workflow—visual discovery followed by molecular interpretation.

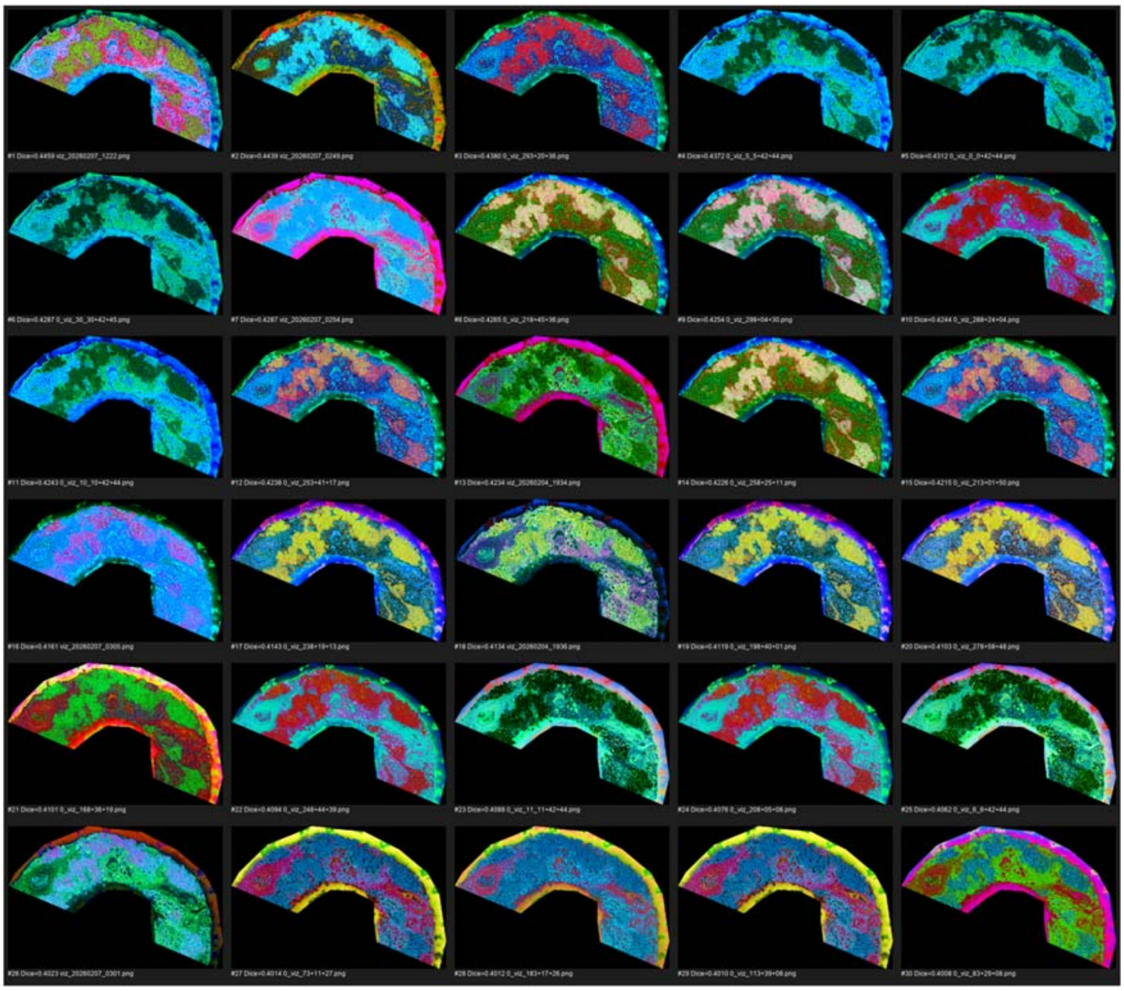

In pathology diagnostics, histological overview stains are generated using various color stains. MSI visualizations are meant to fulfill the same purpose. Immunostains are the ion-images of the MSI data (specific m/z) = virtual pathology stains (as exemplified for GM3 1179.7, see above).

8. **Page 24**. The comment on PR3D not performing well on all METASPACE datasets should be elaborated upon. Why does the method not perform well? It would be helpful to provide guidelines for when the method does and doesn't work well with respect to instrumentation or experimental parameters.

We thank the reviewer for this comment and have expanded the discussion in the manuscript accordingly.

PR3D is designed as a computationally efficient method that enables rapid visualization, even for very large MSI datasets. As shown in the new Figures S22 & S23, it can reveal different structural details depending on the chosen percentile parameters. Both tuned and non-tuned configurations (e.g., mouse brain vs. human kidney) illustrate these effects.

The main limitation arises from the **fixed percentile selection**, which may not generalize well across heterogeneous datasets such as those in METASPACE. These datasets vary substantially in terms of signal distribution, number of m/z features, and overall data quality. As a result, a fixed percentile configuration may either suppress relevant variation or overemphasize noise, leading to suboptimal visualizations.

In practice, we observe that PR3D performs well as a **fast preview or exploratory tool**, particularly for large datasets where more computationally intensive methods are not feasible. However, optimal performance may require adapting the percentile parameters to the specific dataset.

We therefore suggest that percentile selection should ideally be guided by data characteristics or quantitative metrics, and we consider automated parameter tuning an important direction for future work.

**Minor**

1. **Page 2**. While I appreciate that the focus is likely on open source programs, vendor-specific software, such as SCiLS, does offer support for paying customers and has recently implemented UMAP https://download.scils.de/scilslab/changelog.html

We thank the reviewer for this helpful comment. We now cite SCiLS Lab in the Introduction to acknowledge the role of vendor-supported software, including its recent implementation of UMAP.

While this work focuses on open-source methods to promote accessibility and reproducibility, we recognize that both commercial and open-source tools contribute to MSI data analysis.

2. m/z should be in italics throughout for convention.

We thank the reviewer for this comment. We have updated *m/z* to italics throughout the manuscript.

3. Top of **page 8**. Can the authors quantify 'larger'? Do they mean the number of pixels or high mass resolution data with a set number of pixels? A frame of reference is needed to support this statement.

In this context, “larger” refers to the total data size of the MSI dataset (in GB), which reflects a combination of spatial dimensions (number of pixels), spatial resolution, and spectral information per pixel.

In practice, the feasible dataset size is limited by available computational resources (e.g., RAM and GPU memory). For example, in our experiments using systems with up to 320 GB RAM and 64 GB VRAM, methods such as UMAP become increasingly constrained in terms of memory usage and runtime as the dataset size grows.

In contrast, the proposed methods (e.g., PR3D) are designed to scale more efficiently with dataset size, enabling analysis of larger datasets under the same hardware constraints.

4. Regarding **Figure S1**. A major critique of this figure is that there are **not quantitative metrics** to support that the gradient is improved. Within the MSI field, RGB is generally discouraged as it can be misleading for visualization. doi/10.1002/jms.4898

We thank the reviewer for this important comment.

We agree that RGB-based visualizations can be misleading if interpreted as quantitative representations of analyte abundance. In our framework, however, these visualizations are not intended to provide quantitative readouts, but rather to reveal spatial structure and highlight patterns that may not be apparent using conventional methods.

To address the concern about quantitative support, we emphasize that the improvements shown are supported by the benchmarking results presented in the main text, in which we evaluate global structure preservation using established metrics. Figure S1 is intended as a qualitative illustration of these improvements, rather than a standalone quantitative comparison.

The role of these visualizations is to facilitate identification of regions of interest, which can then be further analyzed using quantitative methods (e.g., ROI-based statistical comparisons of m/z features). In this sense, they serve a similar purpose to overview stains in pathology, guiding subsequent, more targeted analysis.

We have clarified this distinction between qualitative visualization and quantitative analysis in the revised manuscript.

5. It would be helpful for Figure S2 to point out what PR3D adds vs k-means specifically. The figure legend mentions plaques. Where would these be in this image, and are they validated?

We thank the reviewer for this helpful comment**, and following this, have revised the figure (now S19).**

For **validation**, we provide a more detailed discussion in other responses, including orthogonal validation using IHC.

K-means is a discrete clustering method that partitions the data into a predefined number of clusters, resulting in sharp boundaries and a limited set of assigned colors. As a result, structures such as amyloid plaques—which share much of their molecular profile with surrounding tissue but differ in a small subset of lipids—are often assigned to the same cluster as neighboring regions and are therefore not clearly distinguished.

In contrast, PR3D provides a continuous visualization, mapping the data into a full RGB space without predefined clusters. This allows more subtle variations in molecular composition to be represented, enabling structures such as plaques to appear as distinct patterns rather than being absorbed into surrounding regions.

**In the revised Figure (now S19),** we have added arrows indicating the location of plaques (cortex layers 4–5 in the APPPS1 mouse brain), to make their identification clearer.

These structures were validated using ROI-based analysis, including known plaque-associated lipids such as GM3 (m/z 1179.7 and M+1) and phosphoinositides (PI, m/z 885.5, M+1, 599.4), supporting their biological interpretation (Figure 4g - m/z list).
