## Supplementary Materials for "Truthful visualizations for mass spectrometry imaging enable high spatial resolution interactive *m/z* mapping and exploration"

www.pahnkelab.eu

**This PDF file includes:**

Figures S1 to S24

Tables S1 to S2

### Supplementary Figures

#### Benchmarking visualizations

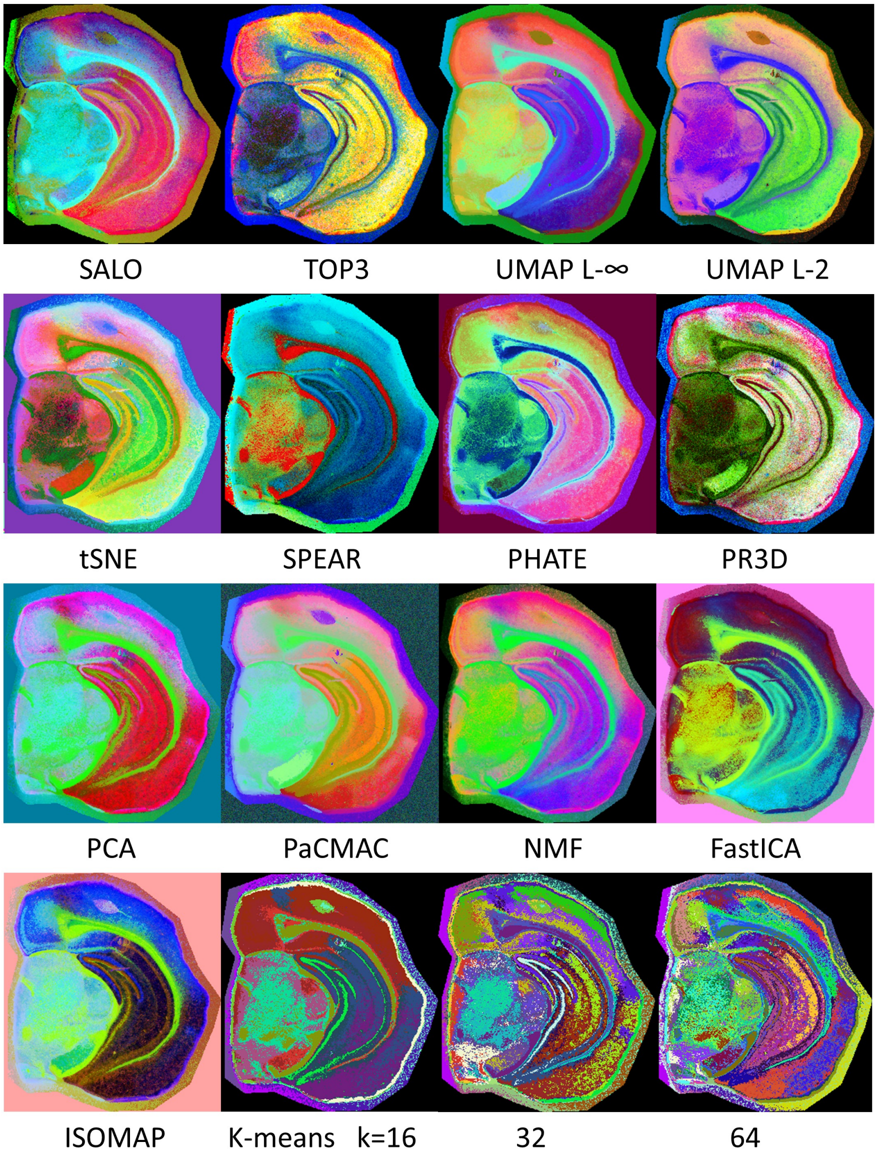


**Figure S1**: **Comparison of benchmarking visualizations and K-means segmentations for one mouse brain hemisphere.** Note the smooth color gradients of the visualizations in the RGB color space versus the strict cluster colors used to highlight the K-means segmentation. K-means is useful for observing anatomical regions; however, inter-cluster anatomical details are invisible because all pixels are assigned the same color.

#### Benchmarking results

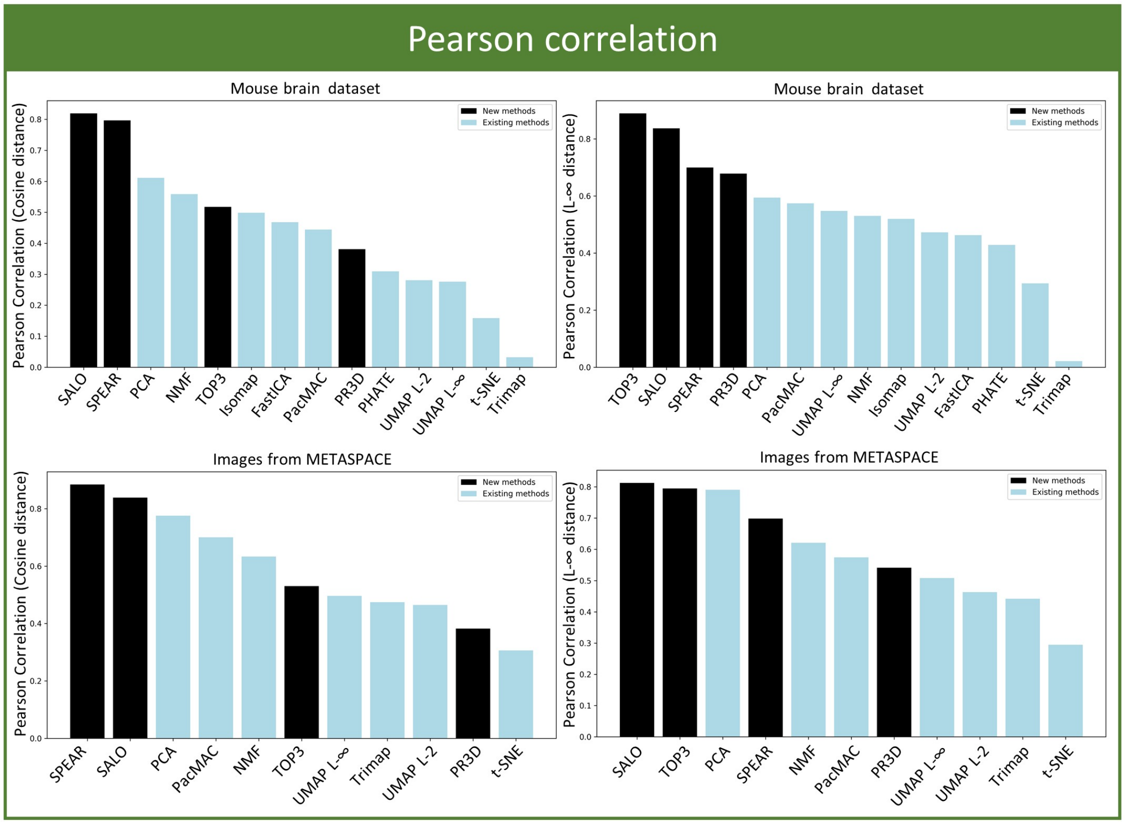


**Figure S2**: **Benchmarking of MSI visualization methods.** Results are sorted from higher (better) to lower. The proposed methods are colored in black (SALO, SPEAR, PR3D, and TOP3). SALO outperforms all existing methods (light blue), whereas SPEAR ranks behind PCA on the METASPACE dataset with the $L_\infty$ distance.

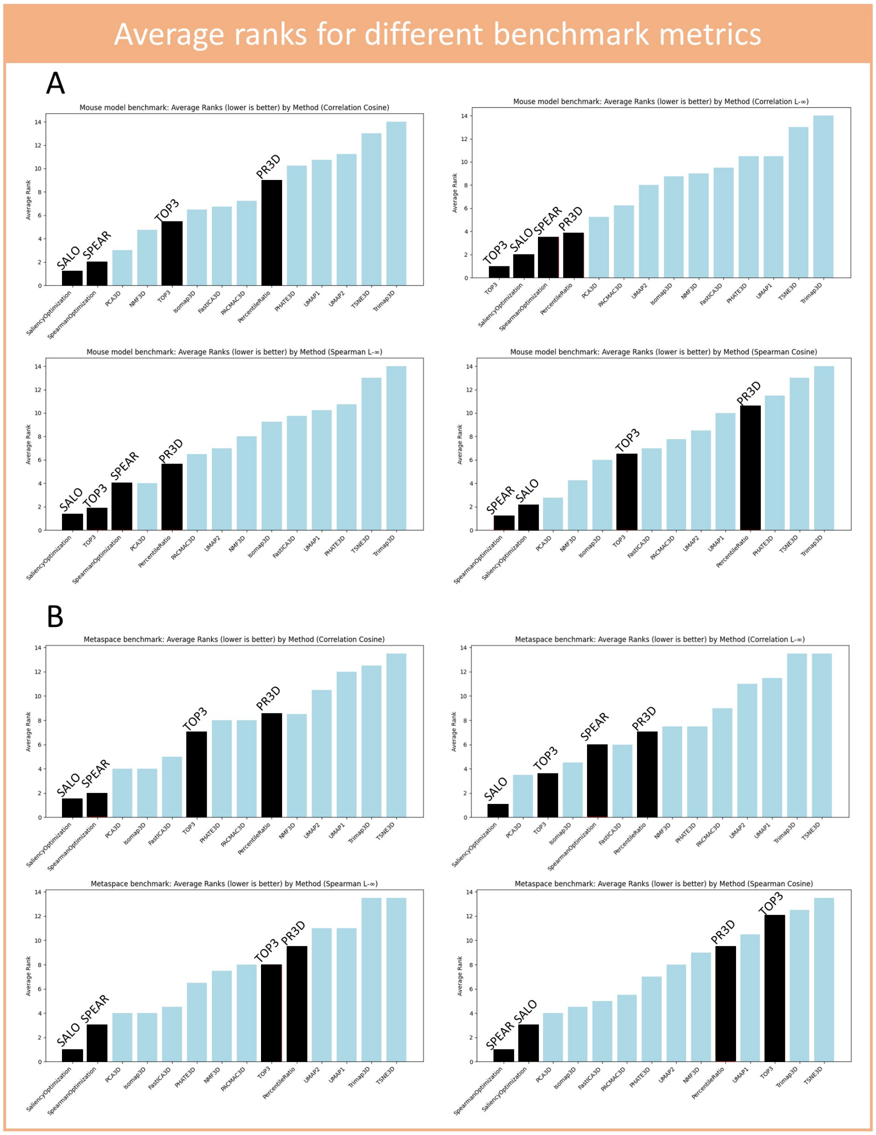


**Figure S3**: **Benchmarking of MSI visualization methods — ranking.** (**A**) Ranking of the visualizations using the mouse brain dataset. (**B**) Ranking of the visualizations using the METASPACE dataset. A lower average rank is better. The proposed methods are in black. SALO outperforms all previously existing methods; the closest competitor is SPEAR (Spearman/cosine metrics, both datasets) or TOP3 ($L_\infty$ distance metric, mouse dataset), depending on the metric and dataset.

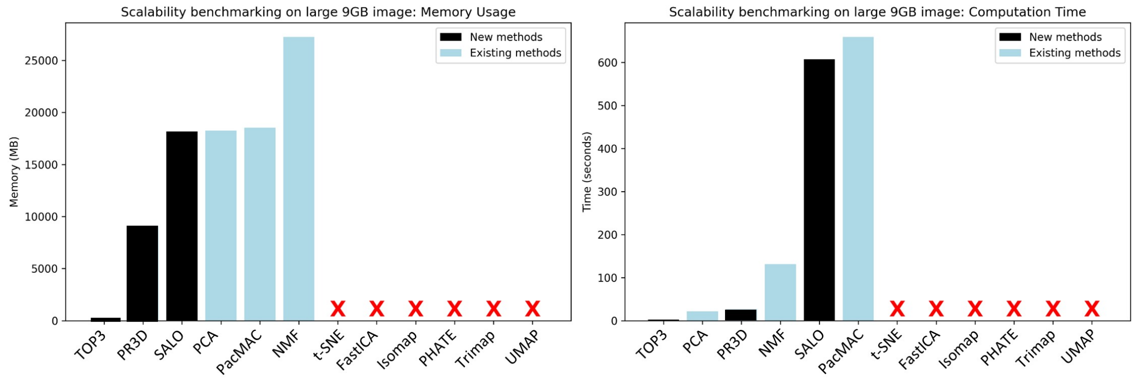


**Figure S4**: **Benchmarking MSI visualization memory and speed on large images.** The proposed new methods are shown as black bars. TOP3 is 600× faster to run and consumes only 2**MB** RAM compared to 25**GB** for NMF and 17**GB** for SALO and PaCMAP. Methods that could not run due to resource needs (>320GB RAM) are marked with ×.

Here, we provide additional benchmarking results using custom metrics that have not previously been used for MSI. We include these for future comparison, as additional evaluation metrics may be valuable, but we do not place them in the main text to avoid confusion about performance on the established MSI metrics. Figure S5 shows results for our custom Saliency metric. Figure S6 shows results for the Spearman correlation metric. SALO is the best-performing method across all datasets. TOP3 and PR3D perform less well on the METASPACE dataset under cosine distance.

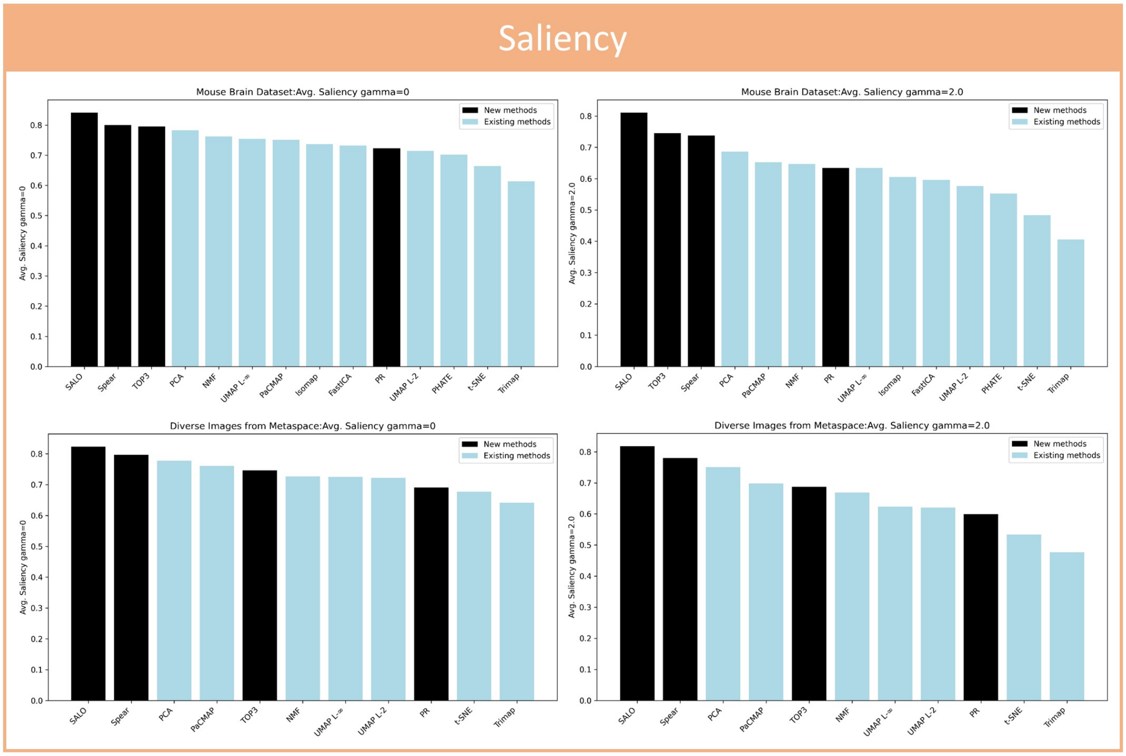


**Figure S5**: **Benchmark of MSI visualizations using the Saliency metric.** Proposed methods are shown in black (SALO, SPEAR, PR3D, TOP3).

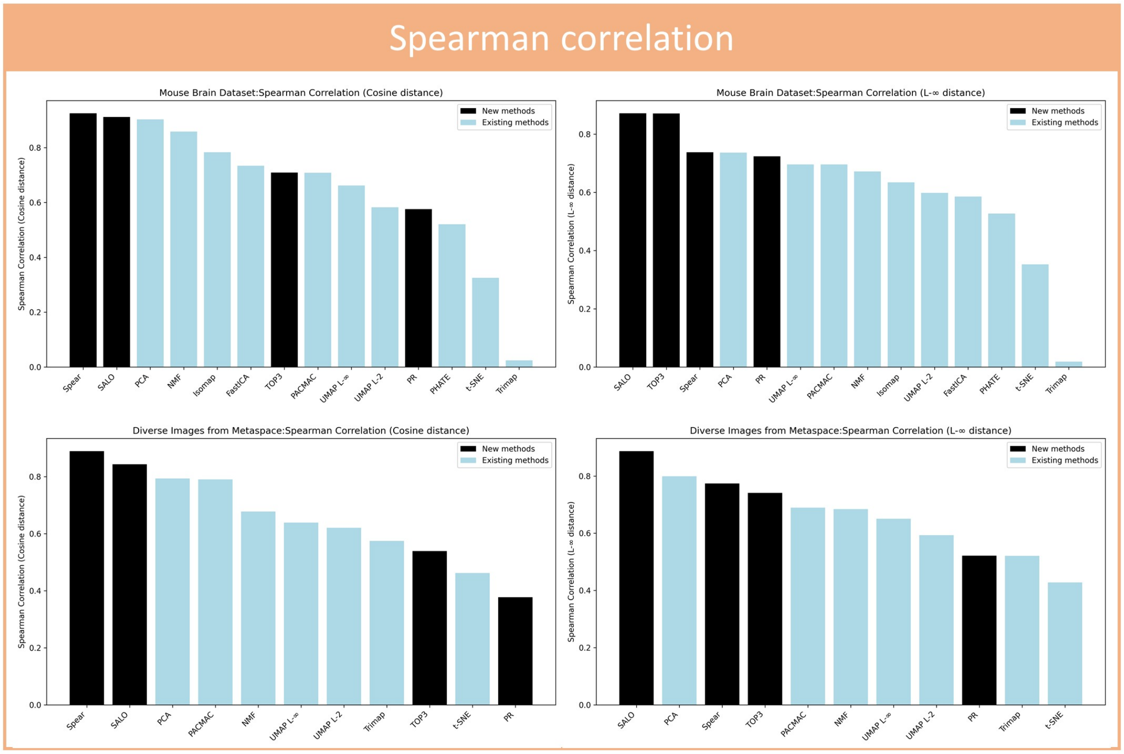


**Figure S6**: **Benchmark of MSI visualizations using Spearman rank correlation.** Proposed methods are shown in black (SALO, SPEAR, PR3D, TOP3).

**Detection of pathological changes of diagnostic relevance.**

In surgical pathology, several histological stains are typically performed, each with a preference for specific tissue properties or pathological changes. We generated multiple visualizations using MSI-VISUAL, highlighting different levels of detail and specificity, similar to immunohistochemical stains, but solely based on lipid MSI data (sample information in Table S2).

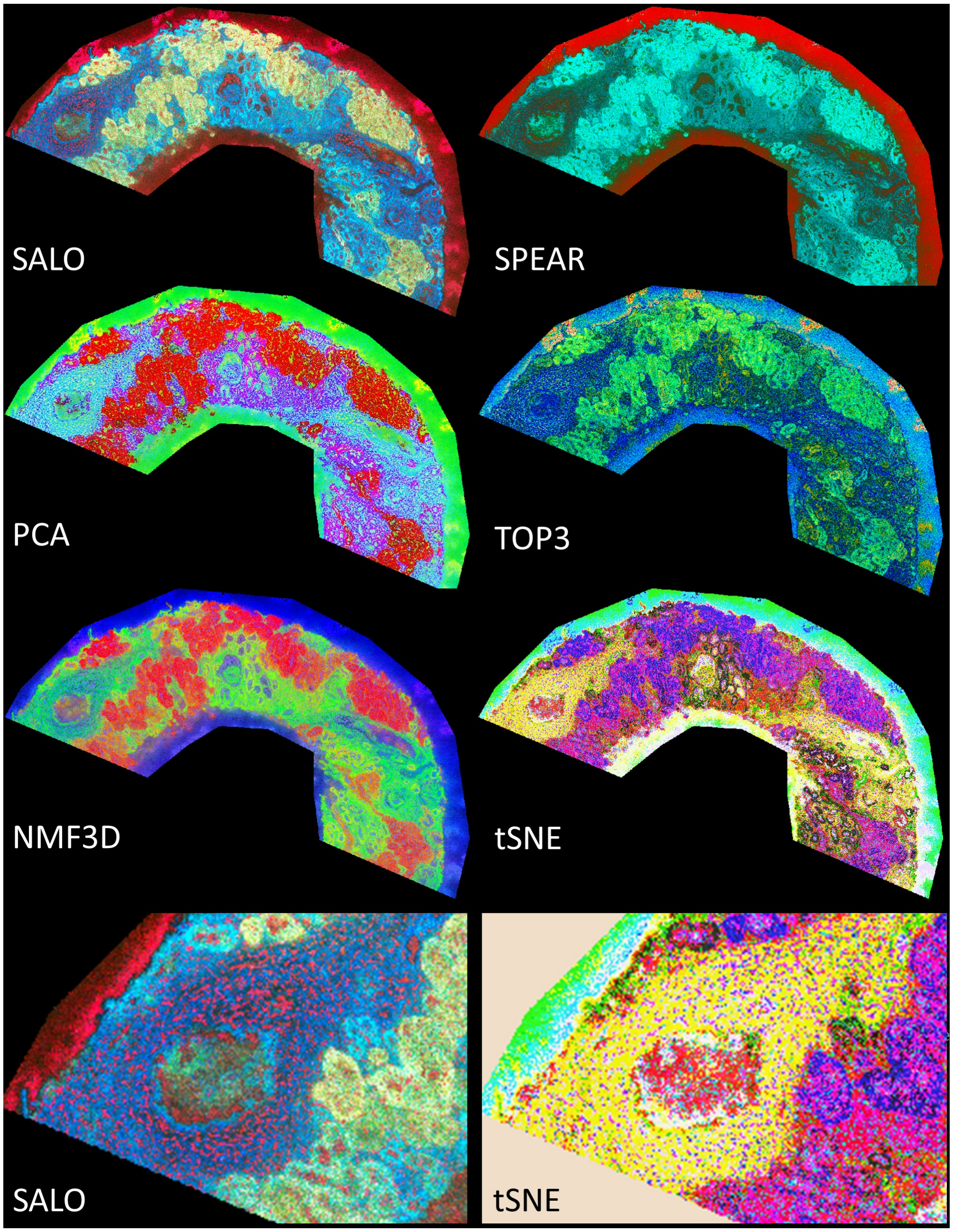


**Figure S7**: **Visualizations of a lipidomics dataset comparing SALO, SPEAR, PCA, TOP3, NMF3D, and t-SNE.** SALO improved the detection of glomerular pathology and of the different sections of the tubular system (proximal and distal tubuli) by generating different interpretable color gradients. The zoomed-in panels highlight a glomerulus, extensive inflammation, and loss of tubular structures (blue in SALO, yellow in t-SNE). Red pixels in SALO and red/violet pixels in t-SNE indicate putative inflammatory infiltrates.

**Detection of functional microanatomy in hippocampus**

Figure S8 shows an overview of UMAP, SALO, SPEAR, PR3D, and TOP3. Figure S9 shows a panel of conventional H&E and immunohistochemical stains, for one hemisphere of our mouse brain dataset (Table S1).

The images show a coronal mouse brain hemisphere section at bregma position –3.08 mm from a C57BL/6J mouse (commonly used B6 genetic background strain), along with a schematic representation of the same location in the Allen mouse brain atlas (*55*). UMAP reveals some gross details in the brain stem, for example thalamic (TH) regional differences and the substantia nigra (SN) (see also 3), and also details in the hippocampal formation (HPF, green region in the Allen atlas), such as dentate gyrus neurons (DG) and, to a lesser extent, Cornu Ammonis field 3 neurons (CA3). In the hippocampal formation (HPF), SALO, SPEAR, and PR3D reveal more details than UMAP and TOP3, with PR3D showing contrasted neuronal bands (CA1, CA3, and DG, all dark pinkish) versus connections between them (layered pinkish or greenish). SPEAR particularly highlights the molecular layer of DG (DG-mo). PR3D highlights cortical layering and differences in myelination extent (CTX, shown by the extent of green signal). TOP3 shows a clear representation of hippocampal neuronal bands (red, CA1, CA3, and DG) against an orange background.

Figure S10 shows a zoomed-in cross-section of hippocampal structures, revealing different levels of detail. UMAP often shows a homogeneous area and misses distinct anatomical features (arrow 1: basal layer of the dentate gyrus; arrow 2: neurons in CA1) that are visible in the new visualizations. Arrow 3 points to single cells detected in the dentate gyrus isthmus using PR3D.

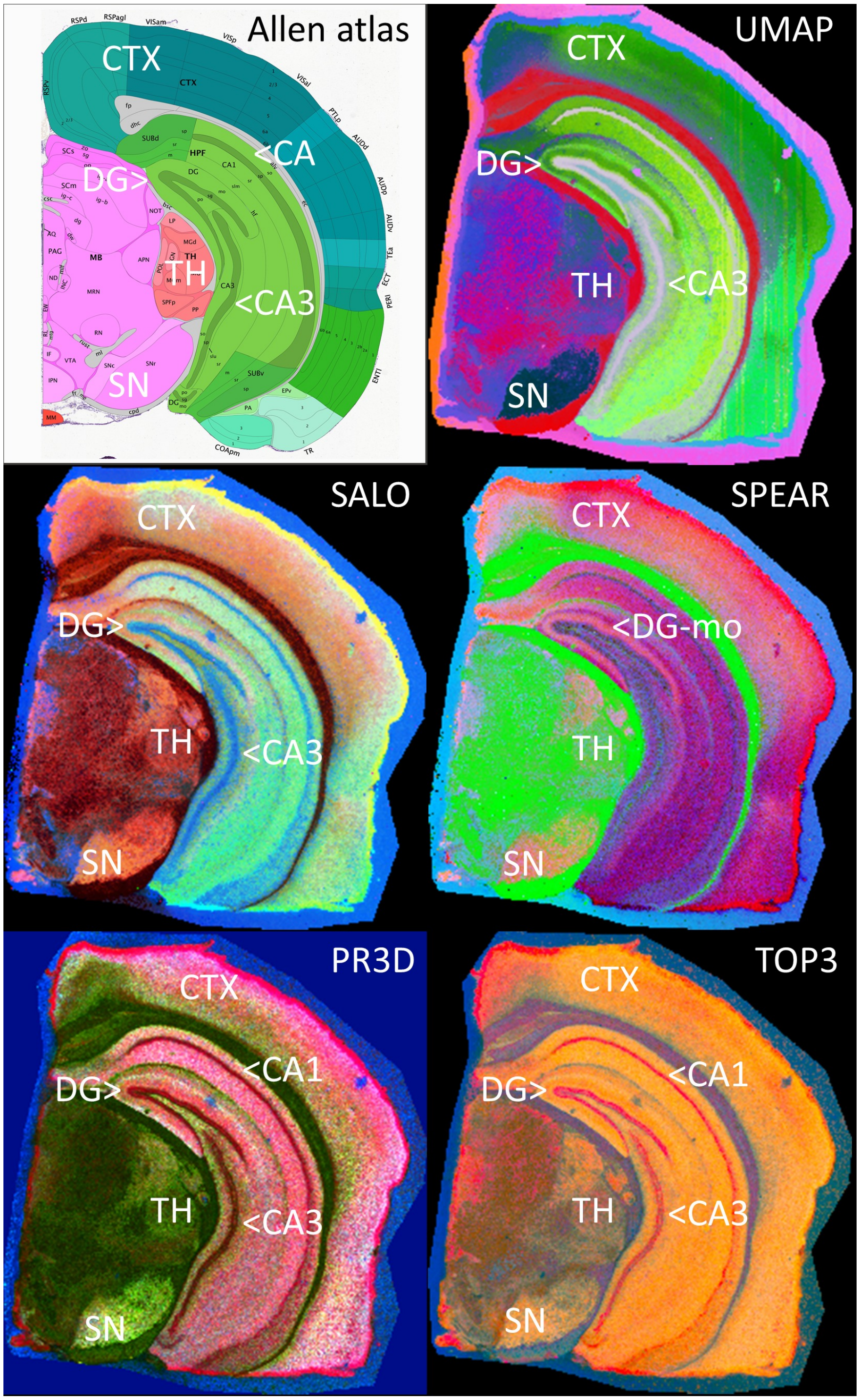


**Figure S8**: **Comparison of UMAP, SALO, SPEAR, PR3D, and TOP3 in mouse brain anatomy.** The visualizations reveal different levels of detail in the thalamus (TH), substantia nigra (SN), hippocampus, and brain stem, consistent with global structure preservation. Legend: CTX, isocortex; CA1, Cornu Ammonis field 1; CA3, Cornu Ammonis field 3; DG, dentate gyrus; TH, thalamus.

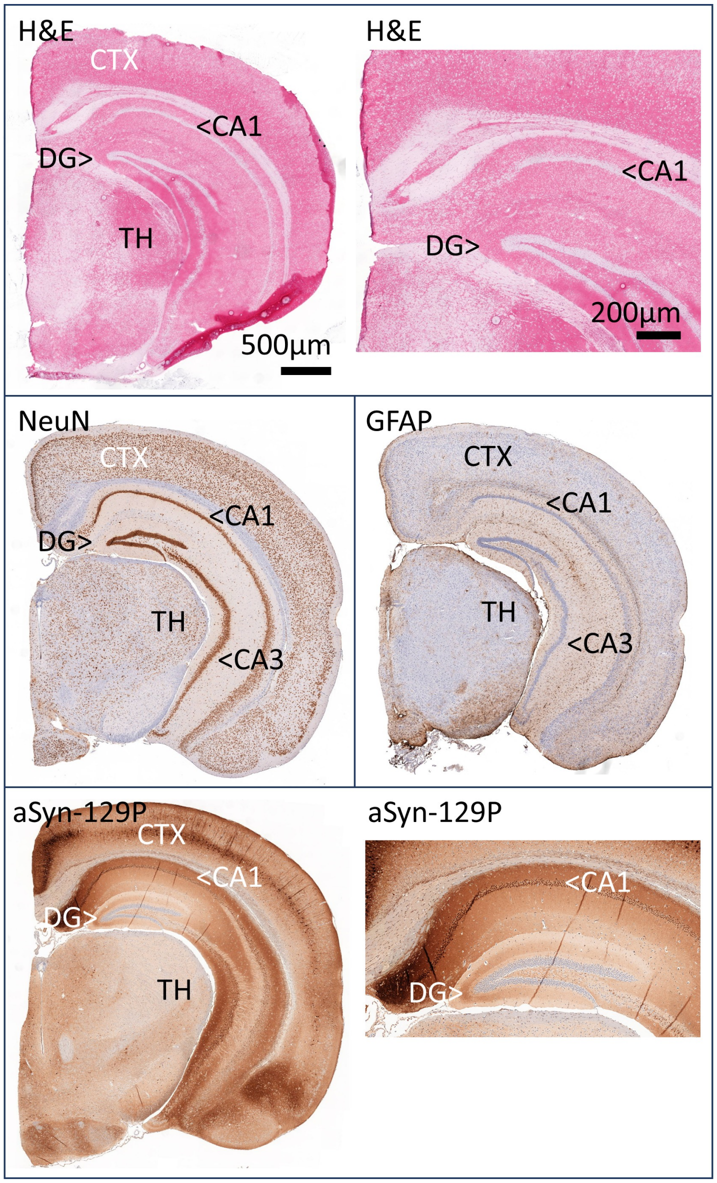


**Figure S9**: **Mouse brain hemispheres stained with H&E and three IHC markers (NeuN, GFAP, $\alpha$Syn-129P).** NeuN for neurons, GFAP for astrocytes, and $\alpha$Syn-129P for neurons and processes showing positivity for phosphorylated $\alpha$-synuclein. The H&E stain magnification of the hippocampal formation is less rich in detail than the MSI visualizations of the lipidomics dataset (Figure S8). The IHC stains show specific details that MSI lipidomics visualizations can reveal in a single dataset, for example hippocampal layering between DG and CA1. Legend: CTX, isocortex; CA1, Cornu Ammonis field 1; CA3, Cornu Ammonis field 3; DG, dentate gyrus; TH, thalamus.

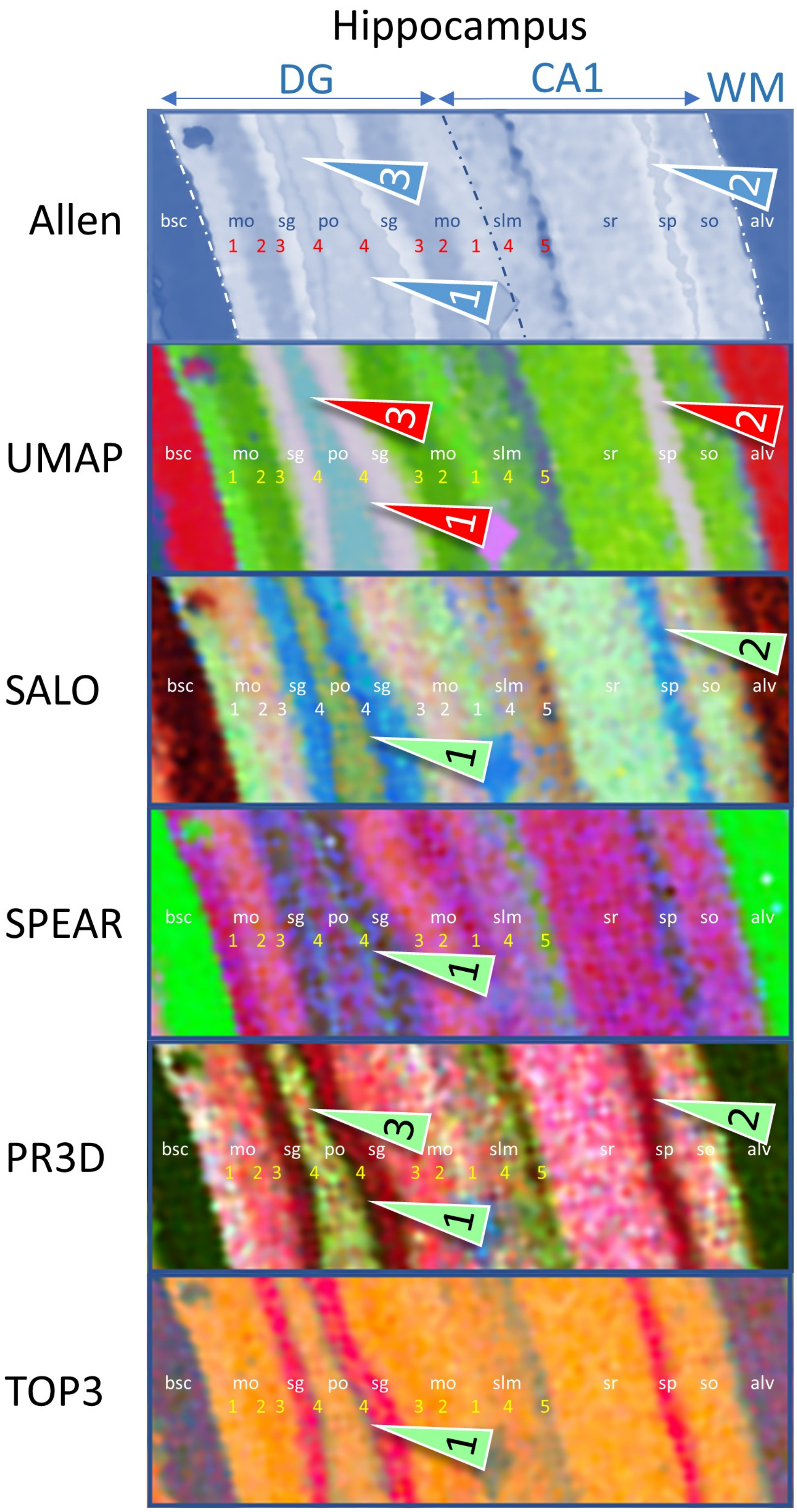

Hippocampus
DG
CA1
WM
Allen
UMAP
SALO
SPEAR
PR3D
TOP3
bsc
mo
sg
po
sg
mo
slm
sr
sp
so
alv
1 2 3 4 4 3 2 1 4 5

**Figure S10**: **Complementary visualizations reveal hippocampal microstructure beyond atlas annotation.** Allen atlas annotations are shown in blue. Red labels (1–5) mark additional subregions visible with MSI-VISUAL but not annotated in the atlas. These regions show distinct lipid signatures. Neuronal-band regions include DG-sg and CA1-sp, whereas DG-mo, DG-po, CA1-slm, CA1-sr, and CA1-so correspond to dendritic and axonal domains. PR3D and SALO resolve most DG–CA1 detail, and SPEAR highlights white-matter (WM) connections, including unannotated myelin-rich areas 4 and 5. Green arrow 1 highlights a connection layer below dentate gyrus neuronal cell bodies (DG-sg). Green arrow 2 highlights the CA1 neuronal cell-body band (CA1-sp). Green arrow 3 highlights dentate gyrus cells in the polymorph layer (DG-po). Legend: red arrows, no detection; CA1, Cornu Ammonis field 1; DG, dentate gyrus; WM, white matter; bsc, brachium of the superior colliculus; DG: mo, molecular layer; sg, granular cell layer (neuron band); po, polymorph layer; CA1: slm, stratum lacunosum-moleculare; sr, stratum radiatum; sp, pyramidal layer (neurons); so, stratum oriens; alv, alveus and external capsule.

**Cross-cohort mapping in public metabolomics/lipidomics datasets.**

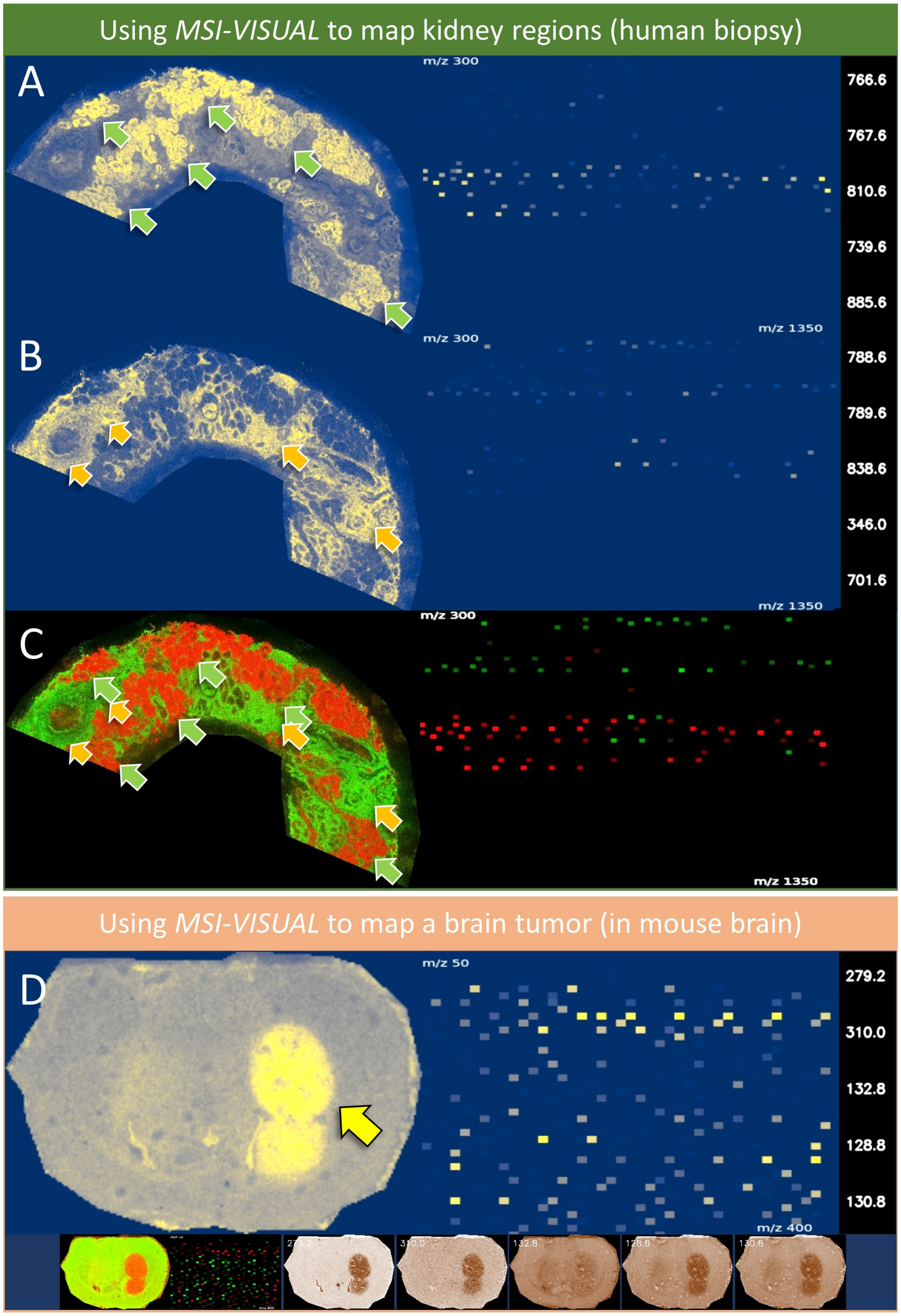

**Figure S11**: **Cross-cohort *m/z* mapping in public METASPACE datasets.** (**A** to **C**) Mapping lipidomics *m/z* values 300–1350 of different regions in a human kidney needle biopsy (Table S2, dataset 4), aggregated images of the 5 most specific *m/z* for (**A**) the kidney tubuli system (green arrows) and (**B**) inflammatory cells (orange arrows) around glomeruli and between the tubuli. (**C**) 2-channel overlay image of tubuli (red) and inflammation (green). The SpecQR nicely presents the separated *m/z* values (green and red squares). (**D**) Mapping metabolomics *m/z* values 50–400 of an implanted brain tumor (yellow arrow) into a mouse brain from (*56*). The SpecQR shows the specific metabolite pattern of the lesion. The lower row of images are virtual pathology stains of the 5 most significant *m/z* values and a differential red-green image of tumor vs other tissue.

**Mapping of brain structures - neurons, connections/white matter, and nuclei**

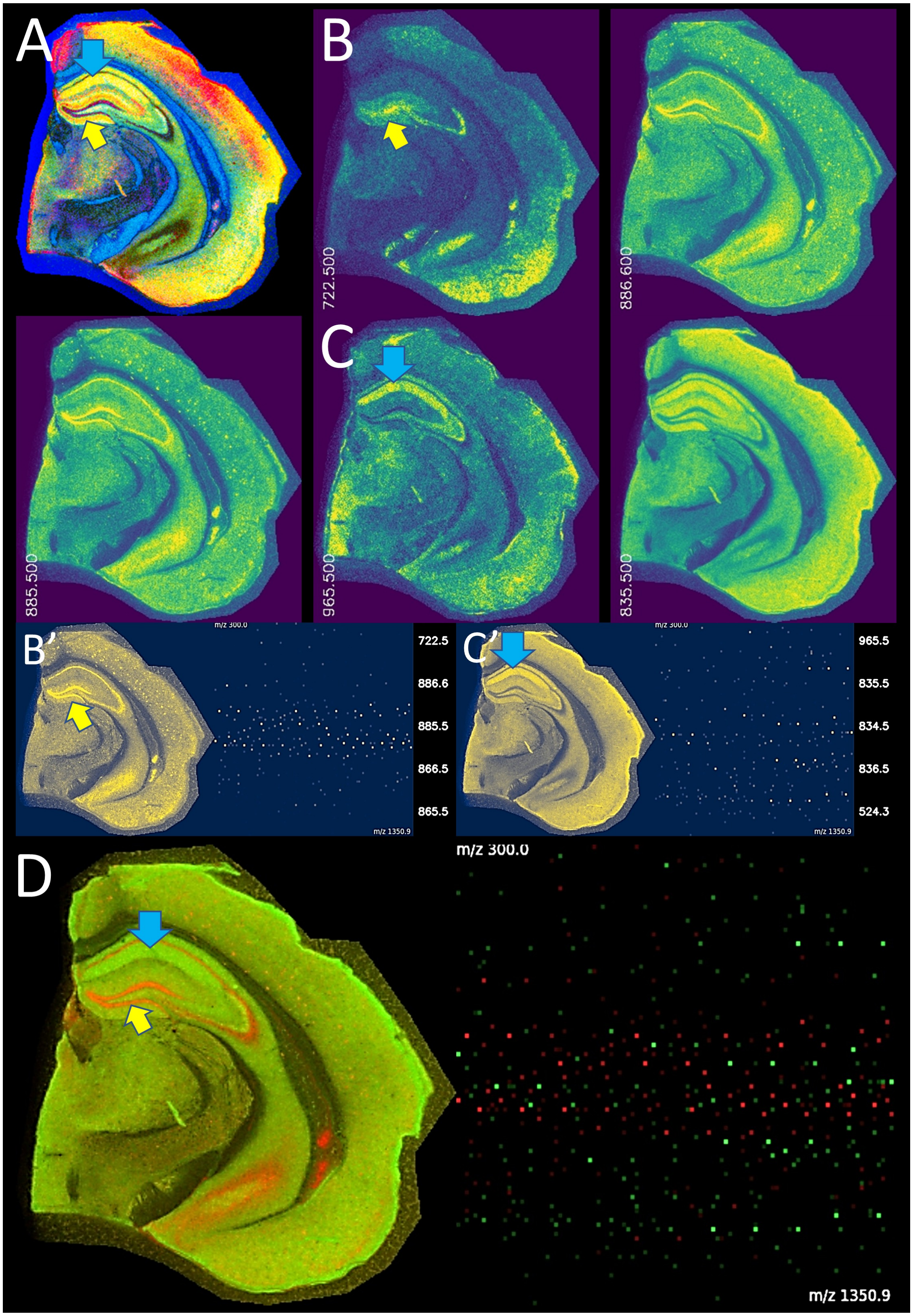

**Figure S12**: **Differential lipid visualization in hippocampal connections.** (**A**) TOP3 visualization of a brain hemisphere. The yellow arrow points to the dentate gyrus neurons (DG granule cell layer, dark band) and the blue arrow towards axonal processes of the Cornu Ammonis neurons (CA1 stratum radiale, yellow band). These were selected for further analysis. (**B**) shows the three best *m/z* ion images (722.5, 885.5, 886.6) corresponding to the selected location for detection (yellow arrow). (**B'**) shows the aggregated ion image of the 5 best *m/z* values (list on the right) and the SpecQR graph. (**C**) shows the two best *m/z* ion images (965.5, 835.5) and the selected CA1 stratum radiale (blue arrow). (**C'**) shows the aggregated ion image of the 5 best *m/z* values (list on the right) and the SpecQR graph. (**D**) shows a red–green overlay image of both aggregated ion images from the selected regions and the SpecQR in the corresponding colors. The SpecQR diagram shows the *m/z* distribution from *m/z* 300–1350 with very distinct, non-overlapping *m/z* values for both structures. Note the detection of A$\beta$ plaques in (**B'**) and (**D**) (red channel) labeled with *m/z* 722.5 and 885.5/886.6 (M+1).

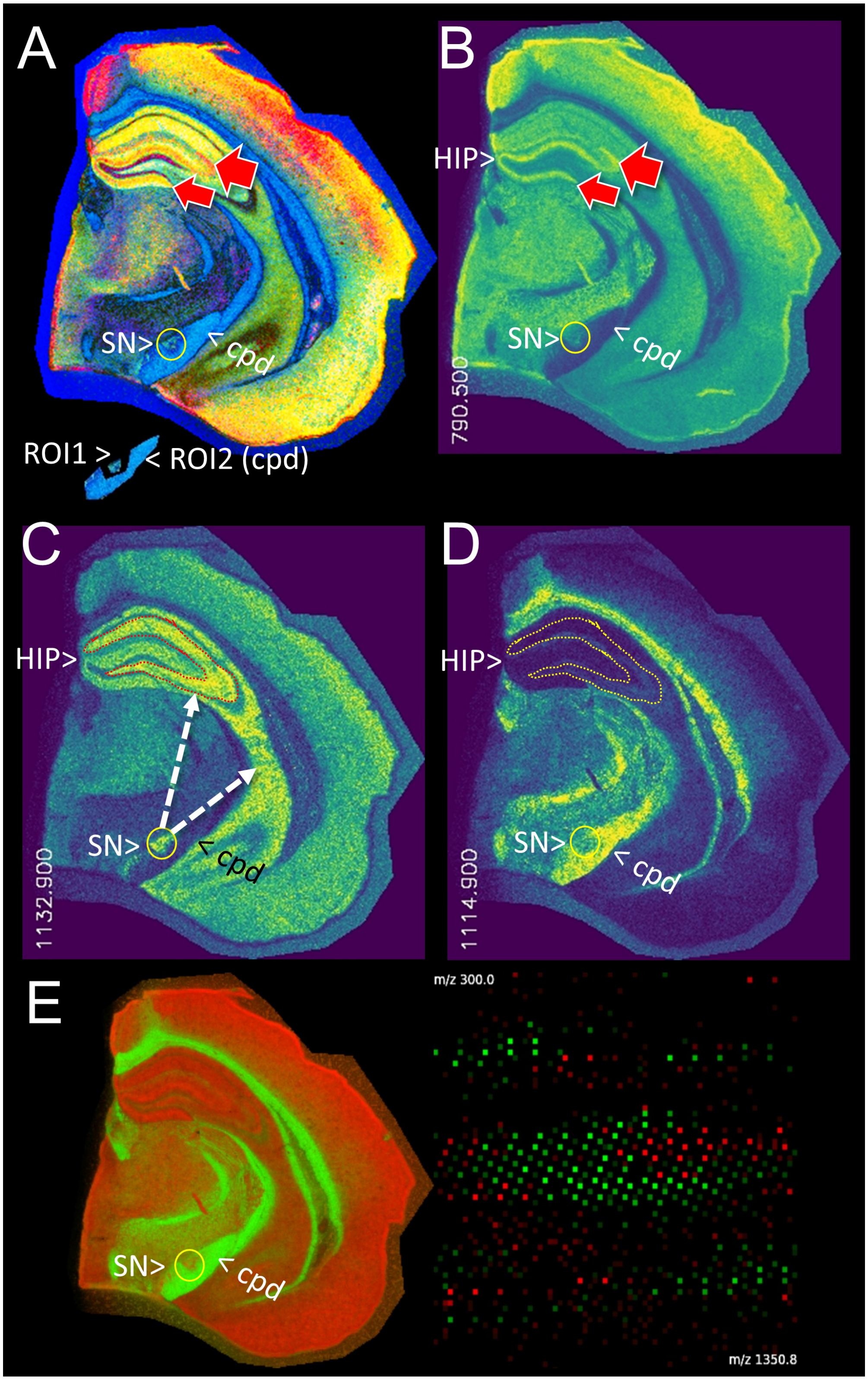
A
SN>
< cpd
ROI1 >
< ROI2 (cpd)
B
HIP>
SN>
< cpd
790.500
C
HIP>
SN>
< cpd
1132.900
D
HIP>
SN>
< cpd
1114.900
E
SN>
< cpd
m/z 300.0
m/z 1350.8

**Figure S13**: **Region-specific *m/z* signatures separate nuclei from white-matter tracts.** (**A**) TOP3 visualization reveals detailed brain structures, including ROI1 (Substantia nigra, SN) and ROI2 (cerebral peduncle, cpd). (**B**, **C**) ROI1 (SN) was identified as a neuronal region, with *m/z* values corresponding to other high-density neuronal regions such as the hippocampus (HIP). (**D**) ROI2 (cpd) was correctly identified as a white matter tract, with no overlap with the SN. (**E**) Overlay of the top five *m/z* values from ROI1 (red) and ROI2 (green), with SpecQR graph showing the *m/z* distribution from 300–1350, confirming distinct *m/z* profiles of gray substance (red) and white matter tracts (light green).

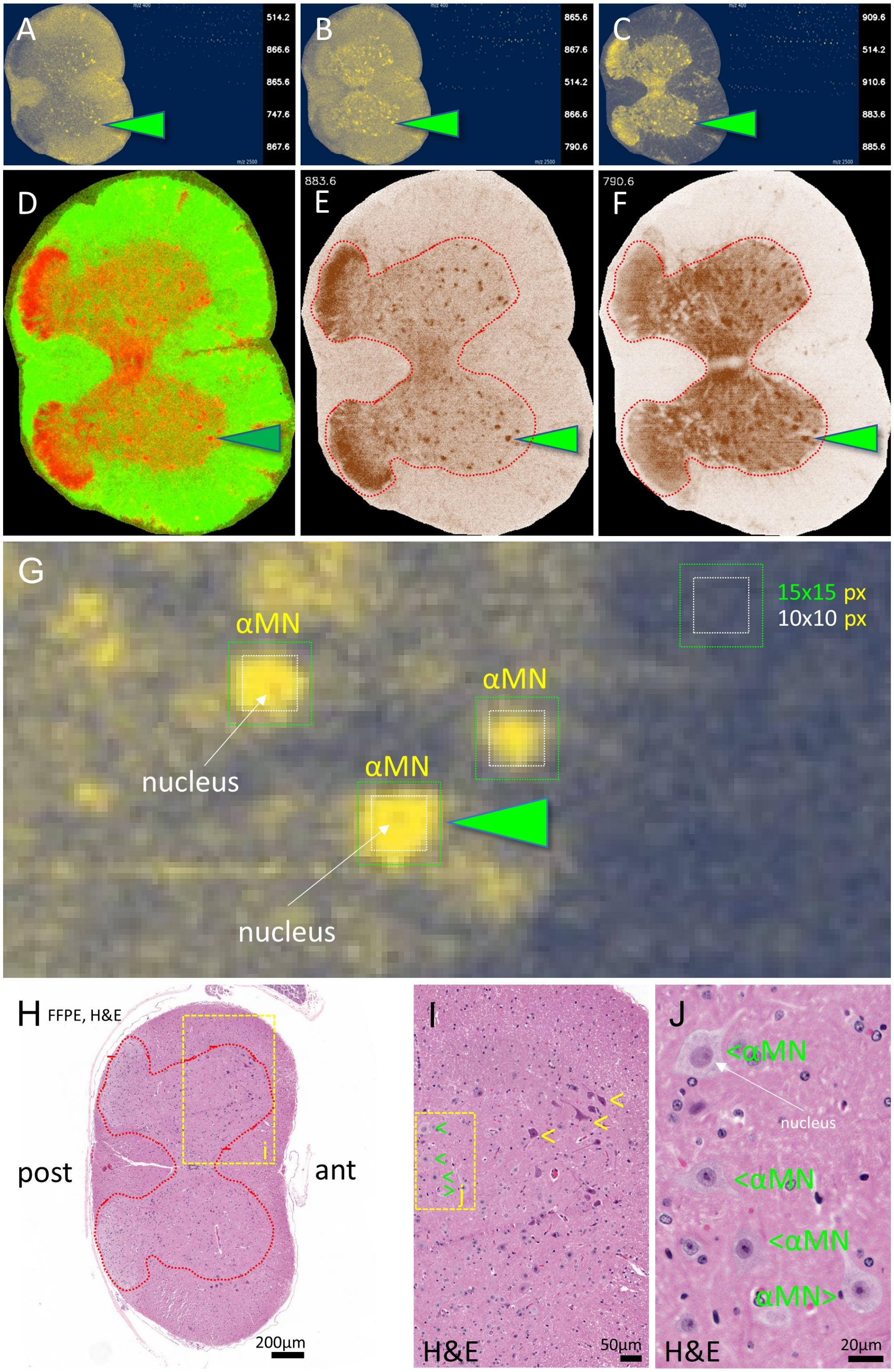
A
B
C
D
E
F
883.6
790.6
G
15x15 px
10x10 px
αMN
αMN
αMN
nucleus
nucleus
H FFPE, H&E
post
ant
200µm
I
H&E
50µm
J
<αMN
nucleus
<αMN
<αMN
αMN>
H&E
20µm

**Figure S14**: **Single-cell detection of spinal $\alpha$ motor neurons (5 $\mu m$/pixel).** (**A** to **C**) show three independent rounds of interactive ROI selection for putative $\alpha$MNs, with corresponding region-specific *m/z* values (right) and SpecQR graphs (middle). MSI was performed on fresh-frozen tissue (METASPACE dataset; Table S2, dataset 5). (**C**) shows the best-performing pixel-wise ROI selection, with the top five *m/z* values displayed as an aggregated ion image. Two significant *m/z* values correspond to phosphoinositols, PI(18:0/20:4) at 883.548 Da and PI(18:0/22:6) at 885.564 Da, recently reported in $\alpha$MNs (*42*). (**D**) shows a red–green overlay highlighting $\alpha$MNs (the second motor-neuron population affected in amyotrophic lateral sclerosis, ALS) and their posterior-horn connections (red channel). Surrounding myelinated white-matter tracts are shown in light green. (**E**, **F**) show two Virtual Pathology Stains (VPS) for selected *m/z* values (883.6 and 790.6). The red dotted line delineates spinal-cord gray matter and the central aqueduct. (**G**) shows a zoomed view of panel (**C**) to assess pixel coverage of $\alpha$MNs. Green arrows indicate an example spanning $>$ 100 pixels. Confident single-cell assessment in MSI is feasible when at least 3×3 pixels cover the structure. (**H**) shows a routine H&E overview of a mouse spinal-cord section from formalin-fixed, paraffin-embedded (FFPE) tissue. FFPE sections are slightly shrunken due to pretreatment (dehydration, delipidation, paraffinization). The MSI sections in (**A** to **G**) were prepared from fresh-frozen tissue. (**I**) shows magnification of the anterior horn highlighting $\alpha$MNs. Green arrows indicate healthy neurons; yellow arrows indicate $\alpha$MNs with harvesting-related artifacts (condensed cytoplasm and shrunken cell body). (**J**) shows high magnification of four $\alpha$MNs. The white arrow points to the nucleus. Abbreviations: ant, anterior; post, posterior.

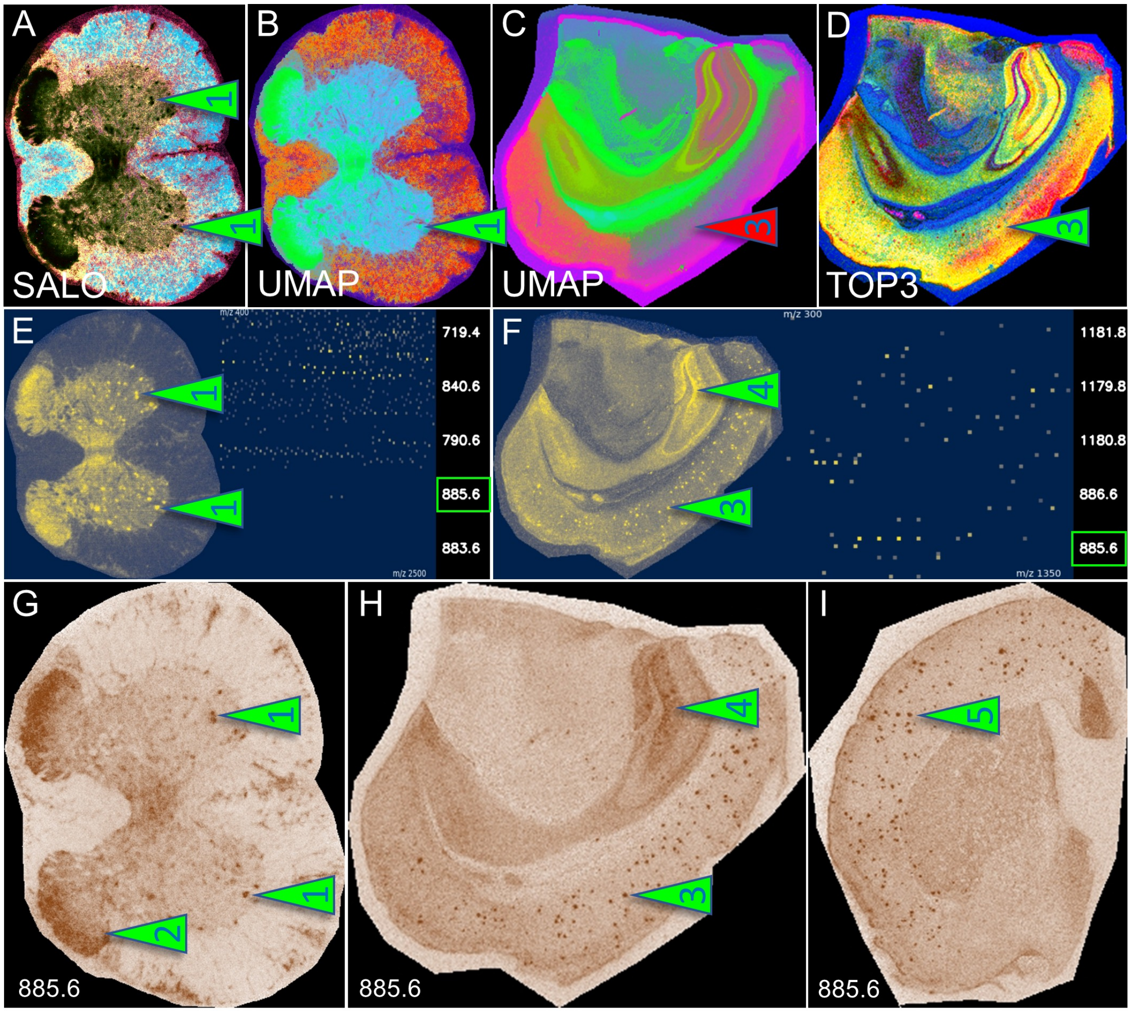


**Figure S15**: **Mapped *m/z* values reveal shared lipid signatures in spinal neurons and cortical A$\beta$ plaques**. (**A** to **D**) Visualizations of spinal neurons (arrows 1) and A$\beta$ plaques (arrows 3). (**C**) UMAP does not highlight plaques (red arrow 3). (**D**) TOP3 successfully detects plaques. (**E** to **F**) the aggregated ion image and SpecQR diagrams show *m/z* values shared between spinal neurons and cortical A$\beta$ plaques (green box). (**G** to **I**) Virtual Pathology Stain images of *m/z* 885.6 - PI(18:0/22:6) - in the spinal cord neurons (*42*), and brains of two AD mice. (**G**) spinal cord anterior horn $\alpha$ motor neurons (arrow 1) and hind horn connections (arrow 2). (**H**) hippocampal connections (Dentate gyrus' molecular layer, arrow 4). (**H**, **I**) cortical A$\beta$ plaques (arrow 3 and 5).

**Histopathological 'ground truth' — the H&E stain**

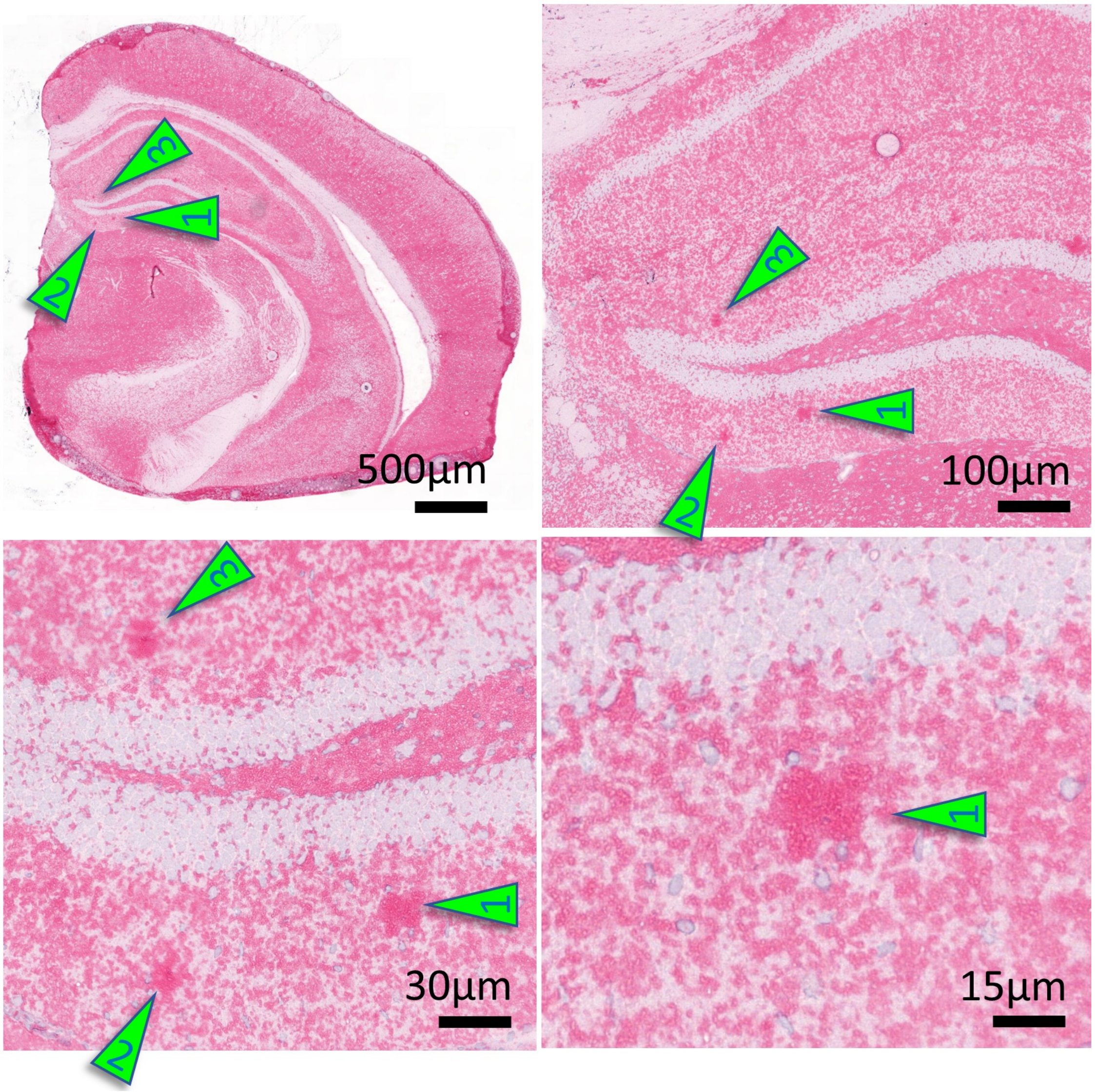


**Figure S16**: **H&E overview of the APPtg-positive mouse hemisphere shown in Figure 4, with numerous cortical and hippocampal plaques.** The arrows indicate condensed eosinophilic A$\beta$ protein aggregates in the hippocampal formation at different magnifications. Arrow 1 corresponds to the plaque highlighted with a red arrow in the Results section (Figure 4). The plaque core diameter is estimated to be approximately 20 µm.

#### Histopathological 'ground truth' for amyloid plaques — the anti-A$\beta$ stain

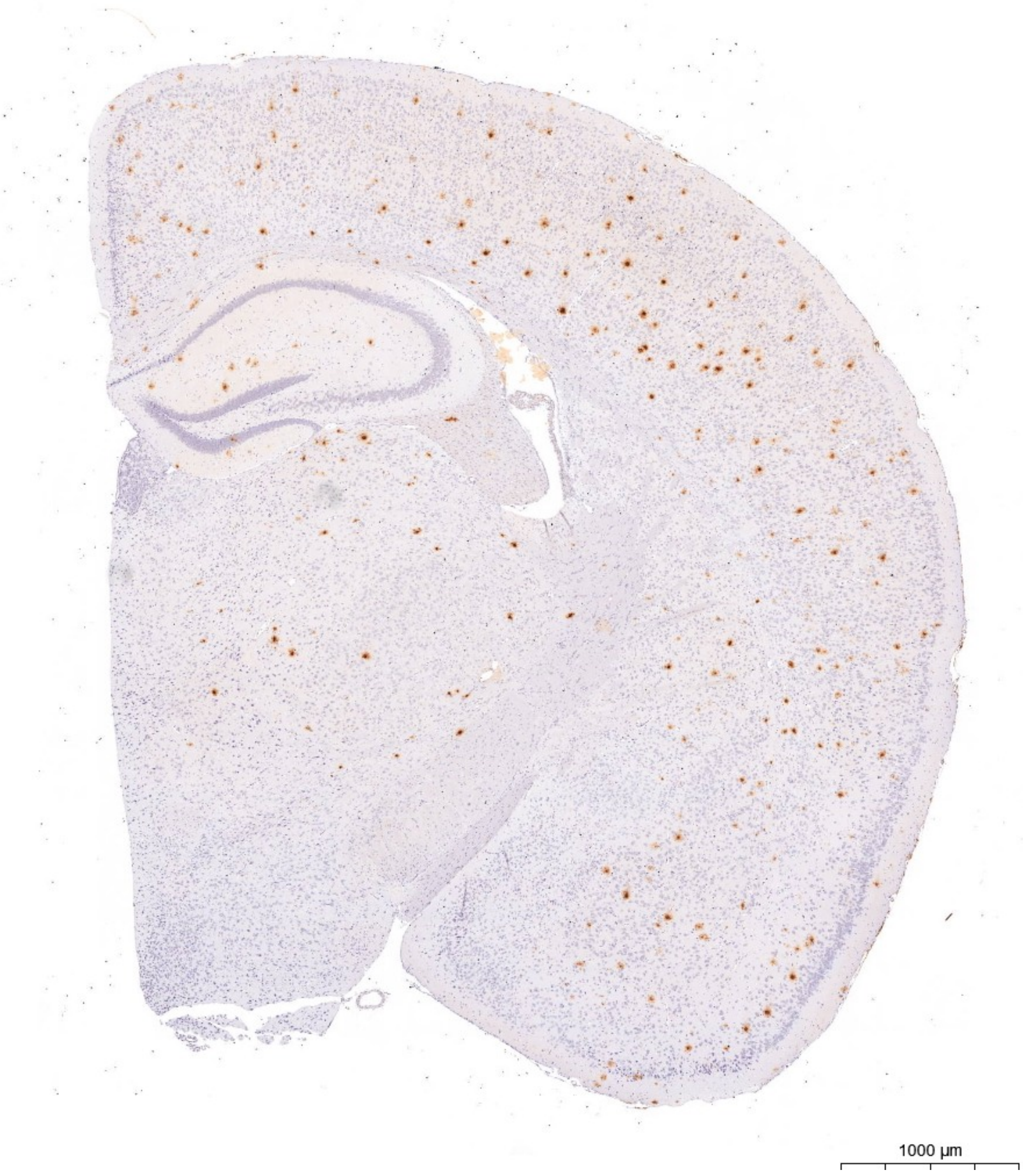


**Figure S17**: **Immunohistochemical stain of a mouse brain hemisphere for A$\beta$.** Here, we show amyloid deposits in a APPtg-positive mouse with numerous cortical and hippocampal plaques using antibody clone 4G8 as described in (*32, 33, 36, 57, 58*).

#### The SpecQR diagram

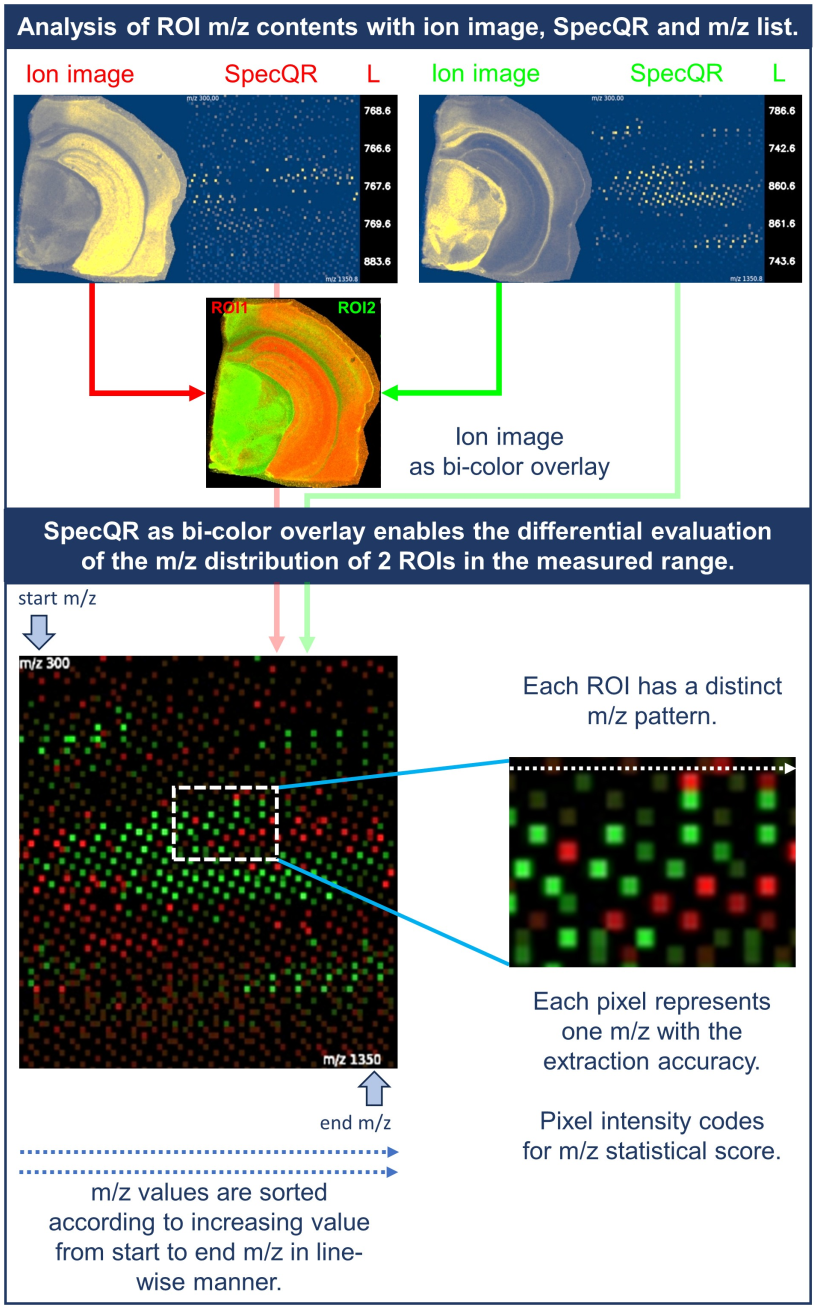


**Figure S18**: **Evaluation of ROIs using the SpecQR diagram.** SpecQR enables rapid comparison of region-specific molecular signatures by summarizing top-scoring ions, showing score-weighted *m/z* values across the acquired range, and providing a red/green overlay of region-enriched signals.

**Comparison of PR3D and K-means segmentation**

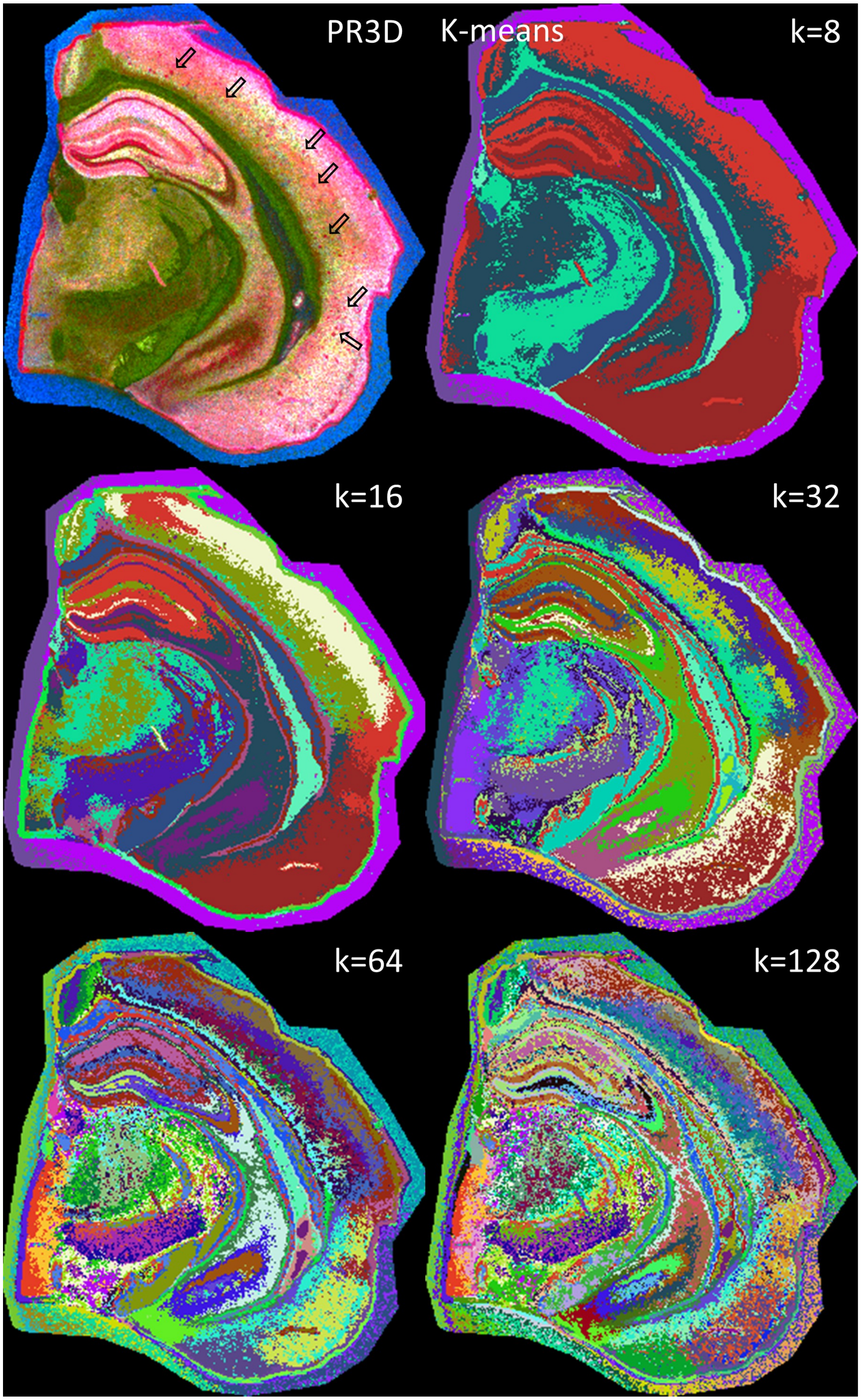

**Figure S19**: **K-means loses intra-cluster anatomical detail and misses small structures compared with continuous-value RGB visualization methods.** The new PR3D visualization is shown alongside five K-means segmentation examples ($k = 8 - 128$) from the same mouse brain dataset. While K-means is effective for identifying and distinguishing broad anatomical regions, it often fails to capture finer structures, such as amyloid plaques (arrows), which are not assigned to their own distinct cluster. This limitation arises because K-means assigns the same color to all pixels within a cluster, resulting in loss of intra-cluster anatomical detail relative to the continuous representation provided by PR3D. Increasing the number of clusters (higher k) can reveal finer details, but often at the cost of interpretability and usability, making precise region annotation more difficult. Although K-means remains a practical and widely used tool in MSI analysis, we advocate broader adoption of continuous-value, dimensionality-reduction-based visualization methods to complement clustering and improve anatomical and molecular interpretability.

**Binning effects on advanced and lightweight visualizations**

Here, we present two panels to show the effects of data binning before dimensionality reduction:

- a hemisphere of our mouse brain dataset with 20µm pixel resolution (Table S1)
- a dataset from METASPACE (Table S2, dataset 5) of a mouse spinal cord with 5µm pixel resolution (Figure S14)

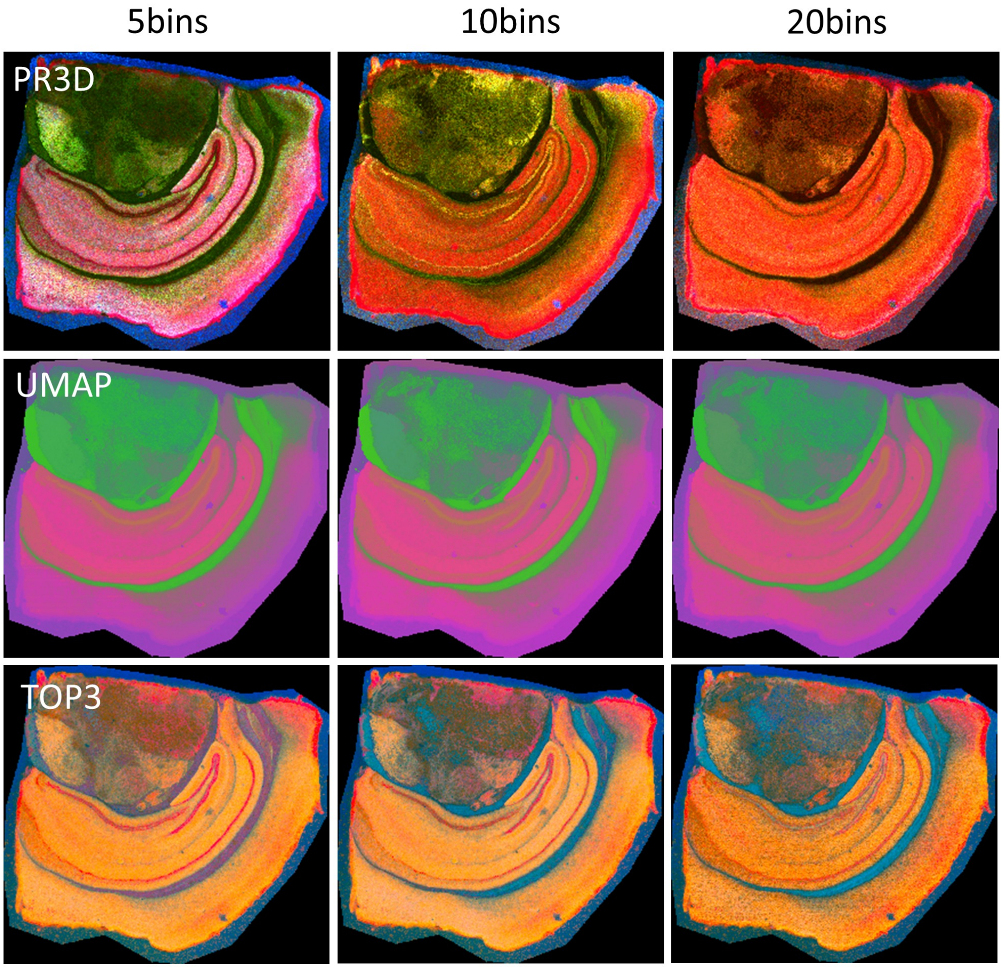

**Figure S20**: **Overview of the effects of different binning for the visualizations of hippocampal details.** Binning with 5, 10, and 20 bins per Dalton (respectively 0.2 = 200 mDa, 0.1 = 100 mDa, and 0.05 = 50 mDa *m/z* value precision) shows slight effects for the lightweight visualization methods PR3D and TOP3, with 5 bins highlighting optimal visualization quality in the hippocampus. UMAP, as a more complex DR method, is only marginally affected by the different binning ranges.

To compare binning effects on the spinal cord dataset visualizations, we used an extensive panel of the following methods:

1. K-means: k=8, max-iter:200, color-scheme:gist-rainbow
2. K-means: k=16, max-iter:200, color-scheme:gist-rainbow
3. K-means: k=32, max-iter:200, color-scheme:gist-rainbow
4. K-means: k=64, max-iter:200, color-scheme:gist-rainbow
5. Non-parametric UMAP: min-dist: 0.5, n-neighbors: 100, metric: euclidean
6. Non-parametric UMAP: min-dist: 0.5, n-neighbors: 500, metric: euclidean
7. NMF3D: max-iter=200
8. NMF3D: max-iter=1000
9. NMF clustering: k=8, max-iter=2000, spatial-norm
10. NMF clustering: k=16, max-iter=2000, spatial-norm
11. NMF clustering: k=32, max-iter=2000, spatial-norm
12. PR3D: static percentiles
13. SALO: epochs=200, reg.=0.001, sampling=coreset, points=1000
14. SALO: epochs=200, reg.=0.01, sampling=coreset, points=1000
15. SPEAR: epochs=200, reg.=0.001, sampling=coreset, points=1000
16. TOP3: low=99.9

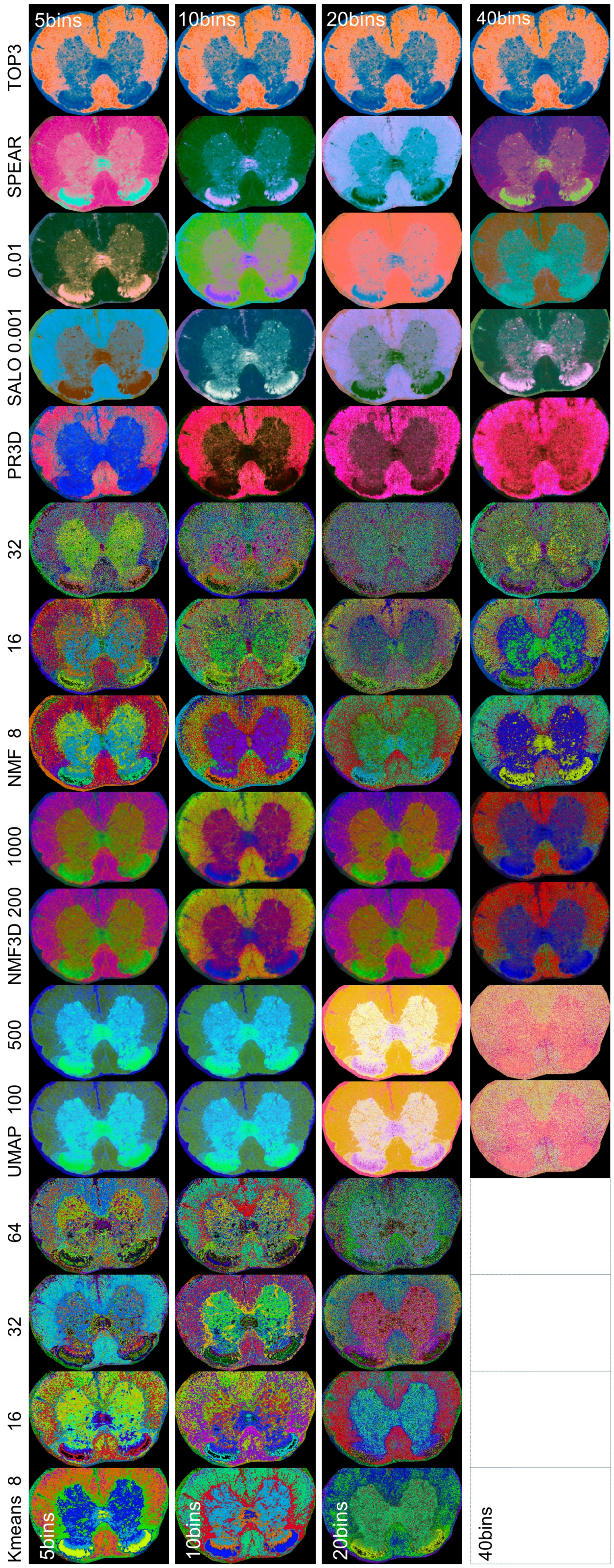
5bins
10bins
20bins
40bins
TOP3
SPEAR
0.01
SALO 0.001
PR3D
32
16
NMF 8
1000
NMF3D 200
500
UMAP 100
64
32
16
Kmeans 8
5bins
10bins
20bins
40bins

**Figure S21**: **Overview of binning effects on advanced and lightweight visualizations.** We used a 5 µm-resolution mouse spinal cord dataset and applied binning with 5 (12 GB .npy file size), 10 (24 GB), 20 (48 GB), and 40 (96 GB) bins per Dalton, corresponding to precisions of 0.2 = 200 mDa, 0.1 = 100 mDa, 0.05 = 50 mDa, and 0.025 = 25 mDa. High spectral resolution (i.e., a high number of bins) requires substantial computational and memory resources. Binning effects strongly depend on the selected visualization or dimensionality-reduction method:
The lightweight TOP3 visualization is not affected by binning. PR3D performs best with 5 bins. Our advanced visualization methods, SALO and SPEAR, are robust to binning. NMF3D shows only minor differences across binning settings. In contrast, UMAP shows substantial noise at 40 bins. Clustering methods (K-means, NMF) are strongly affected by binning; in this example, 5 bins produced the best output. For K-means and NMF, visualization quality decreases as the number of bins increases. Visualizations left blank in the 40-bins column could not be computed because of resource limitations. We used 320 GB RAM and 64 GB GPU memory.
In summary, for truthful visualization of large datasets, binning with 5 bins provides the best balance of quality and computational feasibility.

**PR3D percentile-parameter sensitivity**

Here, we show how different fixed percentile settings in PR3D produce markedly different visualizations of the same MSI image. This sensitivity helps explain why PR3D is less robust on the diverse METASPACE benchmark when one static parameter set is used across datasets. Going forward, PR3D will require automatic parameter tuning.

However, when parameters are well chosen, PR3D can reveal strong anatomical detail, e.g., tuned PR3D for mouse brain in Figures S8 and S19.

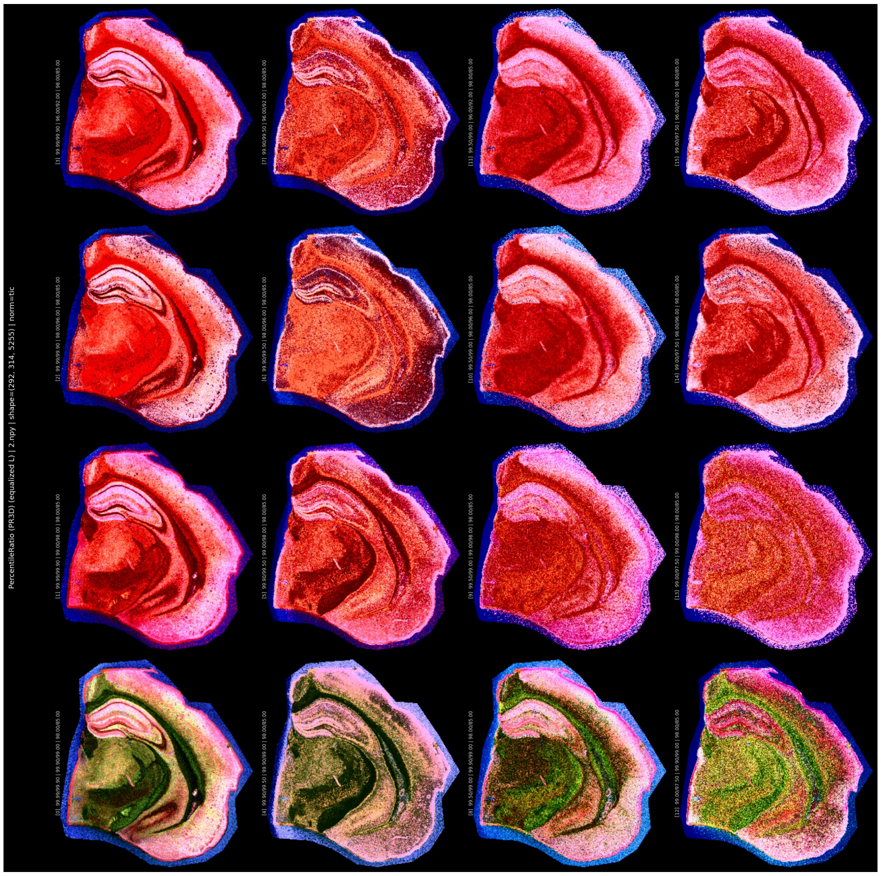

**Figure S22**: **PR3D visualizations across fixed percentile settings of one of our brain hemispheres.** Each panel shows the same MSI dataset rendered with a different percentile combination. The grid demonstrates that PR3D output depends strongly on percentile choice: some settings suppress tissue structure, whereas others enhance fine anatomical detail and lesion contrast. This sensitivity is a key reason for future automatic parameter tuning.

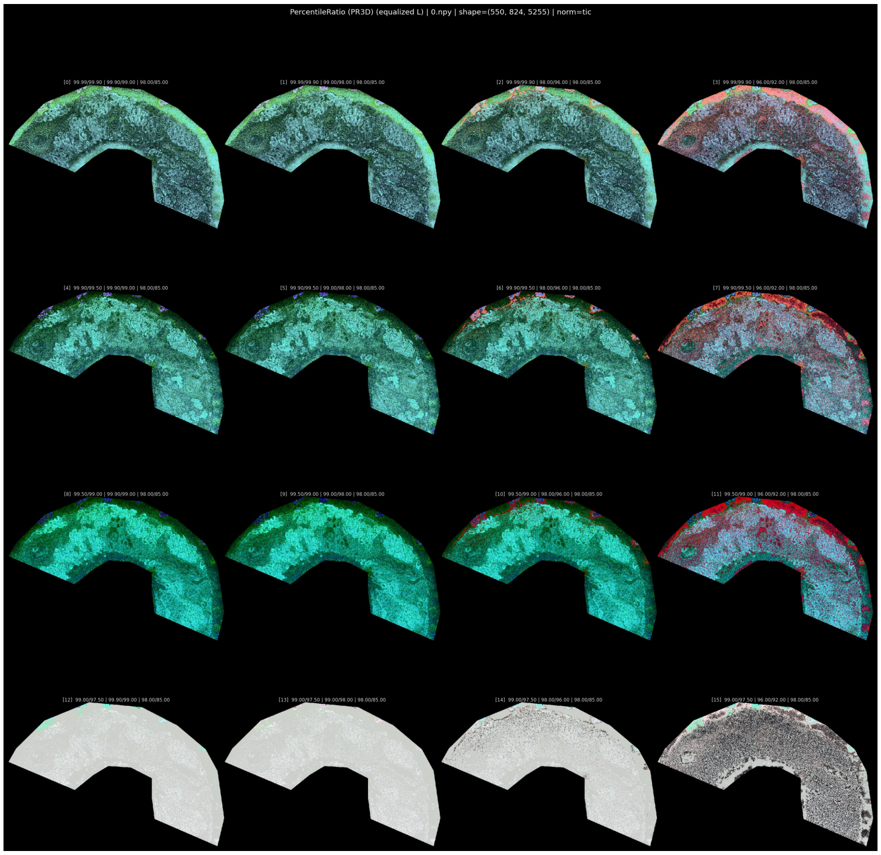


**Figure S23**: **PR3D visualizations across fixed percentile settings in a METASPACE dataset.** Each panel shows the same MSI dataset rendered with a different percentile combination. The grid demonstrates that PR3D output depends strongly on percentile choice: some settings suppress tissue structure, whereas others enhance fine anatomical detail and lesion contrast. This sensitivity is a key reason PR3D is less robust across diverse benchmarks when static settings are used, and it motivates future automatic parameter tuning.

#### Reference point selection and visualization metrics

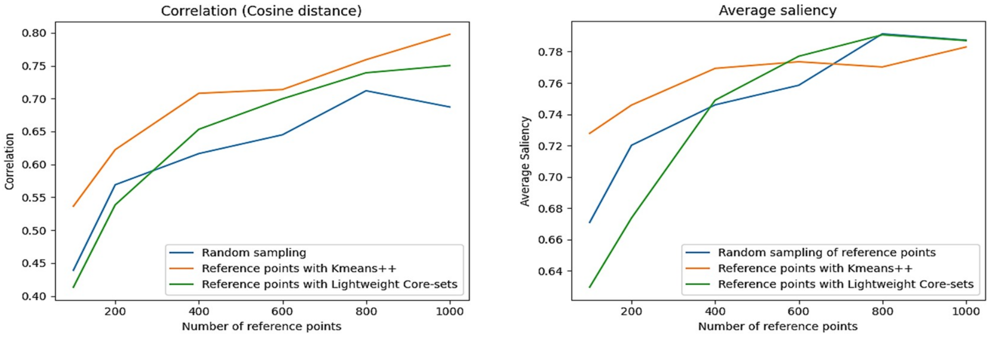


**Figure S24**: **Comparison of reference point selection methods on visualization metrics.**

### Supplementary Tables

#### The benchmarking datasets

| Image size | Features | Tissue type | Resol. | Npy file size |
|---|---|---|---|---|
| 286×262 | 5255 | Mouse brain hemisphere (hA7ko) | 20µm | 1.5GB |
| 331×248 | 5255 | Mouse brain hemisphere (APPPS1 × hA7ko) | 20µm | 1.7GB |
| 314×294 | 5255 | Mouse brain hemisphere (APPPS1) | 20µm | 1.9GB |
| 314×285 | 5255 | Mouse brain hemisphere (C57BL/6J wild-type) | 20µm | 1.8GB |

**Table S1**: **Mouse datasets.** Dataset characteristics for the mouse brain benchmarking dataset (PXD056609). The mouse models were recently described in (*33*).

| Link | Image size | Features | Uploader | Tissue type | Resol. | Npy file size |
|---|---|---|---|---|---|---|
| Dataset 1 | 62×160 | 4505 | Manuel Liebeke (MPI Bremen) | Bacterial vesicles | 100µm | 175MB |
| Dataset 2 | 135×199 | 1755 | Schwaiger-Haber et al. (*56*) | Brain with tumor (mouse) | 50µm | 184MB |
| Dataset 3 | 104×108 | 4285 | Tingting Lu | Ovary (mouse) | 20µm | 188MB |
| Dataset 4 | 550×824 | 5255 | Brittney Gorman (PNNL) | Kidney (human) | 5µm | 9.3GB |
| Dataset 5 | 521×417 | 10505 | Lars Gruber (HS Mannheim) | Spinal cord (mouse) | 5µm | 8.9GB |
| Dataset 6 | 772×596 | 16505 | Dušan Veličković (PNNL) | Lung (human) | 20µm | 30GB |

**Table S2**: **METASPACE datasets.** Dataset characteristics for the diverse benchmarking dataset with slides from METASPACE (*9*).
